# Binding Thermodynamics and Kinetics Calculations Using Chemical Host and Guest: A Comprehensive Picture of Molecular Recognition

**DOI:** 10.1101/155275

**Authors:** Zhiye Tang, Chia-en A. Chang

## Abstract

Understanding the fine balance between changes of entropy and enthalpy and the competition between a guest and water molecules in molecular binding is crucial in fundamental studies and practical applications. Experiments provide measurements. However, illustrating the binding/unbinding processes gives a complete picture of molecular recognition not directly available from experiments, and computational methods bridge the gaps. Here, we investigated guest association/dissociation with β-cyclodextrin (β-CD) by using microsecond-timescale molecular dynamics (MD) simulations, post-analysis and numerical calculations. We computed association and dissociation rate constants, enthalpy, and solvent and solute entropy of binding. All the computed values of *k*_*on*_, *k*_*off*_, ΔH, ΔS, and ΔG using GAFF-CD and q4MD-CD force fields for β-CD could be compared with experimental data directly and agreed reasonably with experiment findings. Both force fields resulted in similar computed ΔG from independently computed kinetics rates, ΔG=-RTln(k_on_ · C° / k _off_), and thermodynamics properties, ΔG=ΔH – TΔS. The water entropy calculations show that entropy gain of desolvating water molecules are a major driving force, and both force fields have the same strength of non-polar attractions between solutes and β-CD as well. Water molecules play a crucial role in guest binding to β-CD. However, collective water/β-CD motions could contribute to different computed *k*_*on*_ and ΔH values by different force fields, mainly because the parameters of β-CD provide different motions of β-CD, hydrogen-bond networks of water molecules in the cavity of free β-CD and the strength of desolvation penalty. As a result, q4MD-CD suggests that guest binding is mostly driven by enthalpy, while GAFF-CD shows that gaining entropy is the major driven force of binding. The study further interprets experiments, deepens our understanding of ligand binding, and suggests strategies for force field parameterization.

## 1. Introduction

Molecular recognition determines the binding of two molecules — a common phenomenon in chemical and biological processes. Thus, understanding molecular recognition is of interest in both fundamental studies and has practical applications in chemical industries and drug discovery. Molecular binding can be characterized by thermodynamic and kinetic properties, which are governed by factors such as polar and non-polar interactions between solutes, solvent effects, molecular diffusion and conformational fluctuations. Molecular binding affinity is a straightforward characterizer of recognition and can be obtained experimentally by enthalpimetry such as calorimetry ^1-2^. However, it is difficult to measure the binding entropy directly from experiments, and it is impossible to measure separate entropy contributions from the solvent and solute. Recent studies have revealed the importance of kinetic properties ^3-5^ and showed that drug efficacy is sometimes correlated with the kinetic properties better than binding affinity ^6-7^. Investigating residence time ^8-10^ was proposed in the last decade and has become a useful criterion in predicting drug efficacy. Efforts have been devoted to studies of kinetic–structure relations ^11-14^. Computational methods, which have the advantage of atomistic resolution, are useful tools to investigate the fundamentals of thermodynamics and kinetics. With the recent breakthrough in computational power, molecular dynamics (MD) is able to simulate up to microsecond or millisecond timescale molecular motions ^15-16^. Therefore, computational methods can be used to sample a larger time scale of dynamics of host–guest systems and take advantage of numerical methods to directly extract the enthalpic and entropic profiles of both the solvent and solute, as well as the kinetics from unbiased long-time-scale MD. New findings observed from chemical host–guest systems with theoretically sound computational methods has advanced our knowledge of molecular recognition and brought new insights into ligand–protein systems.

Cyclodextrins (CDs) are a class of cyclic oligosaccharide compounds that can be obtained by degradation of starch by α-1,4-glucan-glycosyltransferases. Depending on the number of glucopyranose units (D-glucose), these compounds can be classified into 6 units (α-CD), 7 units (β-CD), 8 units (γ-CD) and CDs of more glucopyranose units. In β-CD (Figure 1), the 7 glucopyranose units (D-glucose) enclose a cavity with a diameter of about 6.5 Å. The glucopyranose units bring the center of the cavity a hydrophobic surface of the carbon chains in the glucopyranose, whereas the rims of β-CD consist of hydrophilic hydroxyl groups. Notably, the wide and narrow rims of β-CD are asymmetrical. Because of the structure and size of the cavity, β-CD can host a wide variety of guest molecules via nonpolar or polar attractions. With these properties, β-CD and its derivatives have many applications in many fields, such as the cosmetic industry, pharmaceuticals, catalysis, and the food and agricultural industries ^17-23^. Thus, the experimental thermodynamics and kinetics data are available for a variety of β-CD complexes ^24-30^.

**Figure 1.**
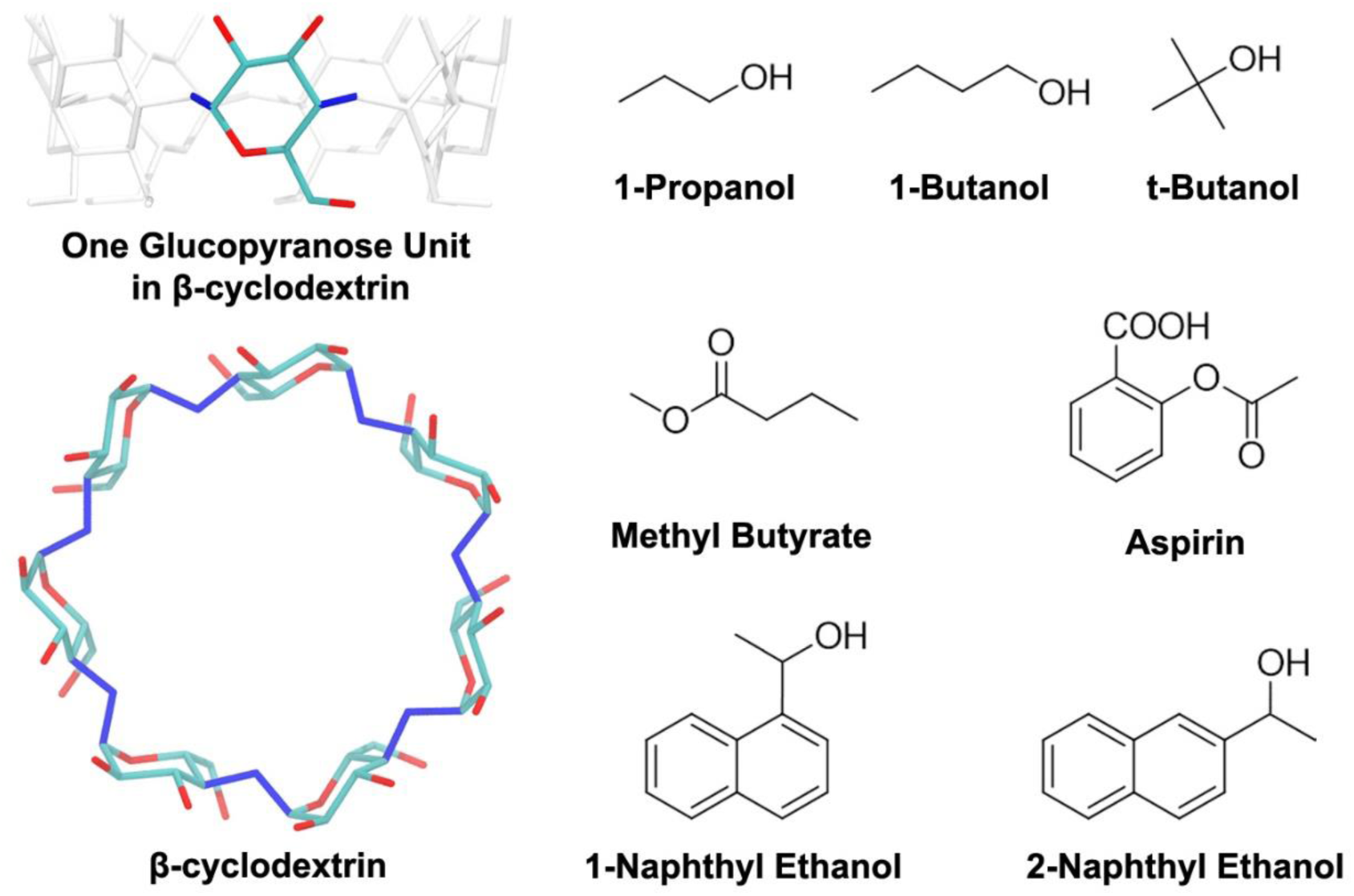
The structure of β-cyclodextrin (β-CD) and the 7 guest molecules. In the structure of β-CD, hydrogen atoms are not shown. For the β-CD internal (vibrational/conformational) entropy calculation, the 14 dihedral angles used to define the conformations of β-CD are in blue.

β-CD consists of 147 atoms, so we can directly investigate the fundamentals of binding thermodynamics and kinetics by MD simulations. Short MD simulations have been used to study ligand binding affinity, dynamics, H-bonds and hydration properties of β-CD complexes ^31-38^. Replica Exchange Molecular Dynamics (REMD) and the Mining Minima 2 method were used to thoroughly sample the bound state of various β-CD complexes and accurately evaluate their binding affinities by using an implicit solvent model ^39-40^. QM/MM methods were also used to investigate the binding pose, binding enthalpy, and properties of the β-CD cavity ^41-44^. In most of these studies, the conformations of β-CD resemble the crystal structures via shorter than 20 ns MD simulations, and no further dynamic information from experiments or long-time-scale MD of β-CD complexes. In addition, although the binding affinity decomposition into enthalpy and entropy has been achieved for some ligands, the use of an implicit solvent model and previous computation techniques cannot obtain both solvent and solute entropy contributions that allow direct comparison with experimental measured ΔH and ΔS. It is still difficult to sample multiple association/dissociation events for kinetic rate calculations ^45^.

The present study applied microsecond-timescale MD simulations with GPU acceleration to compute binding kinetics and thermodynamics of 7 guests with β-CD to reveal the binding mechanisms. Using two force fields for β-CD, GAFF-CD and q4MD-CD, we performed the simulations that sampled multiple association/dissociation events of the complexes to compute *k*_*on*_ and *k*_*off*_, together with MD of free β-CD and guests and a water box to compute entropy and enthalpy from both the contribution of solvent and solutes. The non-polar interactions between solutes and the water entropy gains on binding were the major driving forces of association, regardless of the use of force fields. However, GAFF-CD provided slightly more flexible β-CD, which produced more complicated H-bond networks between the free β-CD and water molecules in the first hydration shell, thereby resulting in slower *k*_*on*_ and larger desolvation penalty. The study also gives details regarding multiple guest association and dissociation pathways and explains how each decomposed energetic and entropic term contributes to binding kinetics/thermodynamics and binding mechanisms.

## 2. Method

### Molecular Systems and Force Fields

We selected 4 weak binders (1-butanol, t-butanol, 1-propanol, methyl butyrate) and 3 strong binders (aspirin, 1-naphthyl ethanol, 2-naphthyl ethanol) as the guest molecules (ligands) for the β-CD host–guest complex. The set of data have similar experimental binding affinity (ΔG) by both kinetics and thermodynamic measurement, and the available rate constants are accessible by MD for sampling multiple association and dissociation events. Structures of guests were manually created by using Vega ZZ ^46^ and the structure of β-CD was obtained from the Cambridge Crystal Data Center (PDB ID: WEWTOJ) (Figure 1). We used two different force fields for β-CD: Amber general force field (GAFF),^47^ termed GAFF-CD, and q4MD-CD force field ^48^. For simulations with GAFF-CD, we took the partial charge of β-CD from a previous work ^40^. For simulations using q4MD-CD, the partial charge of β-CD was provided in the force field. The guests were modeled by only GAFF and the partial charges were computed by using optimization and then ChelpG atomic charge calculation with the Gaussian package ^49^ at 6-31+g(d,p)/B3LYP level. Initial conformations of the complexes were obtained by manually locating the ligand in the center of the cavity of β-CD. All MD simulations and calculations were repeated in the same setting for the free and bound β-CD with GAFF-CD and q4MD-CD.

### Molecular Dynamics Simulations

We performed microsecond-timescale MD runs for a total of 1 to 11 μs simulations (Table SI 1) for each β-CD-ligand system, that is, on the complex, the free β-CD, the free ligand and an empty water box ^50^. For each run, the system was first solvated with exactly 1737 TIP3P water molecules (about 12Å away from the solute) and then optimized to eliminate possible clashes. The water molecules were then equilibrated at 298K for 1 ns, followed by an equilibration of the entire system from 200K to 298K for 150 ps. By using Amber 14 with GPU acceleration ^47, 51^, long production runs were performed at 298K maintained by a Langevin thermostat ^52-53^ in an NPT (isothermal-isobaric) ensemble. A frame was saved every 1 ps for each run. For post-analysis and numerical calculations, we resaved a frame every 10 ps, and the trajectories were visualized by VMD ^54^. Table SI 1 summarizes the total lengths of all MD simulations and numbers of frames used in thermodynamics calculations. We also performed 50-ns MD runs for free β-CD in a vacuum by using GAFF-CD and q4MD-CD to examine the force field parameters, as detailed in SI Section 2.

### Binding Enthalpy Calculation

The binding enthalpy (ΔH) was calculated by ΔH = <E>_Complex_ + <E>_Water_ -<E>_Host_ -<E>_Guest_ ^50^ with the periodicity of the water box, where <E>_Complex_, <E>_Water_, <E>_Host_ and <E>_Guest_ are the averaged potential energy of all molecules from the trajectories of the complex, water, host and guest, respectively. Note that only the bound state conformations from the trajectories of the complexes were used to compute <E>_Complex_. The kinetic energy will be cancelled, so it is not evaluated here. The pressure-volume term is negligible in a solvated system. The computed decomposed enthalpy terms are reported in SI Section 3. We also computed the surface area, number of H-bonds, and number of solvation water (detailed in SI section 4) to explain our decomposed enthalpy terms.

### Solute Entropy Calculation

Solute entropy change (ΔS_Solute_) and water entropy change (ΔS_Water_) are computed separately for binding entropy, ΔS = ΔS_Solute_ + ΔS_Water_, where ΔS_Solute_ was decomposed into host internal (conformational/vibrational) entropy change (ΔS_Host_), guest internal entropy change (ΔS_Guest Int_) and guest external (translational/rotational) entropy change (ΔS_Guest Ext_), illustrated as ΔS_Solute_ = ΔS_Host_ + ΔS_Guest Int_ + ΔS_Guest Ext_.

Notably, the host retains its external degrees of freedom, so there is no change of external entropy for the host. The internal entropy changes of the host (ΔS_Host_ = S_Host Complex_ -S_Host Free_) and guest (ΔS_Guest Int_ = S_Guest Int Complex_ -S_Guest Int Free_) were calculated by taking the difference in internal entropies of their free state and bound state by using the Gibbs entropy formula (eq. 1).

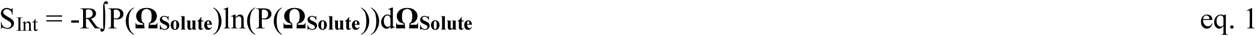

where R is the gas constant and P(**Ω_Solute_**) is the probability distribution of conformations (**Ω_Solute_)** defined by key dihedrals in the species, as detailed in SI Section 5.

In the free states, both β-CD and ligands can diffuse freely in solution, but ligand diffusion may be restricted after binding into the cavity of β-CD. The external entropy loss of the ligand was calculated as ΔS_G__uest Ext_ = S_Guest Ext Complex_ -S_Guest Ext Free_, where S_Guest Ext Free_ = Rln(8π_2_/C_0_) is approximately 6.98 kcal/mol at 298 K ^55-56^. R is the gas constant and C^0^ is the standard concentration (1M). S_Guest Ext Complex_ is the external entropy of the ligands evaluated by numerical integration after aligning the MD trajectories of the complexes to the crystal structure of β-CD,

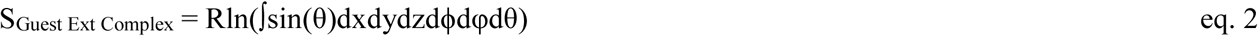

where x, y, z are the Cartesian coordinates of the ligand translation and ϕ, φ, θ are the Euler angles of the ligand rotation. The bin size of x, y, z was 2Å and the bin size of ϕ, φ, θ was 30°. The ϕ and φ were integrated from -π to π and θ was integrated from -π/2 to π/2.

### Water Entropy Calculation

We evaluated the water entropy change on ligand binding (ΔS_Water_) from solvation entropy (ΔS_Solv._) of the free β-CD (ΔS_Solv. Host_), free guests (ΔS_Solv. Guest_) and the complexes (ΔS_Solv. Complex_) by ΔS_Solv._ = ΔS_Solv. Trans_ + ΔS_Solv. Rot_ + ΔS_Solv. Conf_. The ΔS_Solv._ was calculated by the grid cell method ^57-59^. In the grid cell method, the water molecules in the water box are treated as if they are vibrating in a local cell and confined by the forces from surrounding water molecules and solutes. In the cell method, ΔS_Solv_ decomposes. into vibrational (ΔS_Solv. Vib_) and conformational (ΔS_Solv. Conf_) entropies, and ΔS_Solv. Vib_ further decomposes into translational (ΔS_Solv. Trans_) and rotational (ΔS_Solv. Rot_) entropies. The entropy terms were calculated by using eqs. 6.8 to 6.11 in SI, and an in-house script was developed to perform the calculations. The changes in water entropy on ligand binding were evaluated by ΔS_Water_ = ΔS_Solv. Complex_ – ΔS_Solv. Host_ – ΔS_Solv. Guest_. The translational (ΔS_Water Trans_), rotational (ΔS_Water Rot_) and conformational (ΔS_Water Conf_) terms of changes of water entropy on ligand binding were evaluated by the same equation. The setting of solvation water entropy calculation is provided in SI Section 6. We counted the number of solvation water molecules in the free guest, free host, and complexes to explain the water entropy calculations, as detailed in SI section 7.

### Calculations of the Association and Dissociation Rate Constants

As mentioned previously, our MD simulations can sample multiple association and dissociation events in one long run for each system. We define the bound state for a complex when the center of mass of the ligand is within 7.5 Å from the center of mass of β-CD. Any bound/unbound period that lasted longer than 1.0 ns was counted as one bound/unbound event. Because molecules fluctuate during ligand binding, if a ligand stayed or left β-CD for less than 1.0 ns, we did not consider the motion as a bound/unbound event. The average bound/unbound period lengths were calculated and plugged into eqs. 3 and 4 to compute k_on_ and k_off_, where *[solute]* is the concentration of solute estimated by the averaged size of the water box, as detailed in SI Section 8. After obtaining k_on_ and k_off_, we calculated the equilibrium constant K_eq_=k_on_ / k_off_. We also estimated the diffusion-controlled association rate constants by using diffusion coefficients and radii of host and guest molecules for comparison, as detailed in SI section 9.

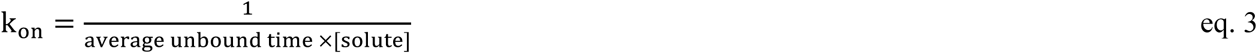

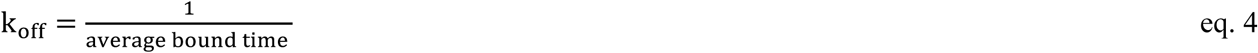

### Binding Free Energy Calculation

The binding free energies can be computed by using the classical thermodynamics and statistical mechanics equations. We evaluated binding free energies (ΔG_Comp1_) from computed binding enthalpy and binding entropy, as in eq. 5. We also evaluated binding free energies (ΔG_Comp2_) from computed association and dissociation rate constants (k_on_ and k_off_), as in eq. 6.

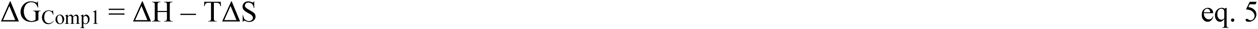

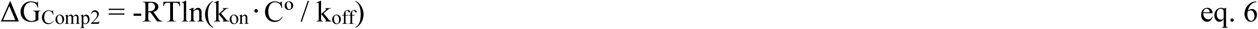

### Uncertainty Evaluation of Computed Properties

We evaluated the uncertainties of binding enthalpy (ΔH), the entropy change of water (ΔS_Water_), internal entropy change of β-cyclodextrin (ΔS_Host_), internal (ΔS_Guest Int_) and external (ΔS_Guest Ext_) entropy change of guests, total binding entropy (ΔS), binding free energy from eq. 5 (ΔG_Comp1_), association rate constant (k_on_), dissociation rate constant (k_off_), and equilibrium constant (K_eq_). The details are provided in SI section 10.

## 3. Results

We used microsecond-timescale MD simulations with an explicit solvent model to compute the binding enthalpies, entropies, and association/dissociation rate constants for β-CD with 7 guest molecules (Figure 1). Binding free energy ΔG was computed by using eq. 5 (ΔG_Comp1_), from the thermodynamic properties and eq. 6 (ΔG_Comp2_), from the kinetics binding rate constants. All computed results yielded reasonable agreement with experimental data (Tables 1 and 2, and Figure 2). Two force fields, GAFF-CD and q4MD-CD, were used to assign parameters for β-CD, and all ligands used the GAFF force field. In general, ΔG computed with GAFF-CD is slightly more positive than experimental data, and binding is driving by entropy. In contrast, q4MD-CD yields slightly more negative computed ΔG than experimental data, and binding is driving by enthalpy. Here we were able to separate enthalpy and entropy calculations, and we therefore use directly computed ΔH and -TΔS to investigate their contributions of a guest and β-CD binding.

**Figure 2.**
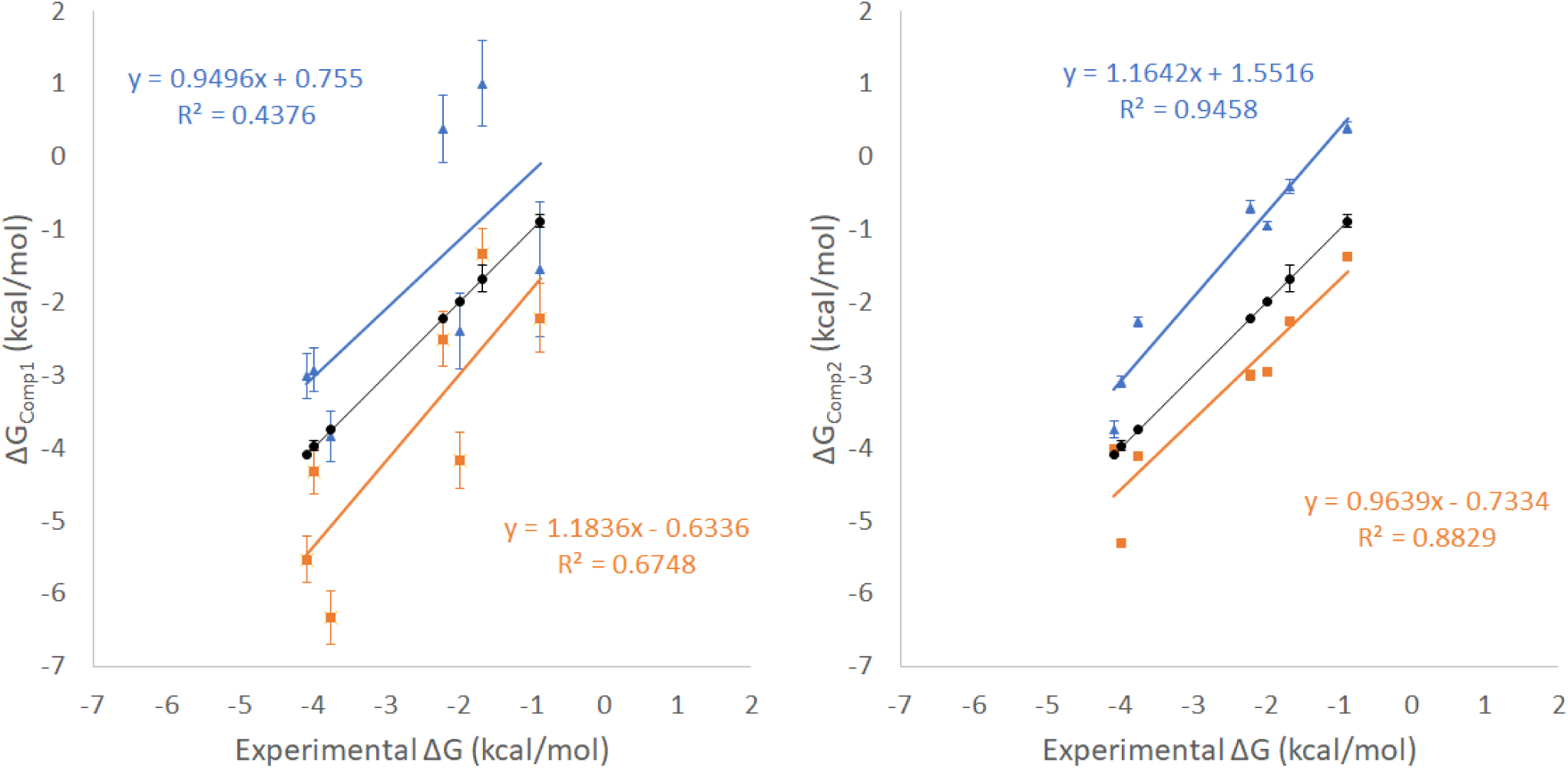
The correlations between ΔG_Comp1_, ΔG_Comp2_ and experimental values. The results from GAFF-CD are shown in blue triangles, and results from q4MD-CD are shown in orange rectangles. The correlations are label correspondingly.

### Binding Enthalpy and Entropy Calculations

The calculated ΔH, -TΔS and ΔG_Comp1_ by using eq. 5 with the GAFF-CD and q4MD-CD force fields are compared with the experimental data in Table 1. The computed ΔG is mostly within 2 kcal/mol of experiments, and they provide a correct trend. Although results from the two force fields reasonably agree with experiment results, GAFF-CD generally underestimated and q4MD-CD overestimated the binding free energy. The intermolecular van der Waals (vdW) attraction between the β-CD and guest is one major driving force for the complex formation (Tables 3 and 4). The water entropy gain on binding is also a major driving force for the complex formation (Table 5). Interestingly, compared to q4MD-CD, the less negative binding affinities using GAFF-CD are primarily from larger desolvation penalty, resulting in the less negative binding enthalpy. The host became more flexible and gained configuration entropy on binding, which is another favorable factor in guest binding (Table 5). In contrast, as compared with GAFF-CD, q4MD-CD modeled a significantly smaller desolvation penalty and more negative ΔH, which become the main driving force for binding. However, the systems need to pay a higher cost in entropy (–TΔS) because the host became more rigid in the bound state.

**Table 1.** The binding free energies (ΔG), enthalpies (ΔH) and entropies (-TΔS) of β-cyclodextrin (β-CD) with 7 guests from experiments and calculations by using GAFF-CD and q4MD-CD force fields. Unavailable experimental values are marked by /. All values are in kcal/mol.

| Calculated (GAFF-CD) |  |  |  |
| --- | --- | --- | --- |
| Guest | $\Delta G_{\text{Comp1}}$ | $\Delta H$ | -T$\Delta S$ |
| 1-Propanol | -1.54±0.93 | 0.54±0.74 | -2.08±0.19 |
| 1-Butanol | 1.01±0.59 | 2.99±0.52 | -1.98±0.06 |
| Methyl Butyrate | -2.40±0.52 | 1.88±0.47 | -4.29±0.05 |
| t-Butanol | 0.38±0.46 | 2.86±0.40 | -2.48±0.06 |
| 1-Naphthyl Ethanol | -3.01±0.31 | -1.41±0.28 | -1.60±0.03 |
| Aspirin | -3.84±0.35 | -0.01±0.31 | -3.83±0.04 |
| 2-Naphthyl Ethanol | -2.93±0.30 | -0.88±0.27 | -2.05±0.03 |
| Calculated (q4MD-CD) |  |  |  |
| Guest | $\Delta G_{\text{Comp1}}$ | $\Delta H$ | -T$\Delta S$ |
| 1-Propanol | -2.21±0.47 | -0.66±0.41 | -1.55±0.06 |
| 1-Butanol | -1.33±0.34 | -0.85±0.30 | -0.48±0.04 |
| Methyl Butyrate | -4.17±0.38 | -2.00±0.33 | -2.17±0.05 |
| t-Butanol | -2.50±0.38 | -2.87±0.33 | 0.37±0.05 |
| 1-Naphthyl Ethanol | -5.53±0.32 | -5.44±0.29 | -0.09±0.04 |
| Aspirin | -6.33±0.37 | -4.30±0.33 | -2.03±0.05 |
| 2-Naphthyl Ethanol | -4.32±0.31 | -4.03±0.28 | -0.29±0.03 |
| Experimental |  |  |  |
| Guest | $\Delta G$ | $\Delta H$ | -T$\Delta S$ |
| 1-Propanol <sup>24</sup> | -0.88±0.09 | 1.43±0.48 | -2.34±0.57 |
| 1-Butanol <sup>24</sup> | -1.67±0.19 | 0.69 | -2.34 |
| Methyl Butyrate <sup>61</sup> | -1.99±0.02 | / | / |
| t-Butanol <sup>90</sup> | -2.22±0.01 | / | / |
| 1-Naphthyl Ethanol <sup>29</sup> | -4.08±0.01 | / | / |
| Aspirin <sup>28</sup> | -3.74±0.00 | / | / |
| 2-Naphthyl Ethanol <sup>29</sup> | -3.97±0.07 | / | / |
| Units: kcal/mol |  |  |  |

**Table 2.** The association and dissociation rate constants of the 7 β-CD complexes from experiments and calculations by using GAFF-CD and q4MD-CD force fields. The standard deviations of rate constants of β-CD-2-naphthyl ethanol with q4MD-CD are not available because of lack of adequate binding events.

| Calculated (GAFF-CD) |  |  |  |  |
| --- | --- | --- | --- | --- |
| Guest | $k_{on}$ (1/s·M) | $k_{off}$ (1/s) | $K_{eq}$ (1/M) | $\Delta G_{Comp2}$ (kcal/mol) |
| 1-Propanol | $2.0 \pm 0.1 \times 10^8$ | $3.9 \pm 0.3 \times 10^8$ | $0.5 \pm 0.06$ | $0.40 \pm 0.08$ |
| 1-Butanol | $2.2 \pm 0.1 \times 10^8$ | $1.1 \pm 0.2 \times 10^8$ | $2.0 \pm 0.3$ | $-0.41 \pm 0.10$ |
| Methyl Butyrate | $3.9 \pm 0.1 \times 10^8$ | $8.0 \pm 0.6 \times 10^7$ | $4.9 \pm 0.4$ | $-0.94 \pm 0.05$ |
| t-Butanol | $2.3 \pm 0.1 \times 10^8$ | $8.5 \pm 1.2 \times 10^7$ | $3.2 \pm 0.4$ | $-0.69 \pm 0.08$ |
| 1-Naphthyl Ethanol | $8.6 \pm 1.1 \times 10^8$ | $1.5 \pm 1.9 \times 10^6$ | $558.9 \pm 100$ | $-3.75 \pm 0.12$ |
| Aspirin | $1.1 \pm 0.01 \times 10^9$ | $2.4 \pm 0.3 \times 10^7$ | $46 \pm 4.3$ | $-2.27 \pm 0.06$ |
| 2-Naphthyl Ethanol | $9.5 \pm 0.4 \times 10^8$ | $5.2 \pm 0.9 \times 10^6$ | $182 \pm 23$ | $-3.08 \pm 0.08$ |
| Calculated (q4MD-CD) |  |  |  |  |
| Guest | $k_{on}$ (1/s·M) | $k_{off}$ (1/s) | $K_{eq}$ (1/M) | $\Delta G_{Comp2}$ (kcal/mol) |
| 1-Propanol | $1.2 \pm 0.02 \times 10^9$ | $1.2 \pm 0.02 \times 10^8$ | $10.1 \pm 0.2$ | $-1.37 \pm 0.01$ |
| 1-Butanol | $1.5 \pm 0.03 \times 10^9$ | $3.3 \pm 0.07 \times 10^7$ | $45.9 \pm 1.2$ | $-2.27 \pm 0.02$ |
| Methyl Butyrate | $1.6 \pm 0.1 \times 10^9$ | $1.1 \pm 0.1 \times 10^7$ | $146.8 \pm 15$ | $-2.95 \pm 0.06$ |
| t-Butanol | $1.1 \pm 0.07 \times 10^9$ | $7.2 \pm 1.0 \times 10^6$ | $157.5 \pm 18$ | $-3.00 \pm 0.07$ |
| 1-Naphthyl Ethanol | $1.2 \pm 0.5 \times 10^9$ | $1.4 \pm 0.5 \times 10^6$ | $876.0 \pm 42$ | $-4.01 \pm 0.03$ |
| Aspirin | $3.2 \pm 0.3 \times 10^9$ | $3.1 \pm 0.9 \times 10^6$ | $1035.3 \pm 83$ | $-4.11 \pm 0.05$ |
| 2-Naphthyl Ethanol | $3.91 \times 10^9$ | $5.0 \times 10^5$ | $7788.3$ | $-5.31$ |
| Experimental |  |  |  |  |
| Guest | $k_{on}$ (1/s·M) | $k_{off}$ (1/s) | $K_{eq}$ (1/M) | $\Delta G$ (kcal/mol) |
| 1-Propanol <sup>90</sup> | $5.1 \pm 0.7 \times 10^8$ | $1.2 \pm 0.07 \times 10^8$ | $4.2 \pm 0.6$ | $-0.88 \pm 0.09$ |
| 1-Butanol <sup>90</sup> | $2.8 \pm 0.8 \times 10^8$ | $3.8 \pm 0.6 \times 10^7$ | $7.2 \pm 2.0$ | $-1.67 \pm 0.19$ |
| Methyl Butyrate <sup>61</sup> | $3.7 \pm 0.3 \times 10^8$ | $1.28 \pm 0.03 \times 10^7$ | $29 \pm 1$ | $-1.99 \pm 0.02$ |
| t-Butanol <sup>90</sup> | $3.6 \pm 0.1 \times 10^8$ | $0.85 \pm 0.01 \times 10^7$ | $42.6 \pm 1.0$ | $-2.22 \pm 0.01$ |
| 1-Naphthyl Ethanol <sup>29</sup> | $4.7 \pm 1.9 \times 10^8$ | $4.8 \pm 1.8 \times 10^5$ | $979 \pm 10$ | $-4.08 \pm 0.01$ |
| Aspirin <sup>28</sup> | $7.2 \pm 0.04 \times 10^8$ | $1.3 \pm 0.03 \times 10^6$ | $549 \pm 2$ | $-3.74 \pm 0.00$ |
| 2-Naphthyl Ethanol <sup>29</sup> | $2.9 \pm 1.6 \times 10^8$ | $1.8 \pm 0.7 \times 10^5$ | $820 \pm 90$ | $-3.97 \pm 0.07$ |

Unlike for experiments and most calculations that measured ΔG and ΔH, we independently computed ΔH and –TΔS. Nevertheless, we still observed entropy-enthalpy compensation in our systems with different force fields, so the compensation in these systems indeed has a physical implication and is not the artifact from mathematics of eq. 5. We compared the calculated ΔH and -TΔS for 1-propanol and 1-butanol for which experimental data are available. Both experiments and calculations with GAFF-CD showed a small positive ΔH, which slightly opposed binding, but the systems gained more entropy to compensate losing enthalpy on binding. Although only 2 guests studied here have experimental ΔH and -TΔS, the binding data for other alcohols of β-CD with calorimetry (ITC) and UV experiments commonly show gaining entropy, and enthalpy was not the predominant determinant for binding in all cases (Figure SI 7). The following subsections present results of binding enthalpy of solute-solute and solute-solvent and binding entropy of solutes and solvent.

### Changes of Enthalpy from Different Components

We evaluated the absolute values of binding enthalpy by using potential energies <E> from microsecond-timescale MD trajectories of the four species, and the convergences of the enthalpy calculations were first examined (Figures SI 8 to 10). For all simulations, the potential energy reaches a stable value within a fluctuation of 0.4 kcal/mol, which is also within reported experimental uncertainty ^24, 28-29, 60-61^. The fluctuations of potential energies of β-CD-1–propanol, β-CD-1–butanol and β-CD-t–butanol complexes are larger with GAFF-CD than the other systems. The fractions of the bound states of these systems are lower than with the other systems; thus, the smaller numbers of sampling result in larger fluctuations.

To understand binding enthalpies, we decomposed the calculated values into various contributions (Table 3). We also provide the decompositions into vdW and Coulombic interactions for ΔH_Solute Inter_, ΔH_Host Conf_, ΔH_Host-Water_ and ΔH_Guest-Water_ (Tables 4, 6, 7). Regardless of type of force field used, we immediately noticed that ΔH_Host-Guest_ data are all favorable (negative), and stronger binders such as naphthyl ethanol and aspirin have large negative ΔH_Host-Guest_ value, ∼ -30 kcal/mol (Table 3). However, ΔH_Desolvation_ largely compensates the solute-solute attraction, for significantly smaller ΔH, ranging from 3.0 to -1.4 kcal/mol with GAFF-CD and -0.9 to -5.7 kcal/mol with q4MD-CD. The computed ΔH with GAFF-CD yielded positive values for weak binders (ΔG > -2.0 kcal/mol), which has been seen in experiments. In contrast, ΔH values are all negative with q4MD-CD. On binding, both β-CD and the guest desolvate water molecules and lose interaction with water molecules; thus, ΔH_Host-Water_ and ΔH_Guest-Water_ values are all positive. The values are larger for stronger binders; presumably, guests such as naphthyl ethanols have larger size as compared with weaker/small binders such as propanol. The water molecules released after binding regain interactions with other water molecules, thereby resulting in all negative ΔH_Water-Water_ values. However, this term is not negative enough to counterbalance the loss from water-solute interactions. As a result, desolvation enthalpy is inevitably all positive and becomes the major force that opposes binding. Of note, the desolvation penalty from the polar contribution is the major force that opposes binding with GAFF-CD, which results from losing more intermolecular H-bonds between water molecules and β-CD (Table 7). The partial charges and the non-polar attractions between guests and β-CD modeled with both force fields are highly similar. However, different parameters for the bonded terms result in different modeled β-CD conformations, which significantly changes the H-bond networks between water molecules and β-CD. As illustrated in Figure 3 and Table SI 2, the conformation results in breaking more intermolecular H-bonding and larger desolvation penalty in GAFF-CD on guest binding.

**Table 3.** The binding enthalpy decompositions of β-CD complexes by using GAFF-CD and q4MD-CD force fields. The total binding enthalpy (ΔH) is decomposed into solute-solute interaction change (ΔH_Host-Guest_) and desolvation energy (ΔH_Desolvation_). ΔH_Host-Guest_ includes conformational enthalpy changes of host (ΔH_Host Conf_) and guest (ΔH_Guest Conf_), and host–guest interaction change (ΔH_Solute Inter_). ΔH_Desolvation_ decomposes into host–water (ΔH_Host-Water_), guest–water (ΔH_Guest-Water_) and water–water (ΔH_Water-Water_) interaction changes. All values are in kcal/mol.

| Calculated (GAFF-CD) |  |  |  |  |  |  |  |  |  |
| --- | --- | --- | --- | --- | --- | --- | --- | --- | --- |
| Guest | $\Delta H$ | $\Delta H_{\text{Host-Guest}}$ | $\Delta H_{\text{Host Conf}}$ | $\Delta H_{\text{Guest Conf}}$ | $\Delta H_{\text{Solute Inter}}$ | $\Delta H_{\text{Desolvation}}$ | $\Delta H_{\text{Host-Water}}$ | $\Delta H_{\text{Guest-Water}}$ | $\Delta H_{\text{Water-Water}}$ |
| 1-Propanol | 0.54 | -10.64 | -0.87 | -0.01 | -9.76 | 11.18 | 12.20 | 6.92 | -7.93 |
| 1-Butanol | 2.99 | -15.28 | -2.25 | 0.00 | -13.03 | 18.27 | 20.55 | 8.34 | -10.62 |
| Methyl Butyrate | 1.88 | -19.31 | -2.85 | 0.02 | -16.48 | 21.19 | 25.58 | 10.28 | -14.67 |
| t-Butanol | 2.86 | -15.71 | -2.55 | -0.04 | -13.13 | 18.57 | 20.71 | 8.39 | -10.53 |
| 1-Naphthyl Ethanol | -1.41 | -30.60 | -4.45 | 0.03 | -26.18 | 29.19 | 35.45 | 14.93 | -21.19 |
| Aspirin | -0.01 | -31.53 | -3.77 | 0.23 | -27.99 | 31.52 | 34.60 | 19.15 | -22.22 |
| 2-Naphthyl Ethanol | -0.88 | -28.45 | -3.97 | 0.13 | -24.61 | 27.57 | 33.21 | 13.76 | -19.40 |
| Calculated (q4MD-CD) |  |  |  |  |  |  |  |  |  |
| Guest | $\Delta H$ | $\Delta H_{\text{Host-Guest}}$ | $\Delta H_{\text{Host Conf}}$ | $\Delta H_{\text{Guest Conf}}$ | $\Delta H_{\text{Solute Inter}}$ | $\Delta H_{\text{Desolvation}}$ | $\Delta H_{\text{Host-Water}}$ | $\Delta H_{\text{Guest-Water}}$ | $\Delta H_{\text{Water-Water}}$ |
| 1-Propanol | -0.66 | -13.61 | -2.05 | 0.04 | -11.60 | 12.95 | 13.72 | 7.74 | -8.50 |
| 1-Butanol | -0.85 | -16.66 | -2.66 | 0.09 | -14.10 | 15.82 | 17.72 | 9.21 | -11.10 |
| Methyl Butyrate | -2.00 | -20.61 | -3.53 | 0.18 | -17.26 | 18.61 | 22.05 | 11.10 | -14.55 |
| t-Butanol | -2.87 | -17.38 | -3.50 | -0.06 | -13.82 | 14.51 | 16.31 | 7.87 | -9.67 |
| 1-Naphthyl Ethanol | -5.44 | -28.56 | -3.42 | 0.20 | -25.34 | 23.12 | 29.65 | 14.51 | -21.04 |
| Aspirin | -4.30 | -29.85 | -3.39 | 0.57 | -27.03 | 25.55 | 28.91 | 18.17 | -21.53 |
| 2-Naphthyl Ethanol | -4.03 | -27.41 | -3.38 | 0.16 | -24.20 | 23.38 | 28.82 | 13.95 | -19.39 |
| Units: kcal/mol |  |  |  |  |  |  |  |  |  |

**Figure 3.**
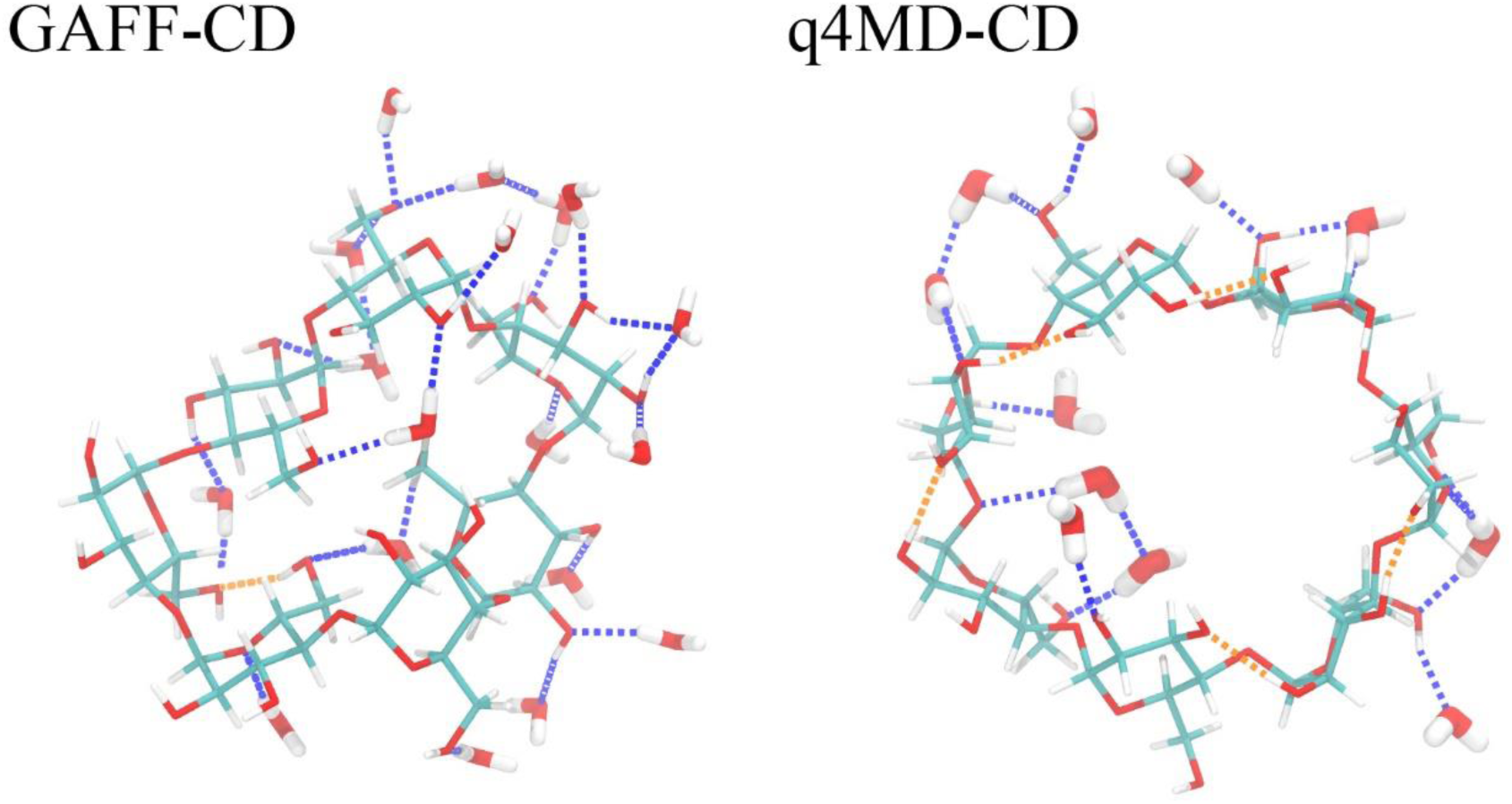
The hydrogen bond (H-bond) patterns of representative free β-CD conformations with GAFF-CD and q4MD-CD. With GAFF-CD, there are 18 water molecules forming H-bonds with β-CD, 24 H-bonds with water (blue dotted lines), and 1 intramolecular H-bond (orange dotted lines). With q4MD-CD, there are 11 water molecules forming H-bonds with β-CD, 16 H-bonds with water, and 5 intramolecular H-bonds.

Strong enough intermolecular attraction ΔH_Solute Inter_ is always required for molecular recognition, and vdW attraction was the major driving force for all the guests (Table 4). The loss of intramolecular H-bonds of β-CD on guest binding is compensated by the intermolecular Columbic attraction in the bound state (ΔH_Host Conf (Coul)_ in Table 6). Although the values of ΔH_Host-Guest_ with both force fields are similar, the decomposition shows significantly larger numbers of each contribution to ΔH_Host Conf_, such as ΔH_Host Conf (vdW)_ and ΔH_Host Conf (Coul)_ of β-CD, with GAFF-CD (Table 6) because of larger conformational changes after ligand binding. The free β-CD prefers flipping 2 glucopyranose units instead of holding an open cavity, as shown in the crystal structure (Figure 4). The glucose rings flipped outward during ligand binding, which lost more water molecules to allow the binding site of β-CD to be accessible for guests to bind. In contrast, q4MD-CD appeared to have crystal-like host structures in both the free and bound states. Note that in vacuum, both GAFF-CD and q4MD-CD sampled predominantly crystal-like host structures (Figure SI 1), which indicates that the glucose ring flipping is largely induced by the hydration shell. GAFF-CD not only changed host conformations upon ligand binding but also makes the host more flexible.

**Table 4.**
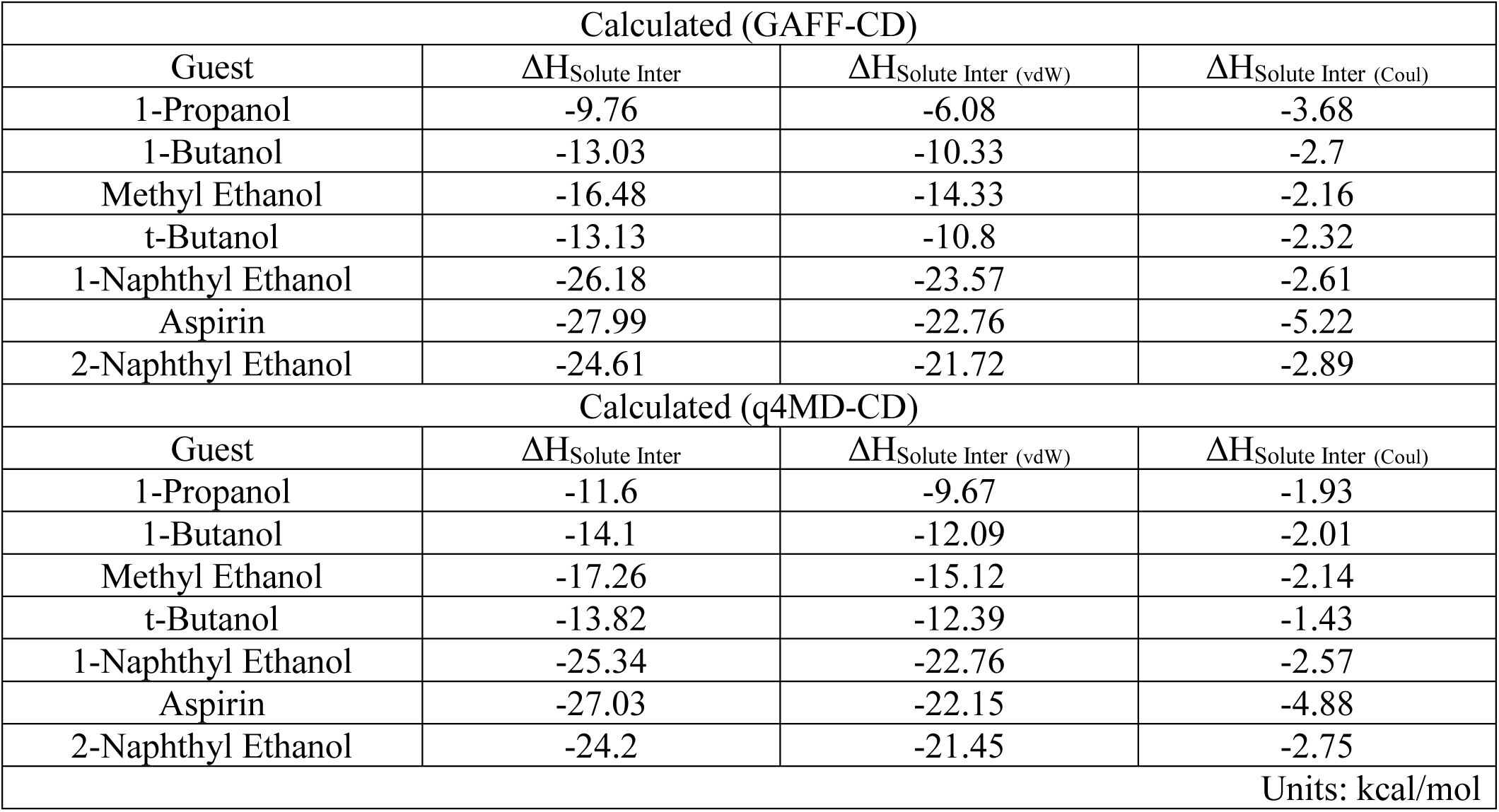
Decompositions of host-guest interaction change (ΔH_Solute Inter_) into van der Waals and Coulombic energies by using GAFF-CD and q4MD-CD force fields. ΔH_Solute Inter_ is decomposed into van der Waals energy (ΔH_Solute Inter (vdW)_), and Coulombic energy (ΔH_Solute Inter (Coul)_). All values are in kcal/mol.

**Figure 4.**
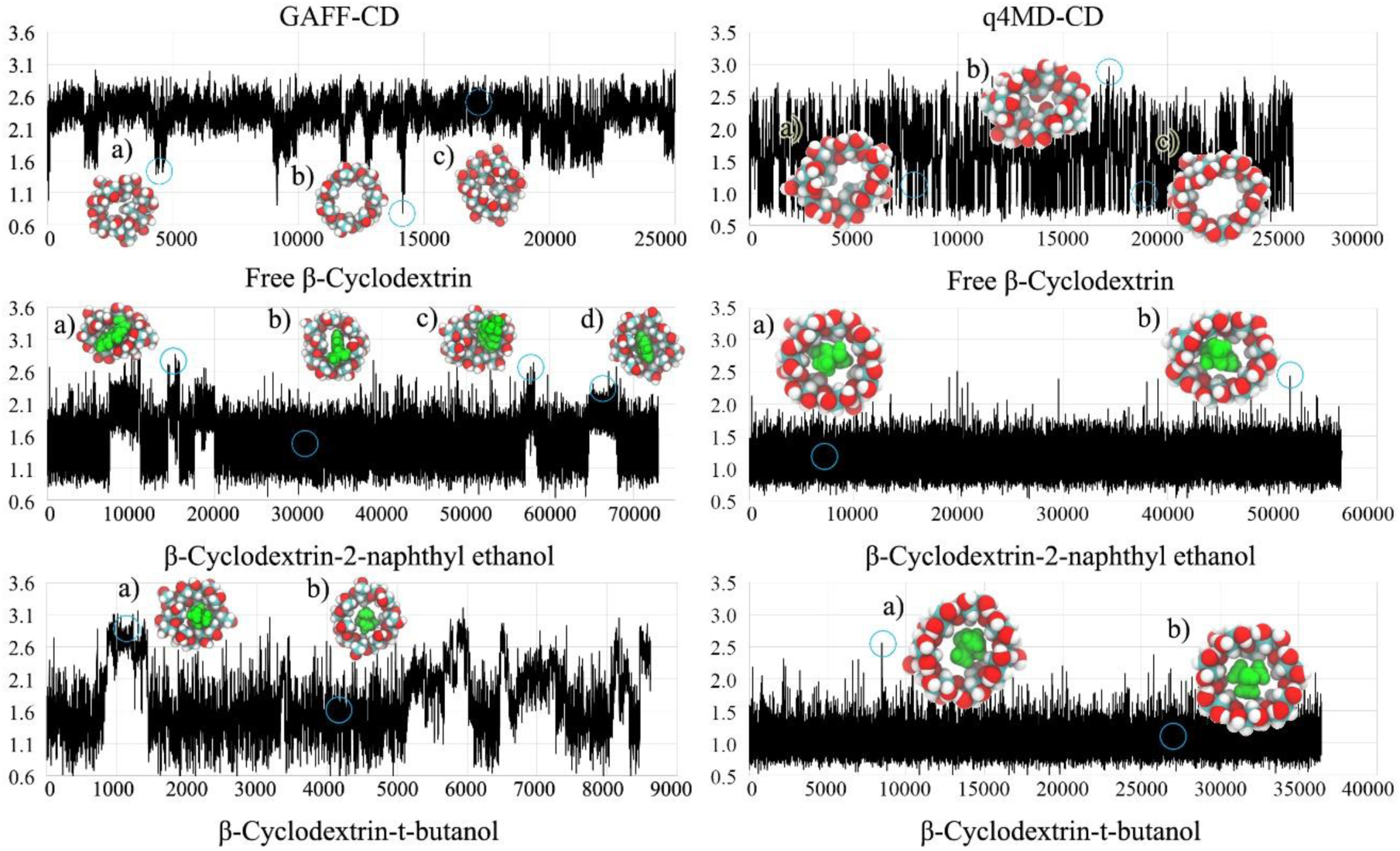
RMSD plots and representative conformations of β-CD for free β-CD, β-CD-2-naphthyl ethanol complex and β-CD-t-butanol complex with GAFF-CD and q4MD-CD. RMSD (Å) are computed against the crystal structure by using conformations chosen every 100 ps from all conformations of free β-CD and bound-state conformations of complexes. Representative conformations are shown near the labels and circles on the plots. In the representative conformations, ligands are in green.

### Changes of Solute Entropy on Ligand Binding

Solute entropy, also termed configuration entropy, reflects the flexibility of a molecular system. Here we used numerical integration to compute each entropy term, by eq. 1 for internal (conformational/vibrational) and eq. 2 for external (translational/rotational) solute entropy. We used the well-defined dihedral distribution analyzed from our MD trajectories to compute internal solute entropy, as detailed in SI Section 5. For the small and weak binding guests studied here, the internal solute entropy of a guest is nearly identical when the guest is in the free solution or bound to β-CD, suggesting a weak correlation between internal degrees of freedom and external translational and rotational degrees of freedom (Table 5 and SI Movie). The external and internal entropy terms therefore can be computed separately in our systems. The calculated entropy values are shown in Table 5. A system is well known to lose configuration entropy because the intermolecular attractions inevitably rigidify the 2 molecules on binding ^39-40, 62-63^. For example, a drug binding to HIV protease can result in >10 kcal/mol entropic penalty from rigidifying conformational and vibrational degrees of freedom ^64^. Additional loss of the configuration entropy also comes from translational and rotational entropy of a ligand, > 7 kcal/mol, by confining itself in a snug binding site ^55-56^. Post-analyzing our MD trajectories showed that all guests and β-CD were not markedly rigidified in the bound state. The guests lost ∼ 1.5–2 kcal/mol external entropy, and β-CD was slightly more flexible, gaining 0.5–1.8 kcal/mol internal entropy with GAFF-CD or being unchanged with q4MD-CD on guest binding (Table 5). Even though different force fields for the host yield considerable differences in β-CD fluctuations, the variation in internal entropy of guests (-TΔS_Guest Int_) is negligible. Also, the interactions between β-CD and guests are not large enough to strongly confine a guest to a handful of well-defined bound guest conformations, thereby allowing a guest to freely tumble and diffuse in the cavity of β-CD. Figure SI 11 illustrates that ligands can pose a wide collection of locations and orientations covering the center and top regions of the binding site, thereby resulting in a small reduction of -TΔS_Guest Ext_.

**Table 5.**
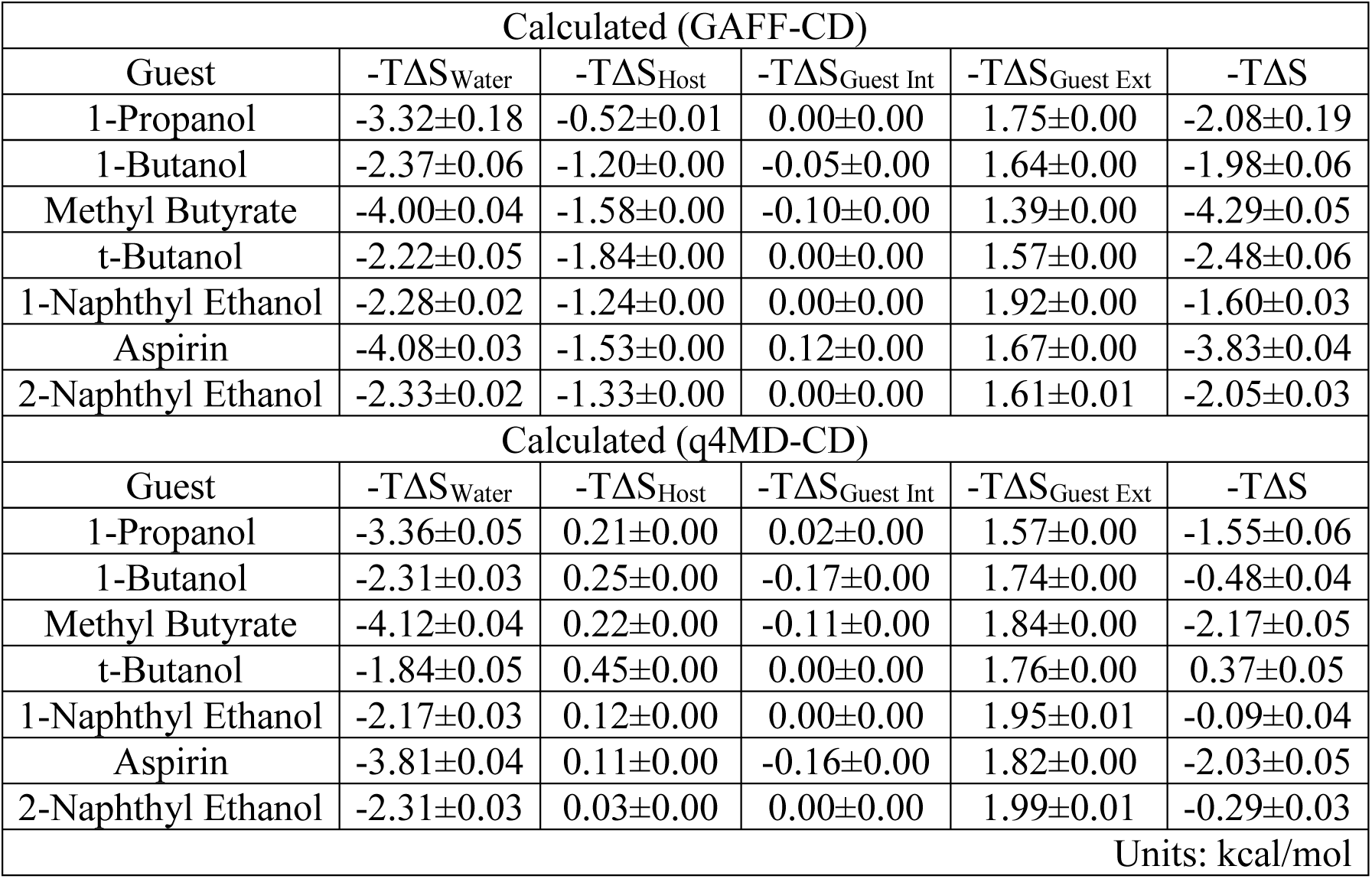
The binding entropy decomposition of β-CD complexes by using GAFF-CD and q4MD-CD force fields. -TΔS_Water_, -TΔS_Host_, -TΔS_Guest Int_, -TΔS_Guest Ext_ and -TΔS are the entropy change of water, internal entropy change of β-cyclodextrin, internal and external entropy change of guests, and the total binding entropy at 298K. All values are in kcal/mol.

**Table 6.** Decompositions of β-CD conformational enthalpy change (ΔH_Host Conf_) into bonded, van der Waals and Coulombic energies by using GAFF-CD and q4MD-CD force fields. ΔH_Host Conf_ is decomposed into bonded energy (ΔH_Host Conf (Bonded)_), van der Waals energy (ΔH_Host Conf (vdW)_), and Coulombic energy (ΔH_Host Conf (Coul)_). All values are in kcal/mol.

| Calculated (GAFF-CD) |  |  |  |  |
| --- | --- | --- | --- | --- |
| Guest | $\Delta H_{\text{Host Conf}}$ | $\Delta H_{\text{Host Conf (Bonded)}}$ | $\Delta H_{\text{Host Conf (vdW)}}$ | $\Delta H_{\text{Host Conf (Coul)}}$ |
| 1-Propanol | -0.87 | 1.37 | 1.76 | -4.00 |
| 1-Butanol | -2.25 | 3.97 | 5.22 | -11.44 |
| Methyl Ethanol | -2.85 | 4.65 | 6.01 | -13.51 |
| t-Butanol | -2.55 | 5.05 | 6.07 | -13.67 |
| 1-Naphthyl Ethanol | -4.45 | 6.23 | 8.71 | -19.39 |
| Aspirin | -3.77 | 5.89 | 7.95 | -17.62 |
| 2-Naphthyl Ethanol | -3.97 | 5.54 | 8.15 | -17.66 |
| Calculated (q4MD-CD) |  |  |  |  |
| Guest | $\Delta H_{\text{Host Conf}}$ | $\Delta H_{\text{Host Conf (Bonded)}}$ | $\Delta H_{\text{Host Conf (vdW)}}$ | $\Delta H_{\text{Host Conf (Coul)}}$ |
| 1-Propanol | -2.05 | -0.38 | 1.79 | -3.46 |
| 1-Butanol | -2.66 | -0.38 | 1.95 | -4.23 |
| Methyl Ethanol | -3.53 | -0.47 | 2.29 | -5.34 |
| t-Butanol | -3.50 | -0.83 | 2.18 | -4.86 |
| 1-Naphthyl Ethanol | -3.42 | 0.62 | 2.57 | -6.62 |
| Aspirin | -3.39 | 0.34 | 2.44 | -6.16 |
| 2-Naphthyl Ethanol | -3.38 | 0.21 | 2.57 | -6.16 |
|  |  |  |  | Units: kcal/mol |

**Table 7.** Decompositions of host-water (ΔH_Host-Water_) and guest-water (ΔH_Guest-Water_) interaction enthalpy changes into van der Waals and Coulombic energies by using GAFF-CD and q4MD-CD force fields. ΔH_Host-Water_ and ΔH_Guest-Water_ decompose into van der Waals energy (ΔH_Host-Water (vdW)_ and ΔH_Guest-Water (vdW)_) and Coulombic energy (ΔH_Host-Water (Coul)_ and ΔH_Guest-Water (Coul)_) terms. All values are in kcal/mol.

| Calculated (GAFF-CD) |  |  |  |  |  |  |
| --- | --- | --- | --- | --- | --- | --- |
| Guest | $\Delta H_{\text{Host-Water}}$ | $\Delta H_{\text{Host-Water}}$<br>(vdW) | $\Delta H_{\text{Host-Water}}$<br>(Coul) | $\Delta H_{\text{Guest-Water}}$ | $\Delta H_{\text{Guest-Water}}$<br>(vdW) | $\Delta H_{\text{Guest-Water}}$<br>(Coul) |
| 1-Propanol | 12.20 | 1.34 | 10.85 | 6.92 | 2.12 | 4.80 |
| 1-Butanol | 20.55 | 0.12 | 20.44 | 8.34 | 4.15 | 4.19 |
| Methyl Ethanol | 25.58 | 0.64 | 24.94 | 10.28 | 5.79 | 4.49 |
| t-Butanol | 20.71 | -1.38 | 22.09 | 8.39 | 4.16 | 4.23 |
| 1-Naphthyl Ethanol | 35.45 | 0.91 | 34.54 | 14.93 | 9.62 | 5.32 |
| Aspirin | 34.60 | 0.40 | 34.20 | 19.15 | 9.13 | 10.02 |
| 2-Naphthyl Ethanol | 33.21 | 1.54 | 31.66 | 13.76 | 9.29 | 4.47 |
| Calculated (q4MD-CD) |  |  |  |  |  |  |
| Guest | $\Delta H_{\text{Host-Water}}$ | $\Delta H_{\text{Host-Water}}$<br>(vdW) | $\Delta H_{\text{Host-Water}}$<br>(Coul) | $\Delta H_{\text{Guest-Water}}$ | $\Delta H_{\text{Guest-Water}}$<br>(vdW) | $\Delta H_{\text{Guest-Water}}$<br>(Coul) |
| 1-Propanol | 13.72 | 4.08 | 9.63 | 7.74 | 3.85 | 3.89 |
| 1-Butanol | 17.72 | 5.45 | 12.27 | 9.21 | 4.99 | 4.22 |
| Methyl Ethanol | 22.05 | 6.51 | 15.54 | 11.10 | 6.34 | 4.77 |
| t-Butanol | 16.31 | 4.75 | 11.56 | 7.87 | 5.12 | 2.75 |
| 1-Naphthyl Ethanol | 29.65 | 8.63 | 21.02 | 14.51 | 9.60 | 4.91 |
| Aspirin | 28.91 | 7.75 | 21.16 | 18.17 | 9.35 | 8.82 |
| 2-Naphthyl Ethanol | 28.82 | 8.95 | 19.87 | 13.95 | 9.37 | 4.59 |
| Units: kcal/mol |  |  |  |  |  |  |

**Table 8.**
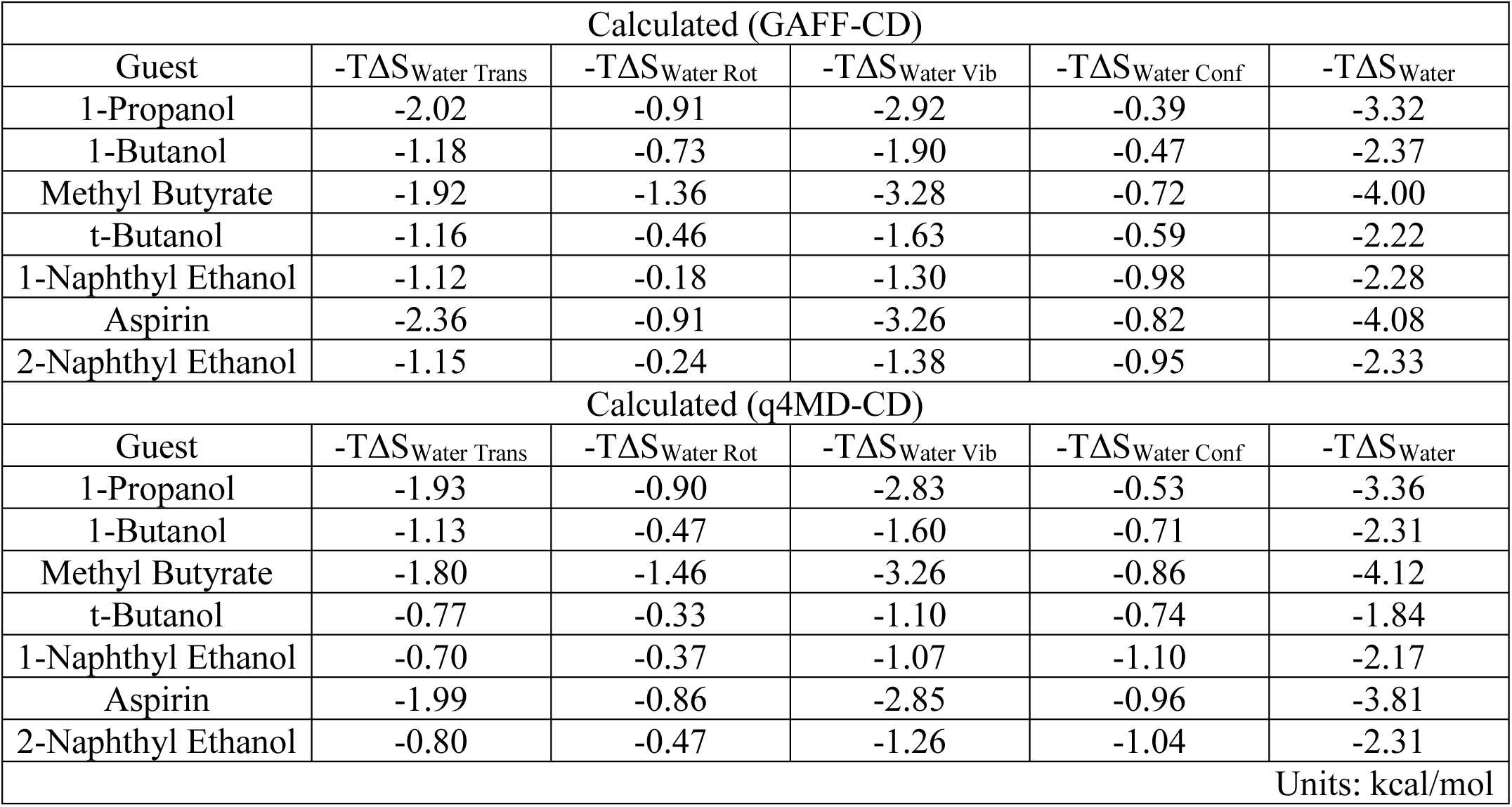
The decompositions of water entropy change by using GAFF-CD and q4MD-CD force fields. The binding water entropy (-TΔS_Water_) decomposes into translational (-TΔS_Water Trans_), rotational (-TΔS_Water Rot_), vibrational (-TΔS_Water Vib_ = -TΔS_Water Trans_ + -TΔS_Water Rot_), and conformational (-TΔS_Water Conf_) terms. All values are in kcal/mol.

### Changes of Water Entropy on Ligand Binding

Water entropy is one major driving force in ligand binding in these systems, contributing to -2 to -4 kcal/mol to the free energy of binding (Tables 1 and 5). Gaining water entropy dominates in the binding of all guests to β-CD with GAFF-CD and the first 3 weaker binders to β-CD with q4MD-CD. The change in water entropy (-TΔS_water_) is a combined effect from rearranging water molecules, which affects their vibrational and conformational entropy, and from releasing the water molecules residing in the cavity of β-CD or interacting with the guest after the complex is formed (Tables 8 and SI 3 to 6). We used a grid cell theory approach to calculate the water entropy over the space near the solute, as compared with the entropy estimated for the bulk solvent. We also used an alternative method that computes the molar entropy of water when different solutes are present to approximate -TΔS_water_ (detailed in SI Section 6). Both methods agree with each other. The theory and implementation were first validated by comparing our computed molar entropy of bulk water at 298K, 5.26 kcal/mol (Tables SI 3 to 5), with standard molar entropy of water at 298K, 4.98kcal/mol. The rigid water TIP3P model performed very well in reproducing standard molar entropy, as reported here and previously, with a two-phase thermodynamic method ^65-66^.

Table 5 lists the changes in water entropy on each guest binding to β-CD, and the contribution from each decomposed entropy term is in Tables SI 3 to 5. Using different force fields to model β-CD did not change the computed -TΔS_Water_, so solute flexibility does not play an important role in water entropy calculations. Aspirin and methyl butyrate have more water entropy gains on binding than do other guests. The 2 guests have more polar functional groups to capture nearby water molecules in their free state, and after forming the complex with β-CD, these water molecules are released, which results in gaining more entropy on binding.

Figure 5 reveals the fluctuation of water molecules presented in the grid space in a pure water box and when β-CD, 2-naphthyl ethanol, or the complex is present using GAFF-CD. The behavior of water molar entropy in q4MD-CD is similar to GAFF-CD, so we only one force field is presented in this figure. The averaged jiggling of water molecules is quantified by computed translational, rotational and conformational entropy, as shown in columns 2-4 of Figure 5, and the darker grid indicates that water molecules in this cell are less mobile. We used an in-house multi-layer visualization program to display the decomposed entropy terms for each grid, and the layer through the center of the solute was selected in the figure. In general, the translational entropy decreases at the surface of the solute because the existence of the solute hinders free diffusion of water molecules. In free β-CD and 2-naphthyl ethanol, the translational entropy of water decreases ∼ 0.2–0.4 kcal/mol around the solutes, as shown by darker cells in the vicinity of each solute in Figure 5B, 5C and 5D.

**Figure 5.**
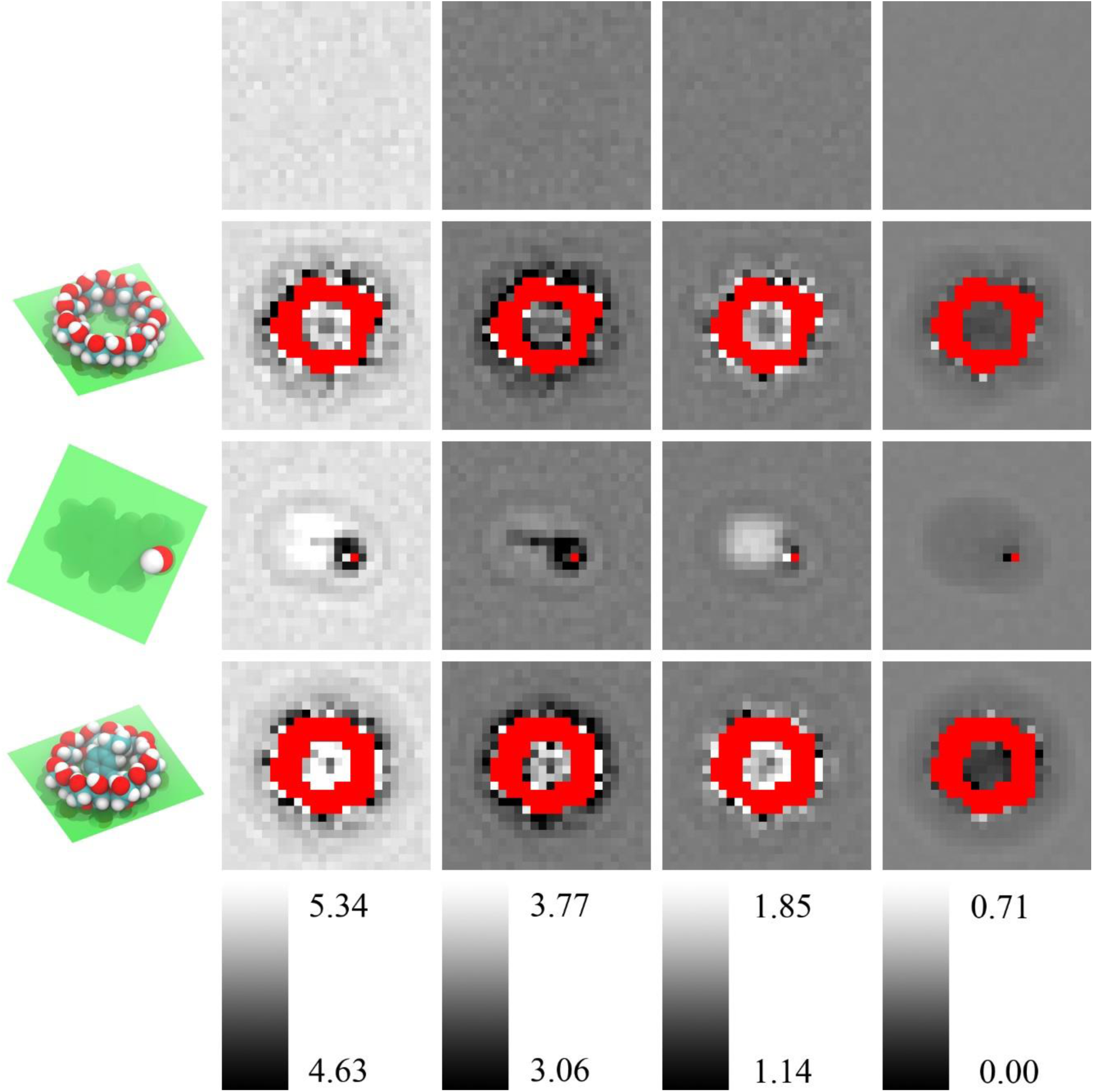
The water entropy decompositions of pure water and in the vicinities of free β-CD, free 2-napthyl ethanol and β-CD-2-napthyl ethanol complex in GAFF-CD. From top to bottom are pure water (A), free β-CD (B), free 2-napthyl ethanol (C), and β-CD-2–naphthyl ethanol complex (D). The plane of the spatial grid in the figures is shown in green in the first column. From left to right are total entropy (1, S_Water_), translational entropy (2, S_Water Trans_), rotational entropy (3, S_Water Rot_) and conformational entropy (4, S_Water Conf_). Each column has a separate color bar (values in kcal/mol). The red areas are regions occupied by the solute molecules, and water entropy cannot be calculated.

The rotational entropy increases on the hydrophobic surface and decreases near hydrophilic regions. For example, with free 2-naphthyl ethanol, the rotational entropy increases ∼ 0.2–0.3 kcal/mol near the carbon chains or benzene ring and in the hydrophobic cavity of free and bound β-CD (see lighter cells in Figures 5B and 5C). The non-polar surface may promote water tumbling, but forming H-bonds with a polar group decreases rotational entropy by ∼ 0.2–0.5 kcal/mol near the hydroxyl group of 2-naphthyl ethanol. The conformational entropy is correlated with the density of water molecules (eq. 6.6 in SI). The cavity of β-CD is large enough to hold ∼ 5–6 water molecules, on average, and the waters slightly decrease their conformational entropy by 0.2-0.3 kcal/mol because the cavity restrains the rearrangement of water molecules. However, the water is not frozen in the cavity and it keeps the fluid-like property, for no significant decrease in translational water entropy. Because the density of water fluctuates during the MD simulations, except for water molecules in the vicinity of solutes, conformational entropy of waters in the bulk solvent or surrounding a solute does not noticeably differ.

### Binding Kinetics: Calculations of Association and Dissociation Rate Constants

The fast kinetics of guest binding to β-CD allowed for directly assessing the association (k_on_) and dissociation (k_off_) rate constants from the bound and unbound lengths during microsecond-long unguided MD simulations. Table 2 summarizes the calculated k_on_ and k_off_ for systems modeled by GAFF-CD and q4MD-CD. The diffusion-controlled limit for the molecular systems is approximated by k_on_diffuse_ = 4πD*R*, where D is the diffusion coefficient obtained by our MD simulations (4×10^-9^ m^2^/s for a guest such as aspirin and 7×10^-10^ m^2^/s for β-CD, regardless of force field used) and R is the sum of the radius of β-CD and a guest (∼ 10Å). The estimated diffusion-controlled association rate constants for all systems are k_on_diffuse_ ∼ 3–4×10^10^ 1/Ms by using the approximated size of each molecule and the diffusion coefficient obtained by MD (Table SI 8). The modeled k_on_ by using GAFF-CD agrees very well with experimental data, and all guests showed 2 orders of magnitude slower k_on_ than k_on_diffuse_. Using q4MD-CD slightly overestimated k_on_ for all guest binding, and the value is one order of magnitude slower than k_on_diffuse_ because of the spatial factor. Because β-CD does not require considerable conformational changes or slow transition to acquire all the guests, experiments revealed no differences in k_on_ for different guests. However, k_on_ modeled with GAFF-CD shows that larger guests such as aspirin and naphthyl ethanols associate marginally faster to β-CD, with k_on_ values close to 10^9^ 1/Ms, as compared with smaller guests such as butanols, with k_on_ ∼ 2–3×10^8^ 1/Ms. The faster association from modeling may result from stronger intermolecular attraction that can more easily retain the guest once 2 molecules collide. In contrast, because k_on_ modeled with q4MD-CD is already fast and close to k_on_diffuse_, a small difference is observed, with k_on_ ∼ 1–4×10^9^ 1/Ms.

The difference in computed k_on_ with the two force fields and from experimental k_on_ result from the intermolecular attractions and the desolvation process. Guest diffusing on the cyclodextrin surface is unlikely due to weak intermolecular interactions, and the β-CD surface features a restricted target area. Assuming no intermolecular attraction, successful binding occurs only with the first collision occurring in the window areas of the cavity (∼ 2.5% of the entire surface), which are the areas covering the top and bottom entrance of the cavity. With q4MD-CD, modeled k_on_ is 20 times slower than k_on_diffuse_, and ∼ 5% of molecular encounters result in successful binding. Therefore, the restricted target area is the main contribution to a slower k_on_. Using the same concept, less than 1% of the initial association results in a stable complex modeled with GAFF-CD, except for aspirin and 2-naphthyl ethanol. The chief reason that causes further slowdown in k_on_ with GAFF-CD is the desolvation process during association. Although the tilt glucose rings in the free β-CD may partly occlude the cavity, rotating the 2 dihedrals in C-O-C for different glucose ring tilting is nearly barrier-less. Replacing water molecules that formed the H-bond network with the free β-CD creates an energy barrier and results in unsuccessful binding, even for a guest already diffused to the cavity. Note, successful binding was considered only when a complex larger than 1 ns was formed in MD simulations.

In contrast to k_on_, k_off_ modeled by q4MD-CD agrees very well with experiments; however, with GAFF-CD, all guests left β-CD approximately one order faster than the measured k_off_ values. Because of the faster k_off_, the equilibrium constants (K_eq_) are systematically smaller than the experimental values, except for 1-naphthyl ethanol. The dissociation rate constants are directly proportional to how long a guest can stay in the pocket of β-CD, also termed residence time in the drug discovery community. Different force field parameters can largely affect k_off_. The longer average bound time indicates more negative ΔH (Table 3), in which ΔH computed with q4MD-CD is stronger than that with GAFF-CD. However, longer bound time does not always require stronger intermolecular attractions, and Table 3 shows that the water effects can be the major differentiating factors. Of note, we sampled hundreds of bound/free states during long simulations (Tables SI 10 and 11) but these are still less than the real experiments.

For the weak binders, we observed one direct association/dissociation pathway in which a guest diffused into the window of the cavity and then stayed with β-CD. The association perturbed the conformations of β-CD to get rid of hydrated waters and flip glucopyranose. We term this pathway the direct binding pathway (SI Movies 1,2,7,8). For the strong binders such as aspirin, for which k_on_ is 3-to 10-fold faster than weak binders modeled by both force fields, we observed one more association/dissociation pathway, termed the sticky binding pathway (SI Movies 3-6, 9-12). The stronger intermolecular attractions allow the guest to stay on the surface of β-CD for surface diffusion to reach the cavity. This situation largely increases the possibility of binding events because the guests can overcome the limitation of a restricted target area of the surface. Note that unlike some ligand–protein binding in which the large biomolecular system needs longer than a microsecond timescale for both molecules to arrange to form a complex, binding processes of guest–β-CD are very fast, in the sub-nanosecond range, without large binding energy barriers. Nevertheless, the intermolecular attractions, possible surface diffusion and desolvation process still play a key role in controlling binding kinetics.

## 4. Discussion

This study demonstrates that unbiased MD simulations can be used to compute CD–guest binding kinetics and thermodynamics. The kinetic properties and binding enthalpies can be extracted directly from the simulations, but post-processing of the MD trajectories by using numerical integration and the grid cell method is needed to compute solute and solvent entropy of binding. Computed values with both force fields, GAFF-CD and q4MD-CD, agree well with experimental measures; however, values strikingly differ depending on whether the binding is driving by enthalpy or entropy.

Binding free energy computed from the kinetic data (ΔG_Comp2_) correlate better with experiment (Figure 2), compared with ΔG_Comp1_ computed from ΔH-TΔS. The calculation only needs one long MD run that includes both solutes in one water box. Eq. 6 may be rewritten as 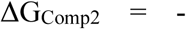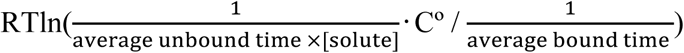. The numbers of bound and unbound events for computing the average time are nearly identical from the same one MD run; thus, another equation to describe binding free energy may be 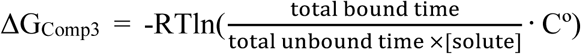. From classical statistical mechanics, the probability of observing a bound complex during this long MD run is *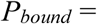*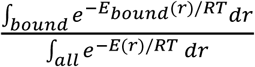. As a result, 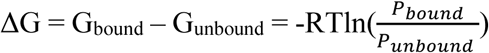 which is the same as equation ΔG_Comp3_. The enthalpy change is also available from the average energy of the bound and unbound states in the trajectories of the complexes, ΔH_Bound-unbound_ = <H_bound_> – <H_unbound_>. The computed enthalpy changes are highly similar to the values computed using four separated MD runs (Figure SI 12 and Table 1). This study aims to investigate the kinetic properties in addition to binding affinities; thus, ΔG_Comp3_ was not used to obtain the results. However, for other weak binding systems that k_on_ and k_off_ measurements are not available, and for the research that focuses on binding free energies, we suggest that one can run a long MD simulation to obtain ΔG and ΔH. It is worth mentioning that although directly computing entropy changes is not needed if ΔG and ΔH are available, the calculation allows researchers to investigate the water effects in a great detail. The calculations also bring further insights into contributions regarding water entropy and solute configuration entropy to binding. However, because configuration entropy calculations inevitably need multi-dimension integral, a larger error than ΔG computed from k_on_ and k_off_ is anticipated.

Intuitively, a more negative binding enthalpy ΔH modeled by q4MD-CD is likely a result of the Lennard-Jones parameters, which could result in more optimized ΔH_Host-Guest 67_. Nevertheless, in the decomposed term ΔH_Solute Inter_ of ΔH_Host-Guest_, the computed ΔH_Solute Inter (vdW)_ and ΔH_Solute Inter (Coul)_ show that both force fields model highly similar inter-solute attractions, and the main determinant is from water. Both force field parameters allow the sugar ring to flip in the free states, and β-CD can easily adjust to an open cavity conformation when forming a complex with a guest. Therefore, unlike existing study showing that substituents attached to decorated β-CDs block a guest from binding ^68^, the ring flipping itself in our study did not hinder guest binding. However, more flipped sugar rings modeled by GAFF-CD allow the formation of more H-bonds between waters and β-CD and a more structured H-bond network as compared with conformations modeled by q4MD-CD. Therefore, the cavity more energetically accommodates stable water molecules, which results in large enthalpy penalty on desolvating those water molecules. The Coulombic term ΔHHost-Water (Coul) becomes the essential energy term that opposes binding due to breakage of β-CD–water H-bonds, and the value is significantly larger with GAFF-CD. We suspected that the bonded parameters with GAFF-CD may highly prefer sugar ring flipping in the free states. However, the MD simulations in vacuum showed that both GAFF-CD and q4MD-CD highly prefer wide-open, crystal structure-like conformations in the free states (Figure SI 1). Because β-CD is reasonably flexible, in particular with GAFF-CD modeling, adding explicit water molecules easily induces the conformation changes. A “slaving” model in which water drives protein fluctuations was proposed ^69^. Recently, a direct measurement of hydration water dynamics in protein systems illustrated that the surface hydration-shell fluctuation drives protein side-chain motions ^70^. Here we showed that the water molecules are highly responsible for molecular recognition in both thermodynamics and kinetics.

Without considering other factors, the theoretical k_on_ may be estimated by multiplying the restricted target area by the diffusion-controlled limit k_on_diffuse_^71^, 2.5% ×4×10^10^ 1/Ms = 10^9^ 1/Ms, which is close to the modeled k_on_ of most guests binding with q4MD-CD. To increase an association rate faster than the theoretical value 10^9^ 1/Ms, when the initially collision did not bring a guest to the restricted target area, intermolecular attractions kept the guest in close proximity to β-CD until the guest reached the cavity. This situation can be observed during MD simulations, as illustrated in the sticky binding pathways with the tighter binder aspirin. Note no long-range electrostatic attractions for rate enhancement because of the neutral molecules. A guest always needs to compete with water molecules during binding. Therefore, k_on_ can be slower than the theoretical value 10^9^ 1/Ms if there exist barriers from removing stable water molecules in the hydration shell. The desolvation barrier increases when the free β-CD forms a larger number of H-bonds between water molecules modeled by GAFF-CD, which results in more unsuccessful guest binding and slower k_on_. As a result, although a less wide-open cavity conformation in free β-CD may seem to directly lead to a slower k_on_ in GAFF-CD, the direct cause is from paying a higher cost to disturb its solvation shell. After forming a complex with a guest, β-CD gained a few kcal/mol, showing a more negative ΔH_Host Conf_ on binding. A similar finding from investigating a binding free energy barriers for a drug binding a protein showed that desolvation of the binding pocket contributes the most to the free energy cost, as opposed to reorganizing the protein binding pocket ^72^. However, for molecular systems that encounter large-scale conformational changes and/or induce fit during ligand binding, rearranging conformations may still significantly affect the association rate constants ^73^, and the kinetic property can be highly system-dependent. With q4MD-CD, because water molecules form a less stable H-bond network, the role of desolvation in binding kinetics is not as important as with GAFF-CD.

Unlike desolvation effects, which are quite different from the 2 force fields, another dominant but similar driving force for binding from both force fields is the attractive component of the vdW energy, ranging from -6 to -23 kcal/mol. This driving force may be expected because of the nonpolar property of the β-CD cavity and the neutral guests. This term mainly accounts for dispersion forces between β-CD and guests in the force-field parameters and is similar for both force fields, with a trend that larger guests have more negative ΔH_Solute Inter (vdW)_. In experiments, measuring the separate contributions for binding from dispersive interactions and classical hydrophobic effects in aqueous environment is challenging, and the absolute values from the dispersive interactions are not available experimentally ^74-75^. As compared with the vdW attraction, the Coulombic attraction between β-CD and guests is significantly weaker because the guests are not highly charged molecules and few intermolecular H-bonds are formed in the complex. The intermolecular attractions are balanced by the enthalpy penalty from disrupting attractions between waters and solutes and gaining water-water enthalpy for those water molecules replaced by a guest, which results in merely a few kcal/mol net enthalpy changes on binding. One may consider hydrophobic effects as the major contributions to β-CD and guest recognition ^76^. Of note, although the pocket of β-CD is non-polar, it is a very tiny cavity, and the rims of β-CD consist of several hydroxyl groups. On binding, ∼ 20–25 water molecules were replaced by a larger guest, which agrees with experimental measurement ^77^ (Table SI 6). However, not all replaced water molecules are “unhappy”. Therefore, although the replaced water molecules also regain water-water attractions in the bulk solvent, the costs to replace the stable water molecules result in the large desolvation penalty.

The systems did not encounter large solute entropy loss, which contrasts with several existing publications that suggested loss of configuration entropy when a drug binds its target protein ^63-64, 78-80^. Unlike most drug-like compounds, which fit tightly to their target protein pocket, our guests only loosely fit in the cavity of β-CD. Therefore, the mobility of β-CD is not reduced considerably by a guest. With GAFF-CD, the hydrated water molecules in the cavity of free β-CD showed an ordered H-bond network and slaved the conformational motions of β-CD. On ligand binding, a bound guest did not form a stable H-bond network with β-CD; thus, β-CD showed a slightly increased flexibility. The guests were also able to form various contacts with β-CD. Similar to alternative contacts provided by the hydrophobic binding pocket of protein systems ^81^, we did not observe rigidity of β-CD with GAFF-CD.

The enthalpy and entropy balance may follow immediately from eq. 5, ΔG_comp1_=ΔH-TΔS ^62^, ^82^-^83^. Therefore, it has been suggested that the entropy-enthalpy compensation is from a much smaller range of experimentally measured ΔG for a series of ligands than the range of ΔH. Different from most experimental techniques, we computed the entropy and enthalpy terms separately, and still observed the entropy-enthalpy compensation. Our guests were all weak binders and did not have a wide spectrum of ΔG. The computed range of ΔH is in a similar ballpark as ΔG, and the range of -TΔS is relatively smaller than ΔG and ΔH. As a result, the enthalpy change mostly governs if a guest is a stronger or weaker binder. Our calculations reveal the physical basis of larger range of ΔH and more similar –TΔS. The enthalpy calculations are based on energy functions in the force fields, but the Gibb’s entropy formula is based on the distribution of the microstates (eq. 1). Unlike protein systems with numerous rotatable bonds and a larger binding site to mostly enclose a ligand, a guest is not completely confined within the cavity of β-CD, and the host remains highly flexible. Interestingly, -TΔS_water_ is similar in both force fields, and -TΔS_water_ is not simply proportional to the size of a guest. Instead, -TΔS_water_ relates more to the hydrophilicity of a guest such as 1-propanol, methyl butyrate and aspirin when forming a complex with β-CD. The free guests reduce more entropy of water in their solvation shell, and these solvation waters gain more entropy on guest binding. For q4MD-CD, because the free β-CD generally has a more open cavity, more waters were released on binding (Table SI 6). However, the ring flipping conformation modelled by GAFF-CD produces a more structured H-bond network for the first hydration shell. As a result, although fewer water molecules were released on binding, those waters gained more entropy than those of β-CD with q4MD-CD, which resulted in a similar computed -TΔS_water_ from both force fields. The results suggest that as in enthalpy, entropy calculations feature a fine balance.

Force-field parameters are critical for accurate modeling and successful prediction ^84-87^. In this study, we used GAFF for β-CD (GAFF-CD) and for all guests, and q4md-CD, a specialized force field for CDs that combines Amber99SB and GLYCAM04 to match experimental geometries from crystal structures and NMR ^48^. It is common practice to seek agreement between the calculated and experimental binding affinities/binding free energies for validating and improving the parameters of force fields or solvent models. Using computed thermodynamics and kinetics (eqs. 5 and 6), both force fields for β-CD showed reasonable agreement with experimental binding affinities, which validated the parameters used. Interestingly, different force fields concluded different driving forces, with GAFF-CD and q4MD-CD showing an entropy-and enthalpy-driven binding, respectively. In addition, q4MD-CD yielded better agreement between computed and experimental association rate constants. Using only binding free energy in the training set for parameterization was suggested to risk an incorrect entropy-enthalpy balance; therefore, binding enthalpy needs to be considered for optimizing parameters ^67^. With continuing growth in computer power, for molecular systems with fast association/dissociation rate constants, we suggest considering computed binding kinetics for validating and optimizing force-field parameters as well. Our studies also showed the importance of and challenge in correctly modeling multiple conformations in which solvent effects may be remarkable and experimental structures are not available. Although our preliminary studies indicated that using TIP3P and TIP4P water models did not yield different sampled conformations during MD simulations, other molecular systems may be more sensitive to the solvent effects with different water models. In the future, we envision a more careful force-field optimization that considers binding free energy, enthalpy-entropy balance and kinetic properties. We also anticipate further investigation into the role of water in the binding kinetics of various guests to a pocket with different polar and/or nonpolar properties ^88-89^.

## 5. Conclusion

In this work, we performed microsecond-timescale MD simulations with GPU acceleration for 7 chosen β-CD complexes. The computed thermodynamic and kinetic properties agree with experimental values reasonably well. The binding of β-CD complexes is mainly driven by the non-polar attraction between β-CD and the guest and the entropy gain of desolvated water molecules, regardless of the force field used. With both force fields, the ligands have only a small entropy penalty, and the entropy term also contributes to the binding process. GAFF-CD reproduced more favorable binding entropy and less favorable binding enthalpy due to stronger desolvation penalty than did q4MD-CD. As compared with GAFF-CD, q4MD-CD produced more rigid dynamics of free β-CD. However, the conformational rearrangement did not contribute to the differences in thermodynamics and kinetics modeled by the 2 force fields, and the real determinant was the different H-bond networks between the solvation water molecules and free β-CD. With GAFF-CD, free β-CD forms more H-bonds with solvation waters in a more distorted conformation; on ligand binding, the ligand needs to pay more enthalpic penalties to remove these stable H-bonded waters, which results in the stronger desolvation enthalpy. Removing the H-bonded waters also slows the association process. In the bound state, free β-CD forms fewer H-bonds with its environment and thus becomes more flexible and gains configuration entropy. Although we computed ΔH and –TΔS separately, the compensation is observed. Our study also showed that different force-field parameters can yield the same computed ΔG but different entropy and entropy balance.

## Acknowledgements

We thank support from the US National Institute of Health (GM-109045), US National Science Foundation (MCB-1350401), and NSF national super computer centers (TG-CHE130009). We also thank Dr. Michael Gilson and Dr. Niel Henriksen for insightful discussions on β-cyclodextrin systems and Dr. Ron Levy for sharing OPLS force field parameters for β-cyclodextrin for our tests.

---

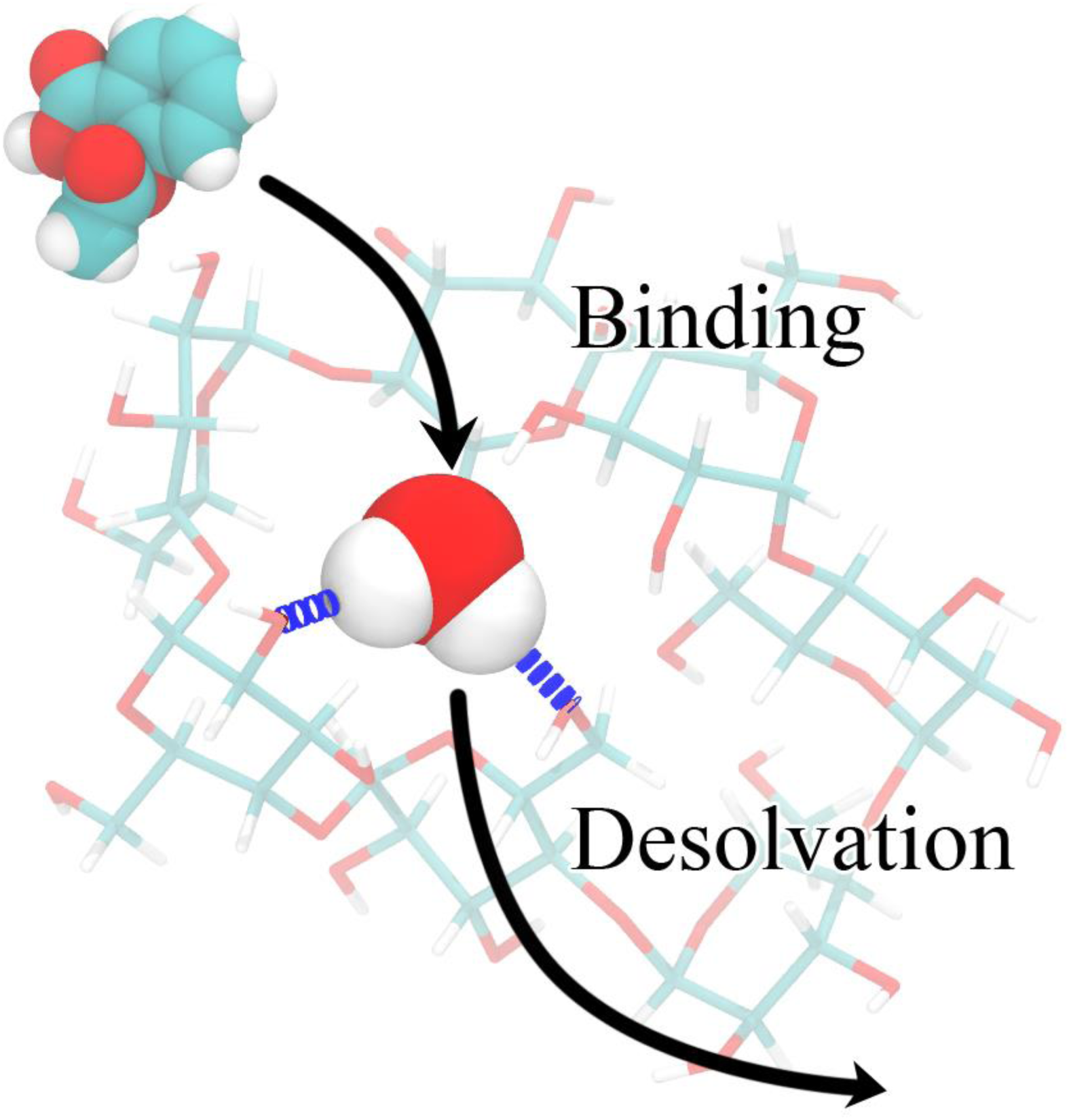

## Supporting Information

### 1. Lengths of MD Simulations

**Table SI 1.** The lengths of MD simulations of β-cyclodextrin (β-CD) complexes and numbers of frames used in calculations of thermodynamic properties. We used GAFF-CD and q4MD-CD for free β-CD and the complexes. We used GAFF for the free ligands and TIP3P for water in all simulations. All trajectories are resaved every 10 ps for thermodynamics calculations. The lengths used in calculations of thermodynamic properties are shown in column 4. Column 5 shows the total numbers of frames in free β-CD, free ligands, and water trajectories, and the numbers of bound state conformations in complex trajectories corresponding to the length indicated in column 4. Only bound state complex conformations were used in thermodynamics calculations. All full lengths of trajectories were used in association and dissociation rate constant calculations.

| Force Field | System | Total (μs) | Length Used (μs) | # of Bound State Conformations |
| --- | --- | --- | --- | --- |
| GAFF | 1-Propanol | 1.00 | 1.00 | 100000 |
|  | 1-Butanol | 2.56 | 2.56 | 256000 |
|  | Methyl Butyrate | 1.00 | 1.00 | 100000 |
|  | t-Butanol | 1.00 | 1.00 | 100000 |
|  | 1-Naphthyl Ethanol | 2.00 | 2.00 | 200000 |
|  | Aspirin | 1.00 | 1.00 | 100000 |
|  | 2-Naphthyl Ethanol | 2.50 | 2.50 | 250000 |
| TIP3P | Water | 2.50 | 2.50 | 250000 |
| GAFF-CD | β-CD-1-Propanol | 6.50 | 6.50 | 23256 |
| | β-CD-1-Butanol | 6.50 | 6.50 | 48869 |
| | β-CD-Methyl Butyrate | 7.50 | 7.50 | 111496 |
| | β-CD-t-Butanol | 8.00 | 8.00 | 86200 |
| | β-CD-1-Naphthyl Ethanol | 11.00 | 4.00 | 369283 |
| | β-CD-Aspirin | 9.50 | 5.50 | 453449 |
| | β-CD-2-Naphthyl Ethanol | 11.00 | 4.50 | 319300 |
| | β-CD | 2.50 | 2.50 | 250000 |
| q4MD-CD | β-CD-1-Propanol | 4.50 | 2.50 | 60225 |
| | β-CD-1-Butanol | 5.00 | 2.50 | 152864 |
| | β-CD-Methyl Butyrate | 4.50 | 2.50 | 214756 |
| | β-CD-t-Butanol | 4.50 | 2.50 | 214310 |
| | β-CD-1-Naphthyl Ethanol | 6.00 | 2.50 | 236130 |
| | β-CD-Aspirin | 6.00 | 2.50 | 241138 |
| | β-CD-2-Naphthyl Ethanol | 6.00 | 4.50 | 442508 |
| | β-CD | 2.60 | 2.60 | 260000 |

### 2. Conformations of β-CD in Vacuum MD Simulations in GAFF-CD and q4MD-CD

We performed MD simulations of free β-CD in vacuum for 50 ns using GAFF-CD and q4MD-CD respectively. Periodic condition was not considered. The trajectories were saved every 10 ps and totally 5000 frames were saved for the two MD runs. The RMSDs were computed against the crystal structures and plotted in Figure SI 1.

**Figure SI 1.**
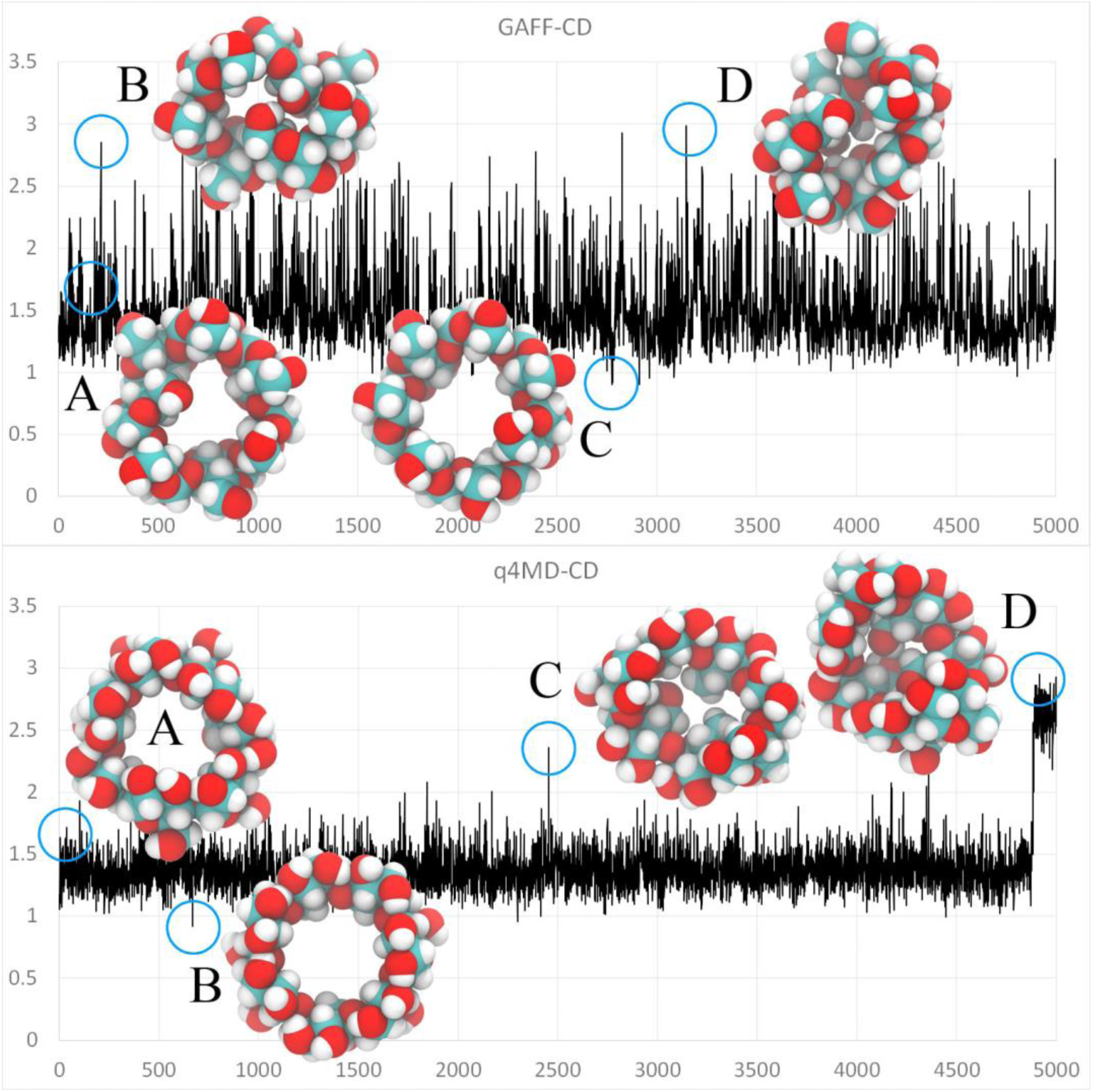
The RMSD plots and representative conformations of β-CD from MD simulations in vacuum in GAFF-CD and q4MD-CD. X-axes of RMSD plots are numbers of frames, and y-axes are RMSD against crystal structures in Å. Representative conformations are circled on the RMSD plots. Conformations A on the two plots have RMSD roughly equal to 1.7 Å. For GAFF-CD simulation, 80.5% conformations have RMSD smaller than 1.7 Å. For q4MD-CD simulations, the percentage is 95.12%.

### 3. Details of Enthalpy Decomposition Calculations

The total enthalpy change (ΔH) is decomposed into changes of solute-solute interaction (ΔH_Host-Guest_) and desolvation energy (ΔH_desolvation_), where ΔH_desolvation_ includes interaction changes of host-water (ΔH_Host-Water_), guest-water (ΔH_Guest-Water_), and water-water (ΔH_Water-Water_). Because both conformational changes and intermolecular interactions contribute to ΔH_Host-Guest_, we further provide decomposition analysis of ΔH_Host-Guest_ into changes of host and guest conformational enthalpy ΔH_Host Conf_ and ΔH_Guest Conf_, respectively, and their interaction energy change (ΔH_Solute Inter_). To compute these decomposed energy terms, we also calculated average potential energies of water (<E_Water_>_Complex_), water+host (<E_Water+Host_>_Complex_), water+guest (<E_Water+Guest_>_Complex_), host+guest (<E_Host+Guest_>_Complex_), host (<E_Host_>_Complex_) and guest (<E_Guest_>_Complex_) in the complex trajectories, water (<E_Water_>_Host_), water+host (<E_Water+Host_>_Host_) and host (<E_Host_>_Host_) in the free β-CD trajectory, water (<E_Water_>_Guest_), water+guest (<E_Water+Guest_>_Guest_) and guest (<E_Guest_>_Guest_) in the free ligand trajectories, in addition to <E>_Complex_, <E>_Water_, <E>_Host_ and <E>_Guest_. Then we used eqs. 3.1 to 3.8 to calculate the decomposed energy terms.

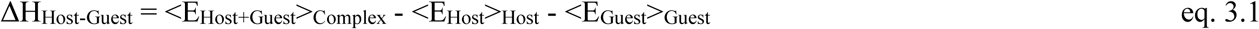

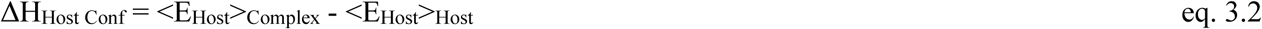

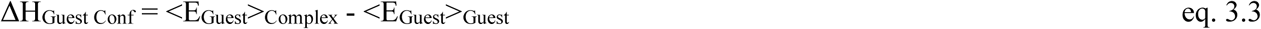

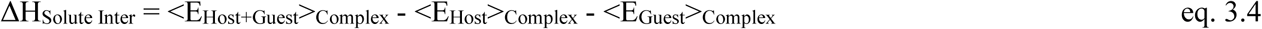

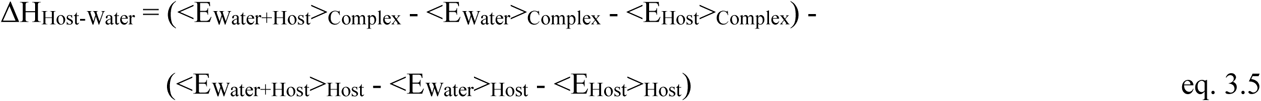

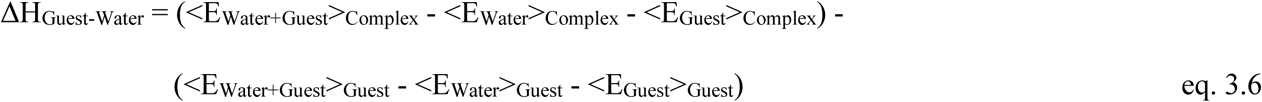

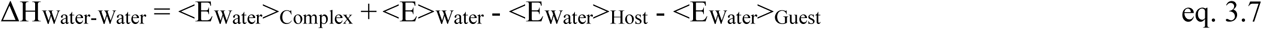

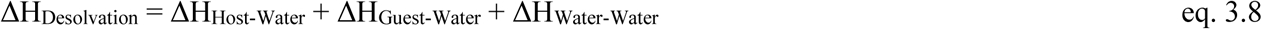

### 4. Surface Area, Hydrogen Bond and Solvation Water

**Figure SI 2.**
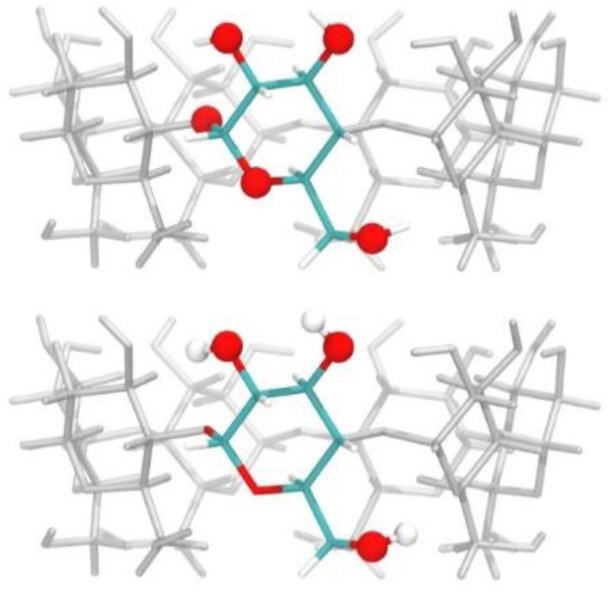
The hydrogen-bonding (H-bond) sites on β-CD. β-CD has totally 56 H-bonding sites, and on each of the seven glucopyranose units there are five H-bond acceptors (top) and three H-bond donors (bottom). The H-bonding sites are shown in spheres.

We evaluated the total solvent accessible surface area (SA_Total_), polar solvent accessible surface area (SA_Polar_) and non-polar solvent accessible surface area (SA_Non Polar_) of β-CD. The solvent accessible surface area was evaluated by generating spherical meshes with radius set to the van der Waals radii of atoms plus 1.4 Å on the atoms and counting the exposed surface areas of the meshes. SA_Total_, SA_Polar_ and SA_Non Polar_ are evaluated by summing up the surface areas of all atoms, of oxygen atoms, and of carbon and hydrogen atoms, respectively. Values are averaged from all conformations of free β-CD and bound state conformations of complex trajectories.

To estimate the H-bonds with water molecules, we computed the average occupancy percentages of the 56 H-bonding sites (Figure SI 2) on β-CD. We only counted the H-bonds formed with water molecules on these H-bonding sites. For each H-bonding site, the number of conformations with water molecules forming H-bond to the H-bonding site was divided by the total number of conformations to compute the percentage of water H-bond occupancy percentage. The percentages of each H-bonding site were then averaged to compute the percentage of the conformation. Finally, the percentage of conformations were then average for all conformations of free β-CD or bound state conformations of complexes to compute the average H-bond occupancy percentage (Δ%_H-Bond_).

The changes of total surface area (ΔSA_Total_), polar surface area (ΔSA_Polar_), non-polar surface area (ΔSA_Non Polar_), and average percentage of H-bond occupancy (Δ%_H-Bond_) are calculated by subtracting values of free β-CD from values of complexes. The results are summarized in Table SI 2.

**Table SI 2.** The solvent accessible surface area and H-bonds with water of β-CD in complexes and free state in GAFF-CD and q4MD-CD. The total surface area (SA_Total_), polar surface area (SA_Polar_) and non-polar surface area (SA_Non Polar_) are in Å^2^. The average H-bond occupancy percentage of 56 H-bond sites on β-CD (%_H-Bond_) are in %. The corresponding changes (Δ) are calculated by subtracting values of free β-CD from values of complexes.

| GAFF-CD |  |  |  |  |  |  |  |  |
| --- | --- | --- | --- | --- | --- | --- | --- | --- |
| Guest | SA<sub>Total</sub> | SA<sub>Polar</sub> | SA<sub>Non Polar</sub> | %<sub>H-Bond</sub> | ΔSA<sub>Total</sub> | ΔSA<sub>Polar</sub> | ΔSA<sub>Non Polar</sub> | Δ%<sub>H-Bond</sub> |
| β-CD-1-Propanol | 1019 | 216 | 803 | 34.25% | -37 | -6 | -31 | -3.58% |
| β-CD-1-Butanol | 1024 | 217 | 807 | 32.17% | -32 | -5 | -27 | -5.66% |
| β-CD-Methyl Butyrate | 998 | 209 | 789 | 32.31% | -58 | -13 | -45 | -5.52% |
| β-CD-t-Butanol | 1038 | 221 | 817 | 32.78% | -18 | -1 | -17 | -5.04% |
| β-CD-1-Naphthyl Ethanol | 994 | 211 | 783 | 29.64% | -62 | -11 | -52 | -8.18% |
| β-CD-Aspirin | 1002 | 212 | 790 | 29.86% | -54 | -10 | -44 | -7.97% |
| β-CD-2-Naphthyl Ethanol | 986 | 210 | 776 | 29.86% | -70 | -12 | -58 | -7.97% |
| β-CD | 1056 | 222 | 834 | 37.83% |  |  |  |  |
| q4MD-CD |  |  |  |  |  |  |  |  |
| Guest | SA<sub>Total</sub> | SA<sub>Polar</sub> | SA<sub>Non Polar</sub> | %<sub>H-Bond</sub> | ΔSA<sub>Total</sub> | ΔSA<sub>Polar</sub> | ΔSA<sub>Non Polar</sub> | Δ%<sub>H-Bond</sub> |
| β-CD-1-Propanol | 1058 | 228 | 829 | 28.74% | -66 | -5 | -61 | -2.41% |
| β-CD-1-Butanol | 1045 | 226 | 819 | 28.64% | -78 | -7 | -71 | -2.51% |
| β-CD-Methyl Butyrate | 1030 | 222 | 808 | 28.09% | -94 | -12 | -82 | -3.05% |
| β-CD-t-Butanol | 1055 | 231 | 824 | 28.44% | -68 | -2 | -66 | -2.70% |
| β-CD-1-Naphthyl Ethanol | 995 | 213 | 782 | 26.61% | -129 | -21 | -108 | -4.54% |
| β-CD-Aspirin | 1005 | 216 | 789 | 27.31% | -118 | -17 | -101 | -3.83% |
| β-CD-2-Naphthyl Ethanol | 993 | 212 | 781 | 27.18% | -130 | -21 | -109 | -3.96% |
| β-CD | 1123 | 234 | 890 | 31.14% |  |  |  |  |

### 5. Definition of Probability Distribution of Conformations of Solutes

**Figure SI 3.**
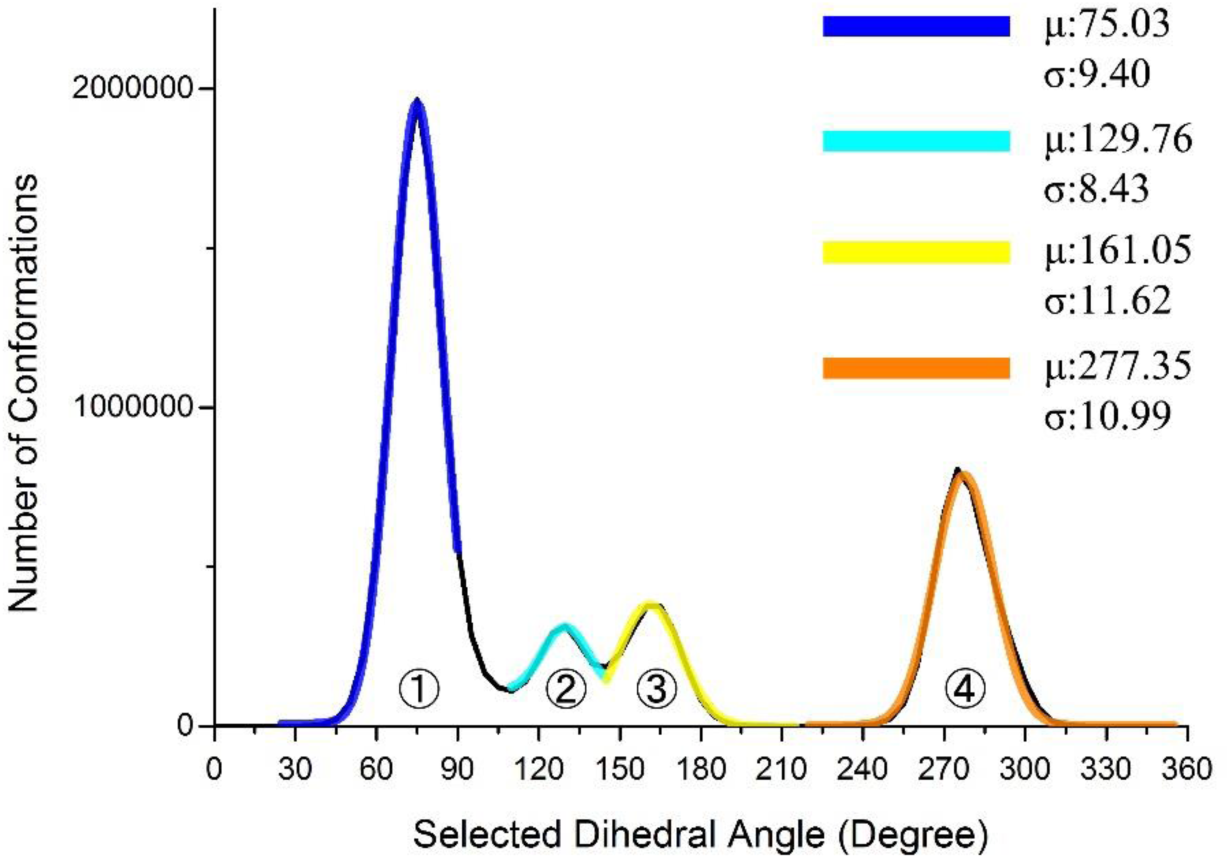
A representative dihedral population histogram used to define the conformation of individual dihedral angles. The population distribution is in black. Four Gaussian functions (1 in Blue, 2 in cyan, 3in yellow and 4 in orange) were fitted onto the four population peaks and the averages (μ) and standard deviations (σ) are labeled.

We used the combinations of dihedral angles to define the conformations of β-CD and guest molecules, and used the probability distribution of conformations in the Gibbs entropy formula (eq. 1 in main text) to compute the internal entropy of β-CD and guest molecules. 14 dihedrals defined in Figure 1 were selected for β-CD while every dihedral except methyl and hydroxyl rotations were selected for the ligands. We obtained the population histograms of the dihedral angles from MD simulation. We defined the conformation of selected dihedral angles use the Gaussian functions illustrated in Figure SI 3. Given any dihedral angle, ***α***, we define it in dihedral conformation ***i*** if,

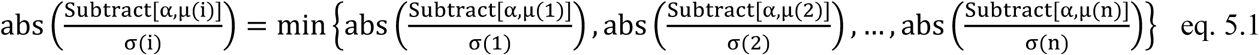

where *abs(****x***) is to take the absolute value of ***x***, *μ(****i***) and *σ(****i***) indicate the average and standard deviation of Gaussian function ***i***, *min(…)* is to take the minimum value of given values, the ***n*** denotes the total number of peaks of the selected dihedral, and *Subtract* is an operator defined as eq. 5.2 used to subtract two dihedral angles,

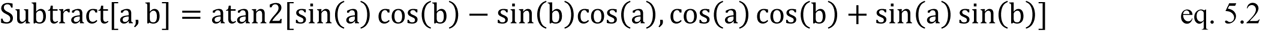

where *atan2* is the standard arctangent function that convert from values of sin(θ) and cos(θ) to θ. We treated conformations that fall into each peak as one identical conformation of the corresponding dihedral angle, and neglected the width of the peaks shown in Figure SI 3.

With the definition of individual dihedral angles, we used the different combinations of individual dihedral conformations to define conformations of β-CD and ligands. Distinct combinations of dihedral angle conformations are considered as distinct conformations of β-CD and ligands. In β-CD case, its seven-fold symmetry was considered. The population distributions of β-CD and ligands were then calculated from the distinct conformations.

### 6. Solvation Entropy Calculation Using Water Molar Entropy

Using grid cell method ^1-3^, the water molar entropy was evaluated by separating it into vibrational entropy (S_Water Vib_) and conformational entropy (S_Water Conf_), as in eq. 6.1,

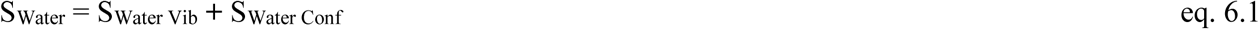

S_Water Vib_ is further separated into translational entropy (S_Water Trans_) and rotational entropy (S_Water Rot_) as in eq. 6.2,

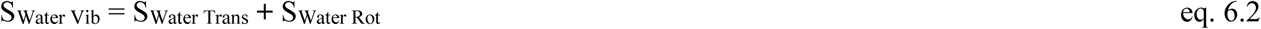

S_Water Trans_ and S_Water Rot_ are computed from forces and torques on the three principal axes of the target water molecule as in eqs. 6.3 and 6.4,

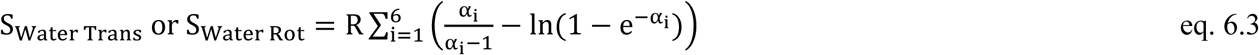

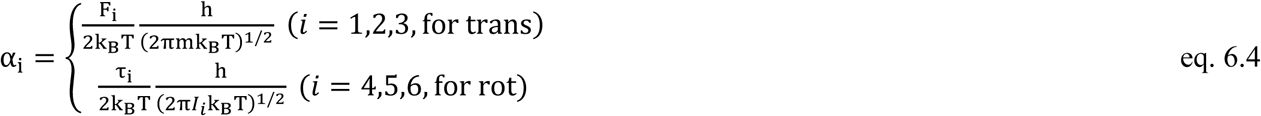

where, R is the gas constant, k_B_ is the Boltzmann constant, h is the Planck constant, T is the temperature, m is the mass of a water molecule, F_i_ are the force and τ_i_ are the torques on the three principal axes. The gas phase values of moments of inertia I_i_ about x, y, and z axes are used.

**Figure SI 4.**
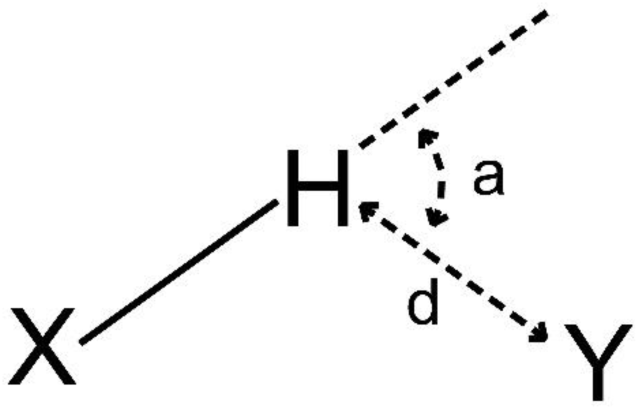
The definition of a loose H-bond. It is based on normal H-bond definition except that the complementary angle of X-H…Y (a) is less than 80° and the distance between H and Y (d) is less than 2.65Å.

**Figure SI 5.**
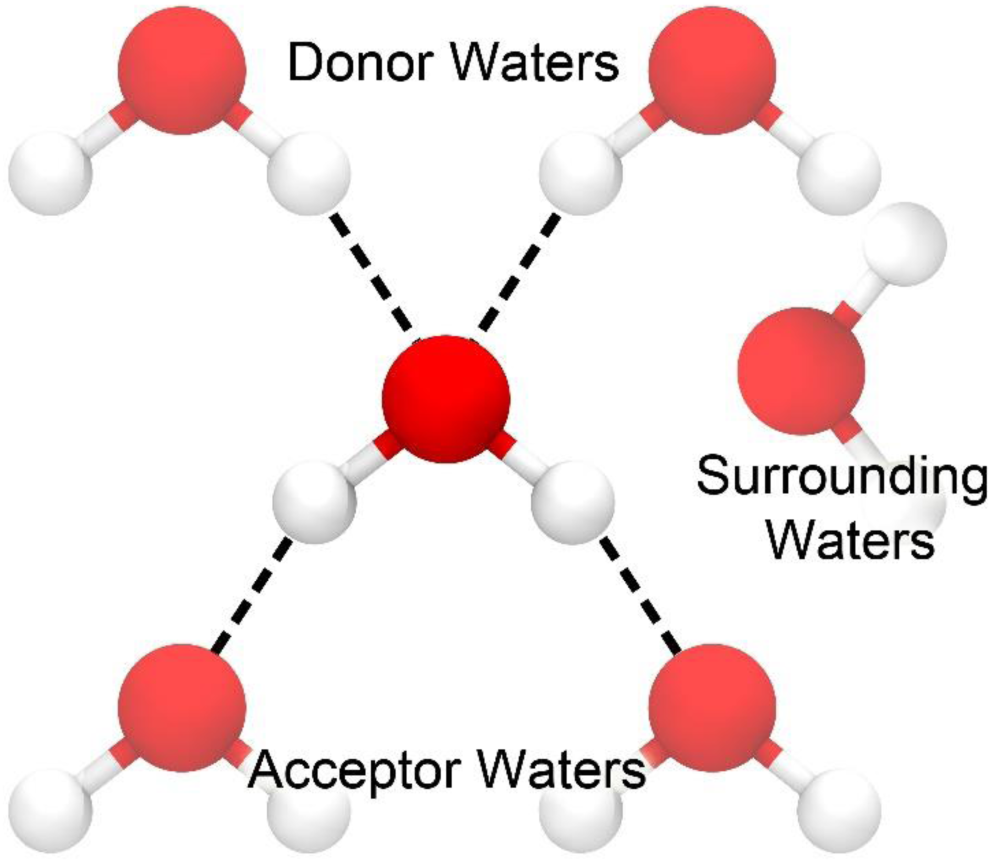
The water conformational entropy definition. It is based on the Pauling’s model. Around the target water molecule, the donor waters (DW) and acceptor waters (AW) are defined with the loose H-bond definition and any water molecule that is within 3.2 Å from the oxygen atom of the target water molecule but cannot form a loose H-bond is considered as a surrounding water (SW).

We derived model to evaluate water conformational entropy (S_Water conf_) based on Pauling’s model ^2^-^4^. First a loose H-bond is defined as shown in Figure SI 4, where the complementary angle of X-H…Y (a) must be less than 80° and the H…Y distance (d) must be less than 2.65 Å. The purpose of the loose definition is to capture the forming and breaking H-bonds due to thermal fluctuation. Based on that, the conformational entropy is defined according to Figure SI 5. The water molecules around the target water molecule are classified into three groups, 1) acceptor waters (AW), which are the loose H-bond acceptors from the target water, 2) donor waters (DW), which are the loose H-bond donors to the target water, and 3) surrounding waters (SW), whose oxygen atoms are within 3.2 Å of the oxygen atom of the target water molecule but form no H-bond with it. The number of conformations (Ω_Water_) of the target water is calculated using eq. 6.5,

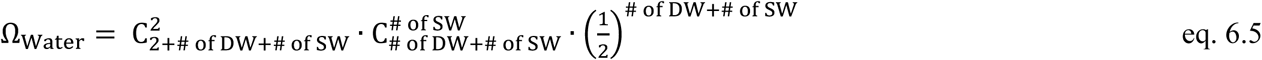

In the first term, both “2”s mean the two AW sites, regardless of the existence of water molecules there. This term can be interpreted as choosing two sites of the water molecules around as the acceptor sties. The second term means in the rest of the water around to choose # of SW of waters as the SW. The appearance of the last term is to compensate the double counting of water molecules around. For example, when there are exactly two AWs and two DWs, 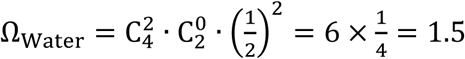, which obeys the Pauling’s model. When there are two AWs, two DWs and one SW, 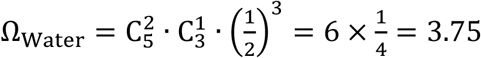, which means an additional water molecule around increases the number of conformations of the target water. Finally, the conformational entropy of the water is calculated as eq. 6.6,

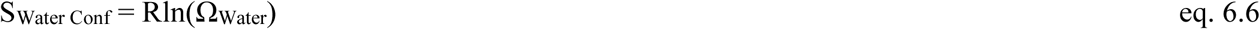

The solvation entropy can be calculated using water molar entropy. The solvation energy of species *X* can be computed using eq. 6.7,

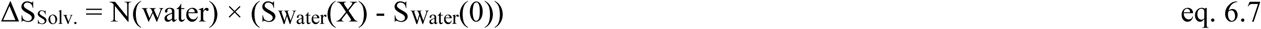

where *N(water)* is given as *# of Water* in Tables SI 3 to 5, *S*_*Water*_*(X)* is the molar water entropy with species *X*, and *S*_*Water*_*(0)* is the molar water entropy of bulk water. The computed water molar entropy and solvation entropy are summarized in Tables SI 3 to 5.

**Table SI 3.** The molar water entropy and solvation entropy of free β-CD and β-CD complexes in GAFF-CD. The molar water entropy (S_Water_) is separated into vibrational (S_Water Vib_), and conformational (S_Water Conf_) terms. The vibrational (S_Water Vib_) term is further separated into translational (S_Water Trans_) and rotational (S_Water Rot_) terms. The molar water entropy terms are in J/mol/K. The solvation entropy (-TΔS_Solv._) is separated into vibrational (-TΔS_Solv. Vib_), and conformational (-TΔS_Solv. Conf_) terms. The vibrational (-TΔS_Solv. Vib_) term is further separated into translational (-TΔS_Solv. Trans_) and rotational (-TΔS_Solv. Rot_) terms. The solvation energy terms at 298K are in kcal/mol. The numbers of water molecules are corresponding to a volume of 13520 Å^3^.

| System | $S_{\text{Water}}$ (J/mol/K) | | | | | $-T\Delta S_{\text{Solv.}}$ (kcal/mol) | | | | | # of Water |
| --- | --- | --- | --- | --- | --- | --- | --- | --- | --- | --- | --- |
|  | Trans | Rot | Vib | Conf | Total | Trans | Rot | Vib | Conf | Total |  |
| β-CD-1-Propanol | 47.69 | 20.83 | 68.52 | 5.01 | 73.53 | 4.62 | 0.59 | 5.21 | 2.48 | 7.69 | 403.01 |
| β-CD-1-Butanol | 47.7 | 20.85 | 68.54 | 5.01 | 73.55 | 4.44 | 0.23 | 4.66 | 2.53 | 7.19 | 401.58 |
| β-CD-Methyl Butyrate | 47.7 | 20.84 | 68.54 | 5.01 | 73.55 | 4.19 | 0.45 | 4.64 | 2.48 | 7.12 | 400.41 |
| β-CD-t-Butanol | 47.71 | 20.84 | 68.55 | 5.01 | 73.56 | 4.03 | 0.43 | 4.45 | 2.48 | 6.93 | 401.74 |
| β-CD-1-Naphthyl Ethanol | 47.72 | 20.85 | 68.56 | 5.01 | 73.57 | 3.74 | 0.21 | 3.95 | 2.46 | 6.41 | 397.53 |
| β-CD-Aspirin | 47.71 | 20.85 | 68.56 | 5.01 | 73.57 | 3.91 | 0.24 | 4.15 | 2.45 | 6.59 | 398.39 |
| β-CD-2-Naphthyl Ethanol | 47.72 | 20.85 | 68.57 | 5.01 | 73.58 | 3.69 | 0.13 | 3.82 | 2.47 | 6.29 | 397.49 |
| β-CD | 47.67 | 20.83 | 68.5 | 5.02 | 73.52 | 5.1 | 0.78 | 5.88 | 2.19 | 8.07 | 407.14 |

**Table SI 4.** The molar water entropy and solvation entropy of free β-CD and β-CD complexes in q4MD-CD. Refer to Table SI 3 for meanings of symbols.

| System | $S_{\text{Water}}$ (J/mol/K) | | | | | $-T\Delta S_{\text{Solv.}}$ (kcal/mol) | | | | | # of Water |
| --- | --- | --- | --- | --- | --- | --- | --- | --- | --- | --- | --- |
|  | Trans | Rot | Vib | Conf | Total | Trans | Rot | Vib | Conf | Total |  |
| β-CD-1-Propanol | 47.71 | 20.85 | 68.56 | 5.01 | 73.57 | 4.11 | 0.09 | 4.20 | 2.35 | 6.55 | 401.93 |
| β-CD-1-Butanol | 47.71 | 20.85 | 68.57 | 5.01 | 73.58 | 3.89 | -0.02 | 3.86 | 2.30 | 6.16 | 400.86 |
| β-CD-Methyl Butyrate | 47.72 | 20.86 | 68.58 | 5.01 | 73.59 | 3.71 | -0.16 | 3.55 | 2.35 | 5.90 | 400.01 |
| β-CD-t-Butanol | 47.72 | 20.85 | 68.57 | 5.01 | 73.58 | 3.82 | 0.05 | 3.88 | 2.34 | 6.22 | 401.15 |
| β-CD-1-Naphthyl Ethanol | 47.72 | 20.87 | 68.59 | 5.01 | 73.60 | 3.57 | -0.49 | 3.08 | 2.34 | 5.43 | 397.77 |
| β-CD-Aspirin | 47.72 | 20.86 | 68.58 | 5.01 | 73.59 | 3.68 | -0.22 | 3.46 | 2.32 | 5.77 | 398.05 |
| β-CD-2-Naphthyl Ethanol | 47.73 | 20.87 | 68.60 | 5.01 | 73.61 | 3.44 | -0.60 | 2.83 | 2.38 | 5.22 | 397.45 |
| β-CD | 47.69 | 20.84 | 68.54 | 5.02 | 73.56 | 4.50 | 0.27 | 4.77 | 2.20 | 6.98 | 406.33 |

**Table SI 5.**
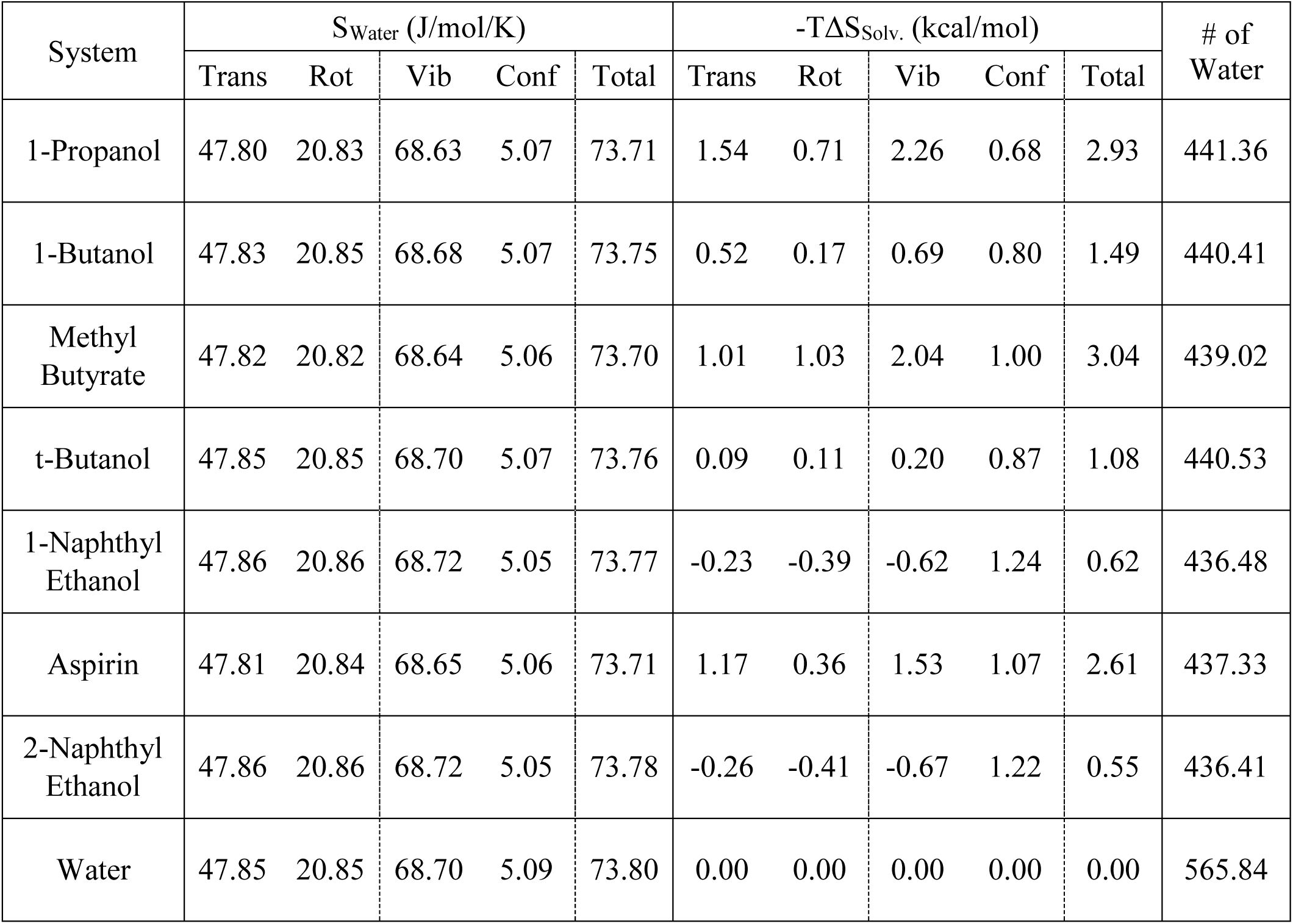
The molar water entropy and solvation entropy of free ligands and water. Refer to Table SI 3 for meanings of symbols.

Alternatively, the solvation water entropy can be computed using forces (F_i, S_), torques (τ_i, S_), and number of water conformation (Ω_S_) directly using eqs. 6.8 to 6.11, instead of computing the water molar entropy first,

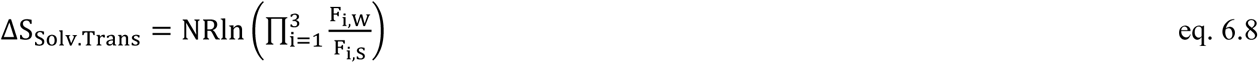

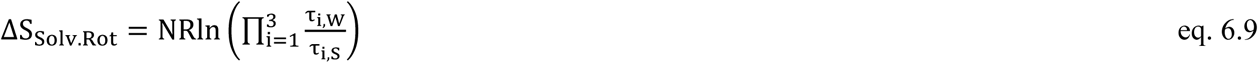

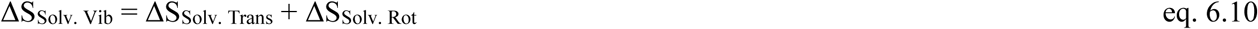

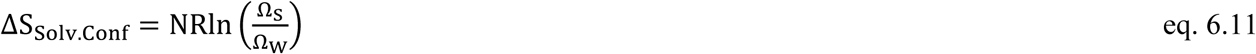

where, R is the gas constant, F_i, W_ and F_i, S_ are the averages of forces on three principal axes without solute and with solute, τ_i, W_ and τ_i, S_ are the averages of torques on three principal axes without and with solute, Ω_W_ and Ω_S_ are the average of number of conformations of water molecules without and with solute, and N is the number of water molecules in the sub water box defined in Figure SI 6.

**Figure SI 6.**
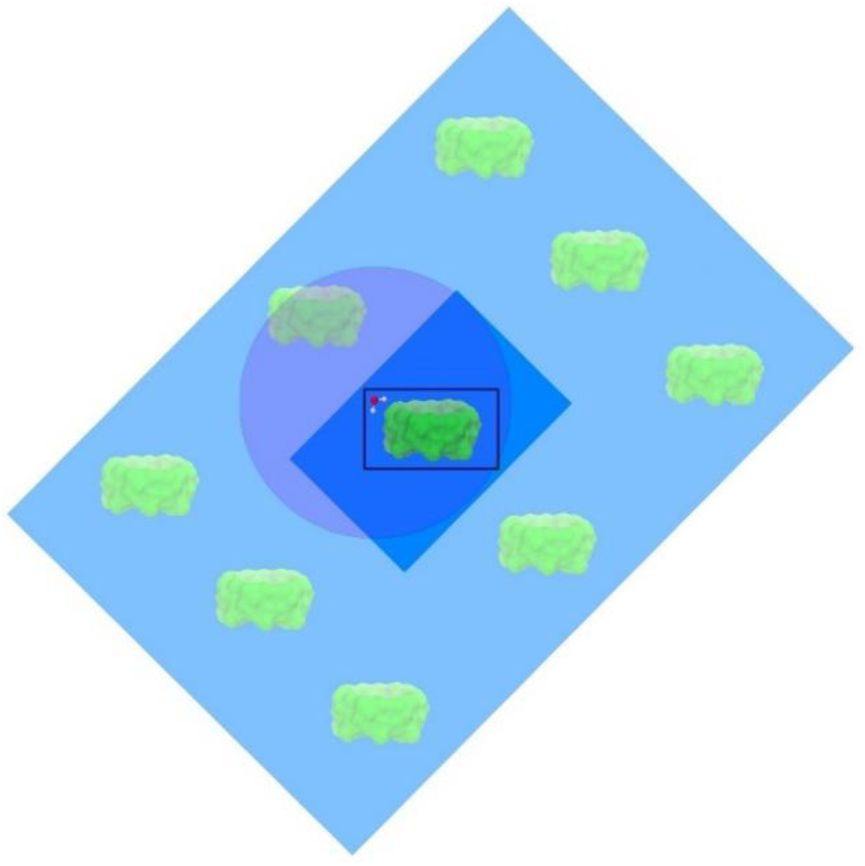
Illustration of water entropy calculation using free β-CD as representation. The original water box (deep blue) is align to the crystal structure of β-CD (deep green). One representative water molecule under calculation is shown in the sub water box (black lined box). Any periodic instances of atoms within the cutoff (purple circle) generated from original water box and periodic images of the original water box (light blue and light green) are used to calculate the forces and torques on the water molecule under calculation.

Practically, the complex and β-CD trajectories were aligned against the crystal structure of free β-CD. The ligand trajectories were aligned against the initial conformation of the ligand. Water molecules in a sub water box centered on the β-CD (in the case of free β-CD and complexes) and guest (in the case of free guests) with a size of 20Å ×26Å ×26 Å were considered in the water entropy calculation. For the empty water box, a sub water box with size 22Å ×22Å ×22 Å was used. A cutoff 50.0 Å was used for force and torque calculation in the vibrational term, and periodical atom instances generated from the full water box within this cutoff were used to calculate the forces and torques on the water molecules in the sub water box. The grid bin size was set to 1 Å in the grid cell method. The illustration of the water box for water entropy calculation is in Figure SI 6.

### 7. Solvation Water

We also counted the numbers of water molecules in the first sovlation shells of free guest (#_Guest_), free β-CD (#_Host_) and the complex (#_Complex_) from the MD trajectory. We counted the number oxygen atoms in the water molecules within 3.5 Å of all atoms in the corresponding solute and took the average of the number of waters over the entire trajectory for free guest and free host and over the bound state conformations in trajectories for complexes. The changes of the numbers of water molecules in the first solvation shells upon ligand binding (Δ#) are calculated by #_Guest_ + #_Host_ – #_Complex_. The results are summarized in Table SI 6.

**Table SI 6.**
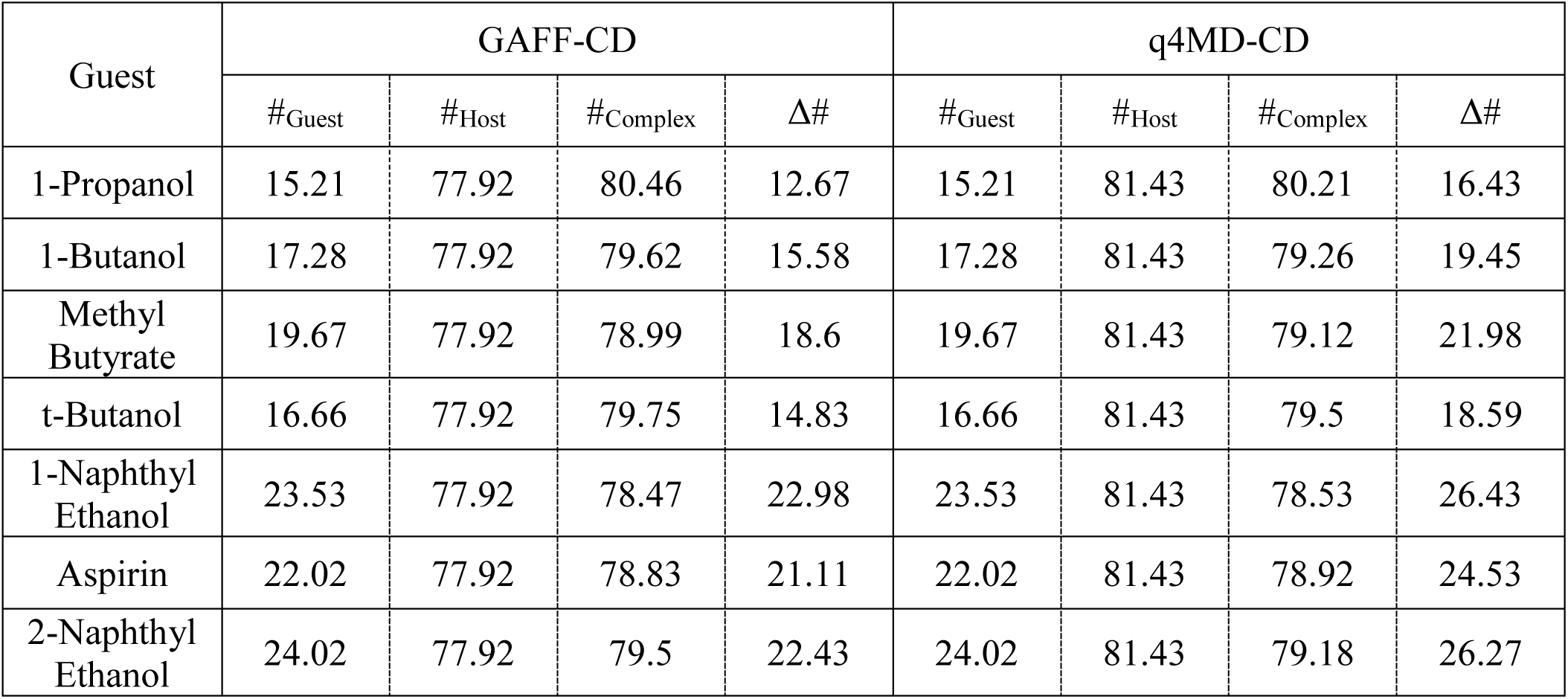
The number of water molecules in the first solvation shells of free guest (#_Guest_), free host (#_Host_), complex (#_Complex_) and number of water molecules released from the first solvation shell to the bulk (Δ#). Values are obtained by averaging the numbers of water molecules within 3.5 Å of atoms in the solute over the corresponding trajectories. The change is computed by #_Guest_ + #_Host_ – #_Complex_.

### 8. Solute Concentrations Used in Association Rate Constant Calculations

We used this protocol to calculate the concentration used in association rate constant (k_on_) calculation and the concentrations are summarized in Table SI 7. We obtained the cell sizes of free state conformations in the complex trajectories saved in original trajectories, and used eq. 8.1 to calculate the volume of the water box,

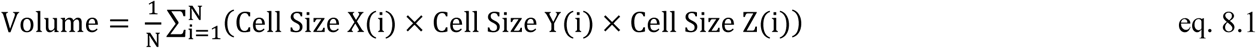

where N is the number of free state conformations in the complex trajectory, and Cell Size X, Y, Z are the cell sizes of the water box in three dimensions.

Then we used eq. 8.2 to calculate the solute concentration ([solute]) of the free state in the complex trajectories,

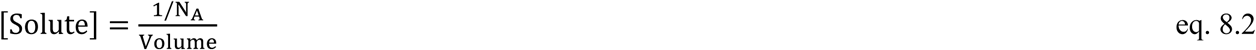

where N_A_ is the Avogadro constant, and “1” is the number of complex particles in the trajectory.

**Table SI 7.** The solute concentrations of β-CD and guest molecules used in association rate constant (k_on_) calculation in GAFF-CD and q4MD-CD. The concentrations correspond to free state conformations in the complex trajectories. All values are in mol/L (M).

| Guest | GAFF-CD | q4MD-CD |
| --- | --- | --- |
| 1-Propanol | 0.03079 | 0.03079 |
| 1-Butanol | 0.03078 | 0.03078 |
| Methyl butyrate | 0.03076 | 0.03076 |
| t-Butanol | 0.03078 | 0.03077 |
| 1-Naphthyl Ethanol | 0.03070 | 0.03072 |
| Aspirin | 0.03072 | 0.03072 |
| 2-Naphthyl Ethanol | 0.03070 | 0.03071 |
| Unit: M |  |  |

### 9. Diffusion Controlled Association Rate Constants Estimated from Diffusion Coefficient

**Table SI 8.** Diffusion coefficients of ligands and β-CD and diffusion controlled association rate constants (k_on_diffuse_) estimated from the diffusion coefficients. Diffusion coefficients (in m^2^/s) are estimated by eq. 9.1 using water viscosities from experiments (D_Expt_) and TIP3P water model (D_TIP3P_), and directly measured from MD trajectories (D_MD_). k_on_diffuse_ values are estimated using D_MD_ in eq. 9.2 for each β-CD-guest complex.

| System | $D_{\text{Expt}}$ ($\text{m}^2/\text{s}$) | $D_{\text{TIP3P}}$ ($\text{m}^2/\text{s}$) | $D_{\text{MD}}$ ($\text{m}^2/\text{s}$) | $k_{\text{on\_diffuse}}$ ($1/\text{s}\cdot\text{M}$) |
| --- | --- | --- | --- | --- |
| 1-Butanol | $8.7\times 10^{-10}$ | $2.4\times 10^{-9}$ | $5.9\times 10^{-9}$ | $4.45\times 10^{10}$ |
| Methyl Butyrate | $8.1\times 10^{-10}$ | $2.3\times 10^{-9}$ | $5.2\times 10^{-9}$ | $4.07\times 10^{10}$ |
| t-Butanol | $8.7\times 10^{-10}$ | $2.4\times 10^{-9}$ | $5.7\times 10^{-9}$ | $4.33\times 10^{10}$ |
| 1-Propanol | $9.4\times 10^{-10}$ | $2.6\times 10^{-9}$ | $6.6\times 10^{-9}$ | $4.76\times 10^{10}$ |
| 1-Naphthyl Ethanol | $7.1\times 10^{-10}$ | $2.0\times 10^{-9}$ | $3.8\times 10^{-9}$ | $3.26\times 10^{10}$ |
| Aspirin | $7.3\times 10^{-10}$ | $2.0\times 10^{-9}$ | $3.7\times 10^{-9}$ | $3.14\times 10^{10}$ |
| 2-Naphthyl Ethanol | $7.0\times 10^{-10}$ | $2.0\times 10^{-9}$ | $3.8\times 10^{-9}$ | $3.28\times 10^{10}$ |
| β-CD | $4.1\times 10^{-10}$ | $1.1\times 10^{-9}$ | $7.2\times 10^{-10}$ | / |

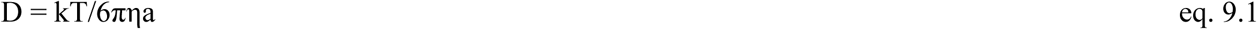

where k is the Boltzmann constant, T is the temperature (298K), η is the viscosity of water, and a is the radius of solute.

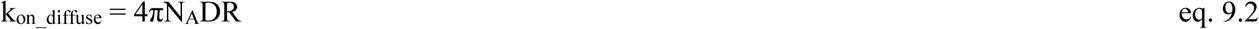

where N_A_ is the Avogadro constant, D is the sum of the diffusion coefficient of β-CD and guest, and R is the sum of radii of β-CD and guest.

### 10. Uncertainty Evaluation of Computed Properties

We evaluated the uncertainty of all computed properties. For potential energy terms (<E>_Complex_, <E>_Water_, <E>_Host_, and <E>_Guest_), we used standard deviation of mean to evaluate the uncertainty in the potential energy and it was calculated by using block analysis ^5^ with 2 ns in a block. The uncertainties of <E>_Complex_, <E>_Water_, <E>_Host_, and <E>_Guest_ were added in quadrature to get the uncertainty of binding enthalpy (ΔH). For properties *A* other than total binding enthalpy, we used bootstrap error analysis to get the uncertainties. We generated random subsets of the data, and calculated values of *A* from the random subsets, and finally calculated the standard deviations of the values of *A* from the random subsets, as the uncertainty of *A*. In principle, roughly 80% of the data were used to generate the random subsets. The number of random subsets is 100000 if non-repeated random subsets can be generated or otherwise the maximum number of possible subsets was used.

We computed the conformational population distribution of β-CD and used the Gibbs entropy formula (eq. 1) to evaluate S_Host Complex_, S_Host Free_, S_Guest Int Complex_ and S_Guest Int Free_. For S_Host Complex_, we generated the random subsets of the conformations, and reconstructed the conformational population distributions for each subset, and calculated the standard deviation of S_Host Complex_ obtained from each subset as the uncertainty of S_Host Complex_. We did identical evaluations for uncertainty of S_Host Free_, S_Guest Int Complex_ and S_Guest Int Free_. For example, for the free β-CD in GAFF-CD, we generated 100000 subsets with 200000 random conformations from the total 250000 conformations, and evaluated S_Host Free_ using the strategy above. The uncertainty of ΔS_Host_ and ΔS_Guest Int_ were evaluated by summing uncertainties of S_Host Complex_ and S_Host Free_ and by summing uncertainties of S_Guest Int Complex_ and S_Guest Int Free_, respectively. We evaluated S_Guest Ext Complex_ using eq. 2. Therefore, we used similar strategy as in internal entropy uncertainty evaluation to create the random subsets of conformations, and reconstructed the spatial and angular distribution histogram for evaluation of S_Guest Ext Complex_ in each subset using eq. 2. The uncertainty was computed, again, as the standard deviation of S_Guest Ext Complex_ obtained from the subsets. The uncertainty of solute entropy change (ΔS_Solute_) was evaluated by summing uncertainties of ΔS_Host_, ΔS_Guest Int_, and S_Guest Ext Complex_. Note that S_Guest Ext Free_ was computed using exact analytical equations so this term does not have uncertainty.

For uncertainties of S_Water Complex_, S_Water Host_ and S_Water Guest_, the calculation is slightly different in the way the random subsets were created. We saved the water entropy values for every 10-ps-block (1000 frames) along the trajectories and obtained N water entropy values from each block. Then we generated subsets that contain random collections of roughly 0.8N water entropy values from the N values. The percentage *P%* is changed for a suitable combination number C_N_, _P%N_ so that C_N_, _P%N_ is within 100000 to 4294967296 (the greatest number in unsigned integer or 2^32^). The standard deviations of water entropy values averaged from the random subsets were reported as the uncertainties of the water entropy terms. For example, for β-CD-1-butanol complex in GAFF, we used 45 blocks from the total 49 blocks to generate 211876 (C_49_, _45_) subsets, and evaluated the uncertainty using first 100000 subsets. The uncertainties of water entropy change (ΔS_Water_) were evaluated by summing up the uncertainties of S_Water Complex_, S_Water Host_ and S_Water Guest_.

The uncertainty of binding entropy (ΔS) was evaluated by summing up the uncertainties of ΔS_Solute_ and ΔS_Water_. The uncertainty of binding free energy from eq. 5 (ΔG_Comp1_) was evaluated by summing up the uncertainties of ΔH and ΔS.

For association (k_on_) and dissociation (k_off_) rate constants, we created the random subsets of the time lengths of bound/free periods, and calculated the rate constants from these subsets, and finally computed the standard deviations of the rate constants from these subsets, as the uncertainties of k_on_ and k_off_. The uncertainty of K_eq_ are evaluated based on subsets of random combinations of k_on_ and k_off_ values obtained from the random subsets used to evaluated uncertainty of k_on_ and k_off_. For example, for β-CD-1-propanol complex in GAFF-CD, we used 34 bound state period lengths out of total 39 to generate 575757 (C_39, 34_) random subsets, computed the uncertainty of k_on_ using the standard deviation of the k_on_ values from the first 100000 subsets.

### 11. Experimental Data of ΔG, ΔH and –TΔS of β-CD-Alcohol Complexes Using ITC and UV

**Table SI 9.**
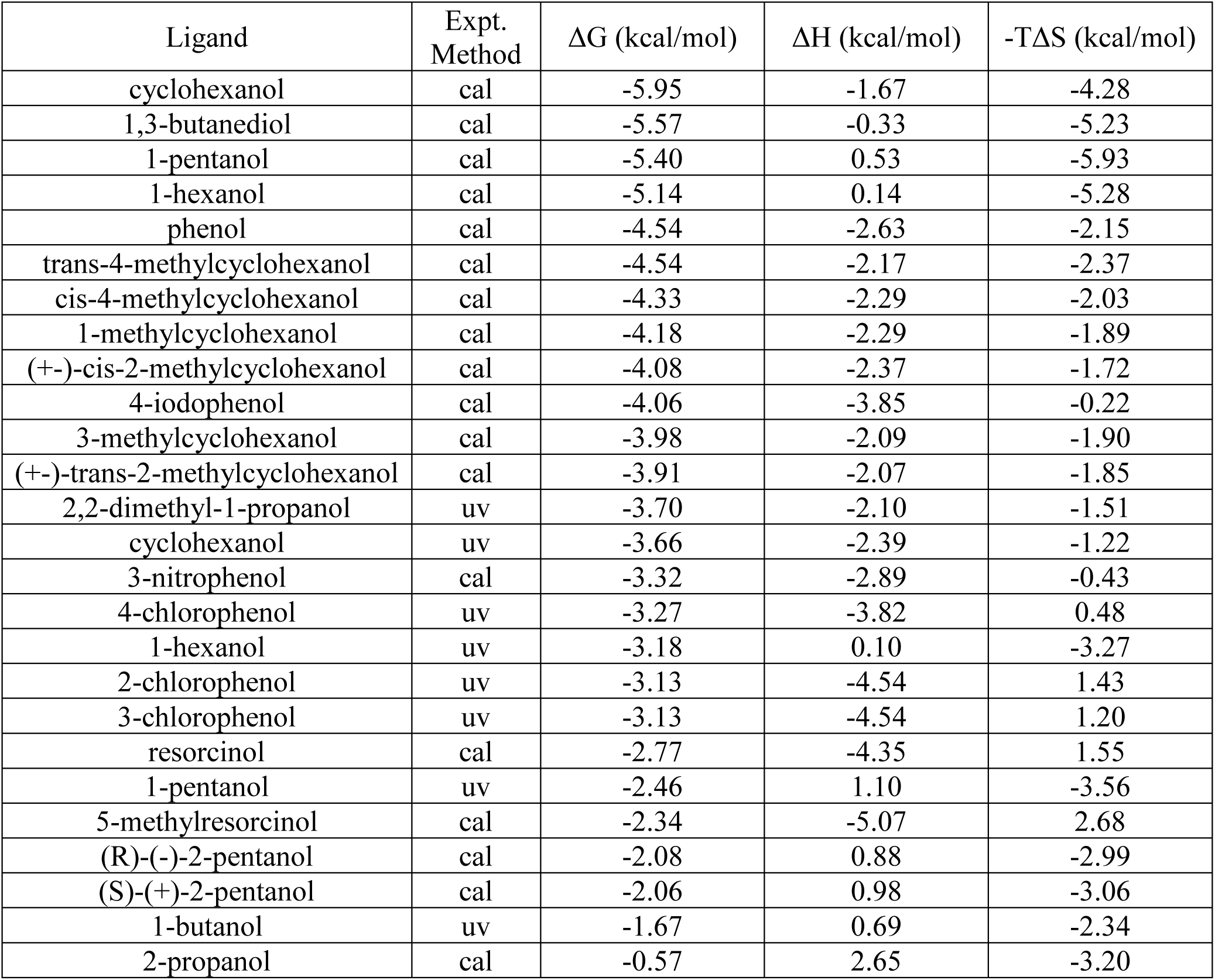
Experimental data of ΔG, ΔH and –TΔS of β-CD-alcohol complexes using calorimetry (cal) and spectrophotometry (uv). The data are taken from ref ^6^.

**Figure SI 7.**
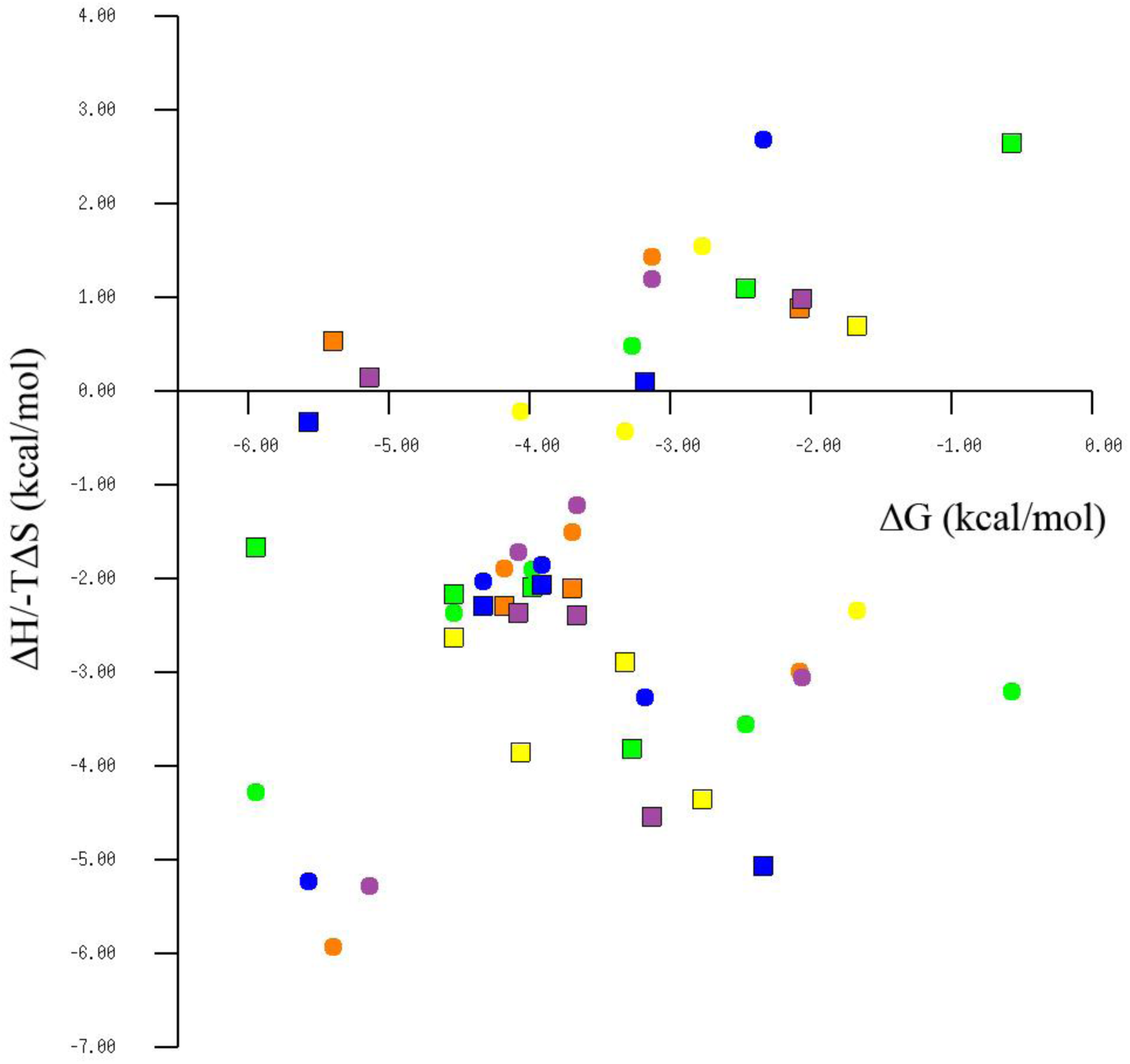
Plot of experimental ΔH and –TΔS against experimental ΔG of β-CD-alcohol complexes using calorimetry and spectrophotometry. The plot uses values from Table SI 9. Enthalpy values are indicated by bordered rectangle, and entropy values are indicated by circles. The enthalpy and entropy of the same ligand are rendered in the same color. For ligands with ΔG ranging from 0 to -3 and -5 to -6 kcal/mol, –TΔS is the major contribution to ΔG; for ligands with ΔG ranging from -3 to -5 kcal/mol, both ΔH and –TΔS contributes to ΔG.

### 12. Convergence of Enthalpy Calculations (<E>)

We investigated the convergence of enthalpy calculation by plotting average potential energy (<E> in kcal/mol) against number of frames used in the average in Figures SI 8 to 10. We calculate the standard deviation (SD) of the last region of the average potential energy to estimate the fluctuation in the calculated enthalpy. The regions used in the SD calculation are indicated on the figures. Note that the SD is different from the uncertainty reported in the manuscript.

**Figure SI 8.**
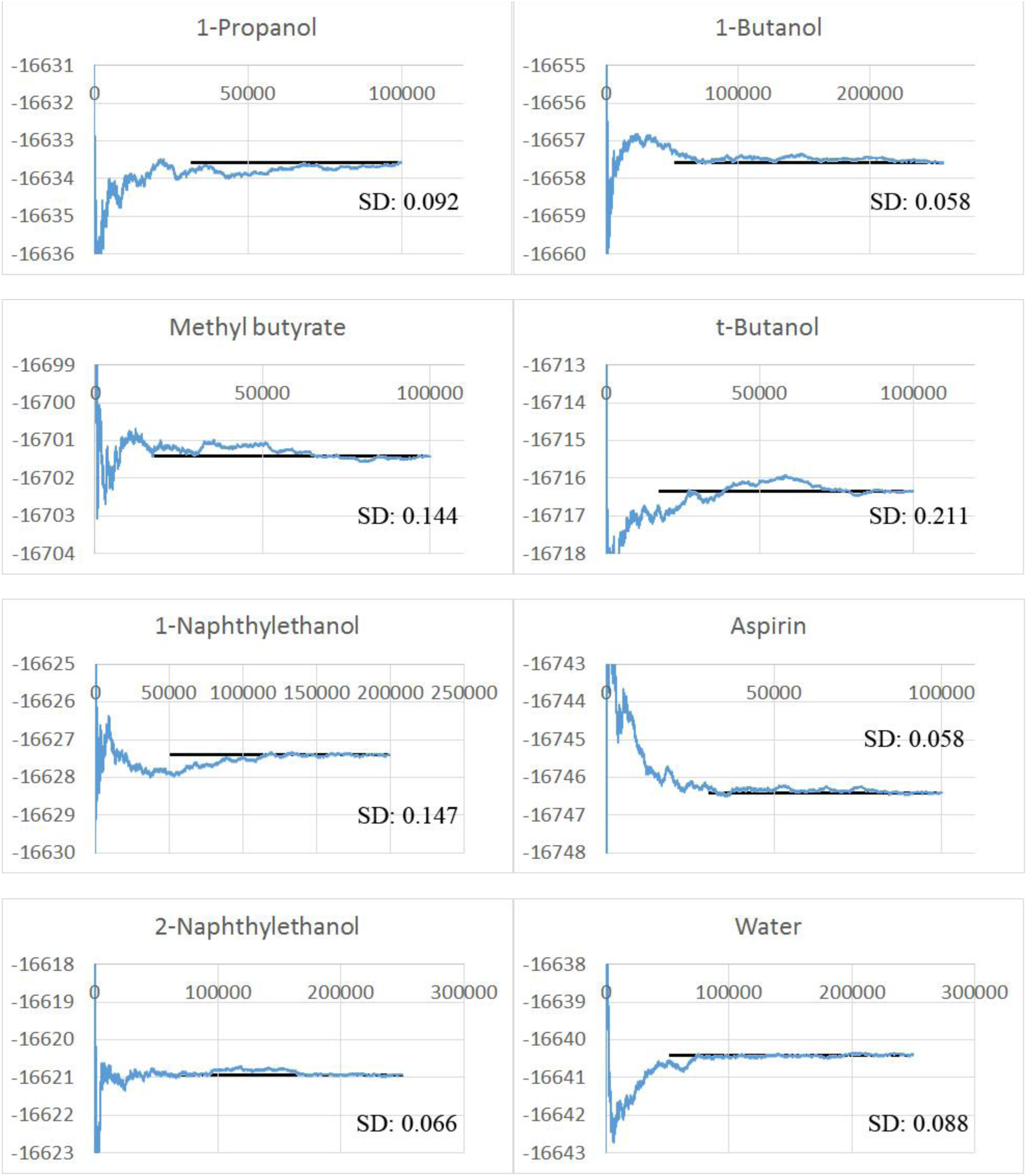
The enthalpy convergence of ligand and water trajectories. The convergence is represented by average potential energy (<E>) plotted against the number of frames. X-axes are numbers of frames used in the average. Y-axes are average potential energies in kcal/mol. The Y value of the black line indicates the value used in reported enthalpy. The X range of the black line indicates the region used to calculate standard deviation (SD) of the averaged potential energy, and the SD (kcal/mol) is labeled on the figures.

**Figure SI 9.**
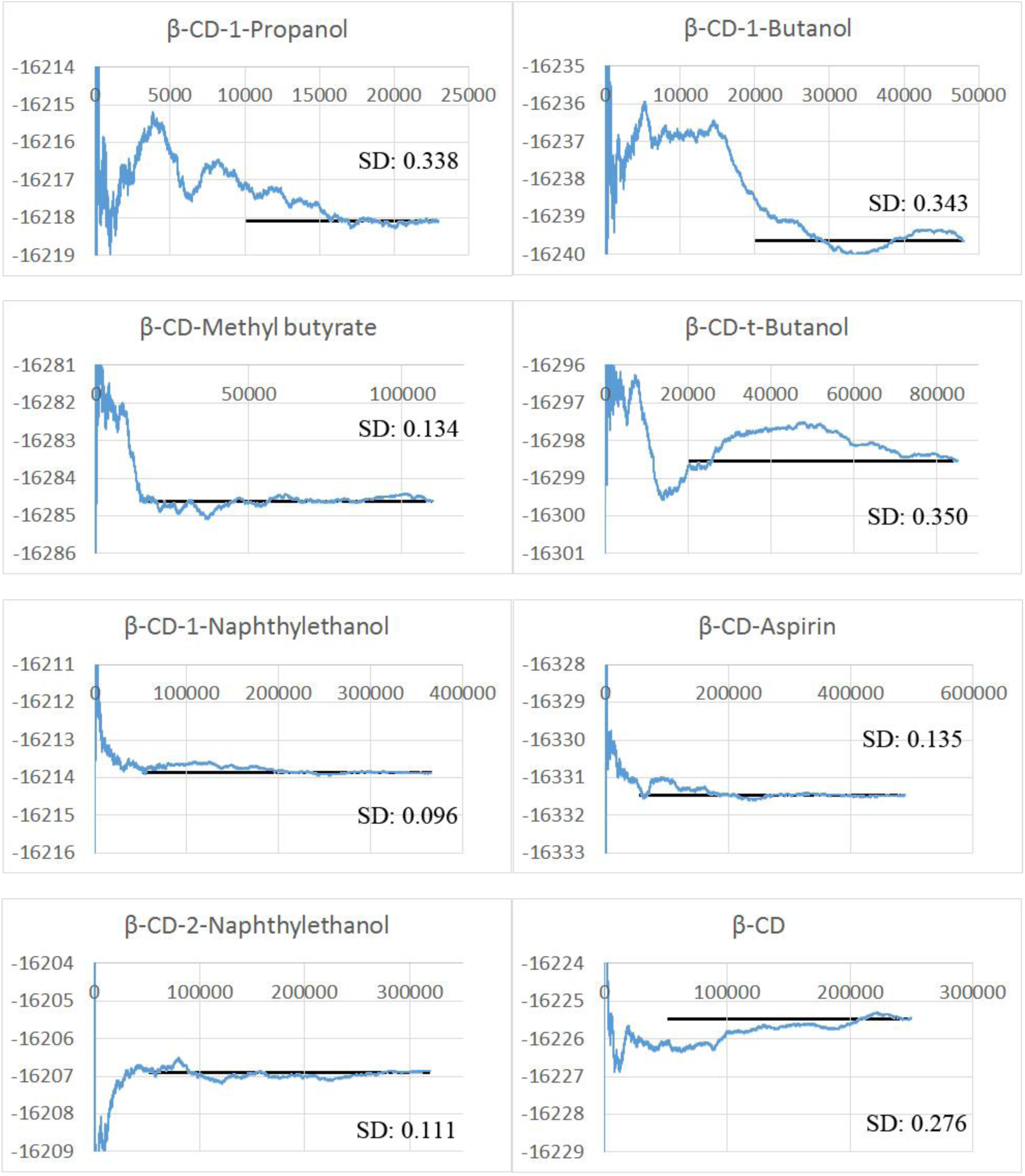
The enthalpy convergence of free β-CD and β-CD complexes using GAFF-CD force field. Refer to Figure SI 8 for the interpretation of the figures.

**Figure SI 10.**
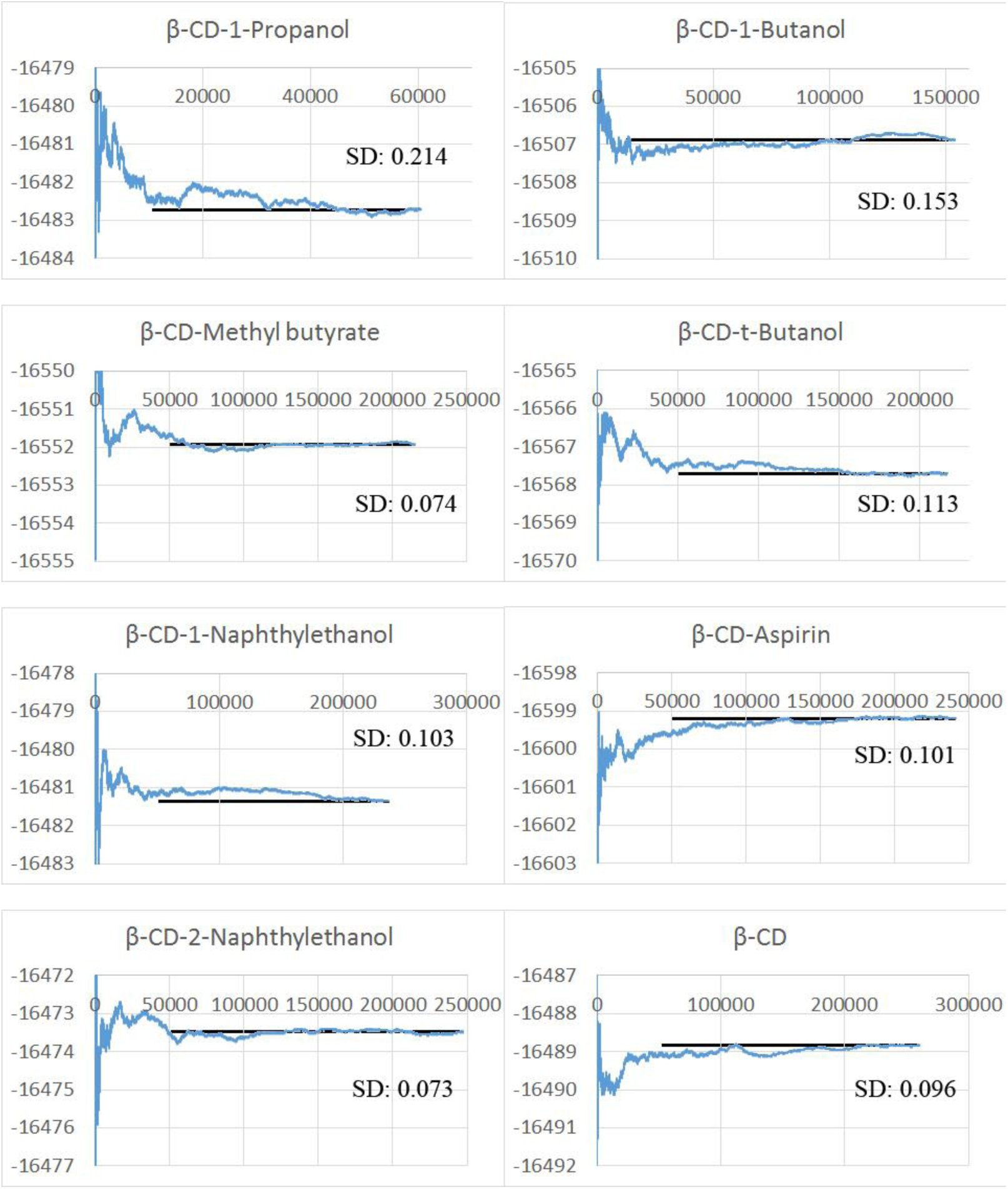
The enthalpy convergence of free β-CD and β-CD complexes using q4MD-CD force field. Refer to Figure SI 8 for the interpretation of the figures.

### 13. Movies of Association and Dissociation Pathways

In the movies, carbon atoms are in cyan and oxygen atoms are in red on β-CD. The ligands are rendered in green. Hydrogen atoms are omitted. Movies are listed as below.

From GAFF-CD force field:

1 β-CD-aspirin direct association: GAFF-CD-b-CD-aspirin_direct_ association.swf

2 β-CD-aspirin direct dissociation: GAFF-CD-b-CD-aspirin_direct_dissociation.swf

3 β-CD-aspirin sticky association: GAFF-CD-b-CD-aspirin_sticky_ association.swf

4 β-CD-aspirin sticky dissociation: GAFF-CD-b-CD-aspirin_ sticky_dissociation.swf

5 β-CD-t-butanol direct association: GAFF-CD-b-CD-t-butanol_direct_association.swf

6 β-CD-t-butanol direct dissociation: GAFF-CD-b-CD-t-butanol_direct_dissociation.swf

From q4MD-CD force field:

7 β-CD-aspirin direct association: q4MD-CD-b-CD-aspirin_direct_ association.swf

8 β-CD-aspirin direct dissociation: q4MD-CD-b-CD-aspirin_direct_dissociation.swf

9 β-CD-aspirin sticky association: q4MD-CD-b-CD-aspirin_sticky_ association.swf

10 β-CD-aspirin sticky dissociation: q4MD-CD-b-CD-aspirin_ sticky_dissociation.swf

11 β-CD-t-butanol direct association: q4MD-CD-b-CD-t-butanol_direct_association.swf

12 β-CD-t-butanol direct dissociation: q4MD-CD-b-CD-t-butanol_direct_dissociation.swf

### 14. Ligand External Degrees of Freedom

**Figure SI 11.**
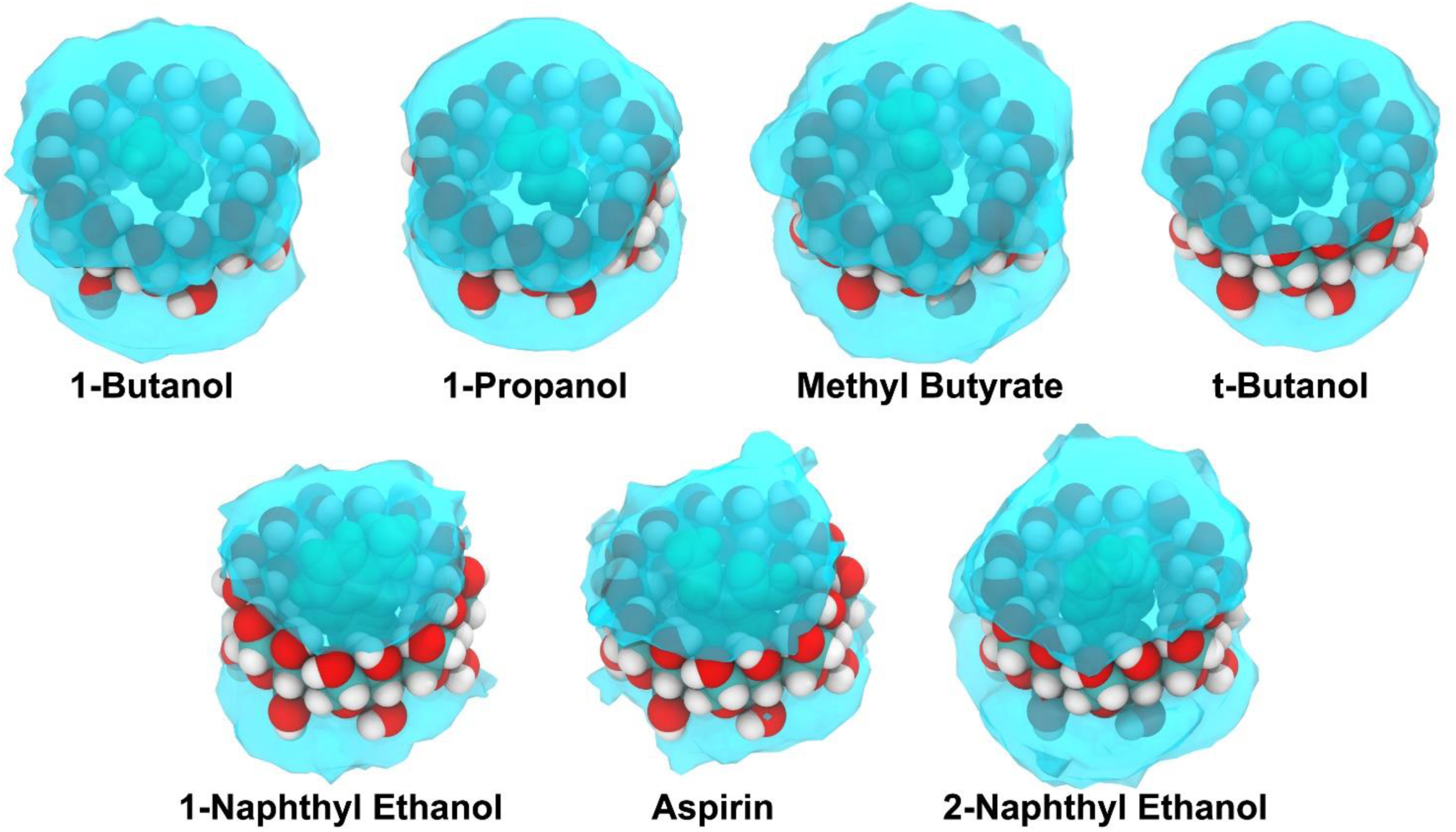
The binding poses of β-CD and ligand complex in GAFF-CD. For simplicity, the crystal structure of β-CD is shown in *VDW* representation. One representative conformation of ligands is rendered in green *VDW* representation. The volumes occupied by the bound state conformations are shown in cyan transparent *Surface* representation. The ligand conformations are obtained by superimposing β-CD coordinates in the trajectories to the crystal structure of β-CD. Similar binding volumes can be observed in q4MD-CD.

### 15. Bound and Free Period Lengths of the Complexes

**Table SI 10.** The bound and free period lengths of the complexes in GAFF-CD. All lengths of periods are in 10 ps.

|  | 1-Propanol |  | 1-Butanol |  | Methyl butyrate |  |  |  | t-Butanol |  |  |  |
| --- | --- | --- | --- | --- | --- | --- | --- | --- | --- | --- | --- | --- |
|  | Bound | Free | Bound | Free | Bound |  | Free |  | Bound |  | Free |  |
| 1 | 1801 | 5834 | 9769 | 899 | 5287 | 189 | 639 | 8354 | 8381 | 119 | 594 | 20517 |
| 2 | 167 | 1521 | 6455 | 8840 | 885 | 298 | 2516 | 2143 | 171 | 119 | 17392 | 7004 |
| 3 | 1090 | 482 | 353 | 77243 | 2931 | 111 | 1730 | 23327 | 418 | 704 | 28343 | 6095 |
| 4 | 731 | 241 | 105 | 33202 | 342 | 394 | 14352 | 41390 | 109 | 108 | 1257 | 1631 |
| 5 | 153 | 57826 | 218 | 2025 | 302 | 1305 | 2672 | 1141 | 179 | 6601 | 54931 | 8604 |
| 6 | 107 | 26676 | 104 | 10408 | 101 | 14187 | 11150 | 13848 | 181 | 347 | 5509 | 42418 |
| 7 | 102 | 31381 | 318 | 32669 | 203 | 103 | 13537 | 37917 | 104 | 364 | 345 | 1985 |
| 8 | 268 | 23494 | 129 | 1050 | 152 | 150 | 6029 | 9514 | 156 | 137 | 4972 | 220 |
| 9 | 199 | 2233 | 136 | 20238 | 153 | 152 | 34568 | 5714 | 105 | 153 | 8409 | 7350 |
| 10 | 184 | 17913 | 205 | 7329 | 252 | 17996 | 26941 | 4260 | 111 | 482 | 57093 | 512 |
| 11 | 741 | 13660 | 161 | 15286 | 198 | 506 | 13582 | 1550 | 103 | 193 | 20737 | 885 |
| 12 | 113 | 42695 | 156 | 9173 | 151 | 217 | 7508 | 6697 | 290 | 1669 | 25645 | 18557 |
| 13 | 145 | 28855 | 141 | 5079 | 109 | 183 | 722 | 348 | 118 | 146 | 2032 | 30213 |
| 14 | 103 | 4623 | 104 | 4886 | 205 | 102 | 163 | 16516 | 178 |  | 10998 |  |
| 15 | 100 | 16875 | 404 | 27418 | 1604 | 112 | 8441 | 32830 | 19253 |  | 3640 |  |
| 16 | 106 | 5690 | 737 | 27469 | 1044 | 153 | 1339 | 2681 | 164 |  | 4157 |  |
| 17 | 148 | 2049 | 320 | 10703 | 1296 | 449 | 1372 | 188 | 18665 |  | 18096 |  |
| 18 | 108 | 18646 | 132 | 4843 | 130 | 175 | 6512 | 128 | 141 |  | 241 |  |
| 19 | 142 | 23696 | 105 | 4345 | 1719 | 282 | 5651 | 1263 | 546 |  | 22927 |  |
| 20 | 226 | 26882 | 116 | 3582 | 111 | 1116 | 16376 | 3612 | 156 |  | 4234 |  |
| 21 | 239 | 128 | 182 | 1894 | 420 | 228 | 5834 | 6019 | 2234 |  | 4107 |  |
| 22 | 364 | 48217 | 121 | 38219 | 102 | 138 | 3798 | 3953 | 120 |  | 345 |  |
| 23 | 159 | 18383 | 264 | 1773 | 2658 | 103 | 5771 | 1967 | 114 |  | 5678 |  |
| 24 | 103 | 43352 | 201 | 24159 | 299 | 171 | 1467 | 119 | 130 |  | 8925 |  |
| 25 | 128 | 12708 | 315 | 2688 | 695 | 134 | 8344 | 20276 | 172 |  | 2267 |  |
| 26 | 401 | 13298 | 102 | 16344 | 465 | 189 | 18912 | 14028 | 128 |  | 5984 |  |
| 27 | 153 | 23600 | 142 | 38714 | 104 | 109 | 2265 | 6755 | 133 |  | 3860 |  |
| 28 | 201 | 787 | 110 | 1534 | 390 | 184 | 5454 | 2378 | 323 |  | 3144 |  |
| 29 | 108 | 720 | 178 | 894 | 4840 | 324 | 1586 | 1427 | 108 |  | 214 |  |
| 30 | 119 | 16617 | 130 | 1318 | 112 | 1490 | 11015 | 12289 | 431 |  | 12414 |  |
| 31 | 105 | 23846 | 115 | 4496 | 123 |  | 2417 |  | 152 |  | 2287 |  |
| 32 | 127 | 5159 | 282 | 1977 | 397 |  | 708 |  | 112 |  | 6350 |  |
| 33 | 111 | 10813 | 113 | 16631 | 126 |  | 1523 |  | 147 |  | 3421 |  |
| 34 | 162 | 1775 | 100 | 23787 | 238 |  | 1279 |  | 263 |  | 12449 |  |
| 35 | 253 | 37611 | 196 | 25623 | 151 |  | 387 |  | 477 |  | 2439 |  |
| 36 | 151 | 11333 | 7042 | 3528 | 196 |  | 16738 |  | 3728 |  | 3933 |  |
| 37 | 174 | 2753 | 418 | 6781 | 600 |  | 3703 |  | 176 |  | 5920 |  |
| 38 | 101 | 16619 | 5800 | 13087 | 224 |  | 21087 |  | 143 |  | 14056 |  |
| 39 | 123 | 993 | 461 | 1300 | 266 |  | 9822 |  | 177 |  | 17870 |  |
| 40 |  |  | 199 | 758 | 156 |  | 34084 |  | 117 |  | 8000 |  |
| 41 |  |  | 188 | 62738 | 152 |  | 4415 |  | 640 |  | 458 |  |
| 42 |  |  | 133 | 18110 | 100 |  | 5236 |  | 236 |  | 17375 |  |
| 43 |  |  |  |  | 9922 |  | 1564 |  | 233 |  | 9313 |  |
| 44 |  |  |  |  | 4159 |  | 635 |  | 108 |  | 21409 |  |
| 45 |  |  |  |  | 12086 |  | 3829 |  | 120 |  | 878 |  |
| 46 |  |  |  |  | 101 |  | 2604 |  | 101 |  | 46032 |  |
| 47 |  |  |  |  | 252 |  | 337 |  | 135 |  | 10473 |  |
| 48 |  |  |  |  | 826 |  | 13528 |  | 209 |  | 31839 |  |
| 49 |  |  |  |  | 195 |  | 4446 |  | 771 |  | 28408 |  |
| Avg. | 256.8 | 16409.8 | 880.0 | 14596.2 | 1250.4 |  | 8243.3 |  | 1171.1 |  | 11732.1 |  |
| # of Periods | 39 | 39 | 42 | 42 | 79 |  | 79 |  | 52 |  | 52 |  |

|  | 1-Naphthyl Ethanol |  | Aspirin |  |  |  |  |  | 2-Naphthyl Ethanol |  |
| --- | --- | --- | --- | --- | --- | --- | --- | --- | --- | --- |
|  | Bound | Free | Bound |  |  | Free |  |  | Bound | Free |
| 1 | 60007 | 2437 | 14310 | 419 | 228 | 1315 | 1955 | 8016 | 79434 | 1094 |
| 2 | 292 | 2171 | 3850 | 122 | 368 | 673 | 2497 | 2020 | 15270 | 2190 |
| 3 | 181 | 767 | 29400 | 2178 | 269 | 3586 | 3234 | 3849 | 4633 | 1380 |
| 4 | 6478 | 3866 | 407 | 169 | 275 | 3655 | 675 | 3969 | 411 | 1117 |
| 5 | 7108 | 4005 | 1002 | 255 | 115 | 148 | 10442 | 2624 | 156 | 4645 |
| 6 | 287 | 853 | 1042 | 120 | 146 | 3279 | 1057 | 2389 | 10619 | 534 |
| 7 | 487 | 3852 | 126 | 104 | 266 | 155 | 11040 | 487 | 609 | 338 |
| 8 | 146 | 11887 | 119 | 153 | 376 | 1959 | 3667 | 4619 | 154 | 227 |
| 9 | 293800 | 9182 | 10190 | 247 | 176 | 1752 | 1603 | 2014 | 1785 | 1130 |
| 10 | 206 | 4737 | 264 | 312 | 370 | 1539 | 586 | 1991 | 363 | 1250 |
| 11 | 512 | 8456 | 1271 | 115 | 199 | 2940 | 6580 | 1221 | 125 | 1230 |
| 12 | 259 | 344 | 1315 | 254 | 105 | 1103 | 825 | 5897 | 288 | 1452 |
| 13 | 599630 | 206 | 46697 | 130 | 3301 | 2840 | 870 | 6453 | 4976 | 10239 |
| 14 | 111 | 1850 | 2729 | 417 | 238 | 293 | 1002 | 4969 | 823 | 5208 |
| 15 | 705 | 2377 | 5417 | 17434 | 125 | 4436 | 3449 | 10530 | 223 | 1495 |
| 16 | 72801 |  | 232 | 5570 | 343 | 1375 | 1431 | 4637 | 432 | 1638 |
| 17 |  |  | 14907 | 848 | 4538 | 799 | 430 | 4674 | 150 | 1935 |
| 18 |  |  | 247 | 217 | 474 | 210 | 640 | 736 | 771 | 8741 |
| 19 |  |  | 849 | 119 | 478 | 650 | 12050 | 7552 | 35439 | 5824 |
| 20 |  |  | 8436 | 119 | 118 | 5095 | 8876 | 5736 | 125 | 3210 |
| 21 |  |  | 233 | 150 | 289 | 391 | 1931 | 3842 | 489 | 722 |
| 22 |  |  | 448 | 271 | 112 | 1437 | 1344 | 189 | 165 | 776 |
| 23 |  |  | 1096 | 114 | 9699 | 2252 | 1266 | 1835 | 717 | 7637 |
| 24 |  |  | 197 | 117 | 7116 | 3407 | 2658 | 680 | 153 | 4746 |
| 25 |  |  | 362 | 130 | 204 | 1072 | 2082 | 1488 | 192 | 1610 |
| 26 |  |  | 16445 | 109 | 764 | 3851 | 8874 | 1440 | 198 | 3272 |
| 27 |  |  | 276 | 218 | 178 | 1648 | 229 | 2132 | 8415 | 3760 |
| 28 |  |  | 124 | 152 | 274 | 2465 | 640 | 432 | 3003 | 4103 |
| 29 |  |  | 167 | 483 | 149 | 1415 | 16437 | 1535 | 180 | 13227 |
| 30 |  |  | 243 | 109 | 184 | 2052 | 1347 | 6401 | 138 | 1061 |
| 31 |  |  | 829 | 108 | 105 | 2467 | 2244 | 3062 | 101 | 15234 |
| 32 |  |  | 360 | 247 | 205 | 6322 | 6913 | 4841 | 193 | 3048 |
| 33 |  |  | 103 | 143 | 148 | 2129 | 2248 | 2453 | 147 | 2372 |
| 34 |  |  | 395 | 713 | 5682 | 2254 | 7830 | 347 | 191 | 2997 |
| 35 |  |  | 141 | 140 | 208 | 1480 | 251 | 1220 | 154 | 7530 |
| 36 |  |  | 172 | 175 |  | 1441 | 11777 |  | 20316 | 1138 |
| 37 |  |  | 387 | 111 |  | 11984 | 1962 |  | 8508 | 737 |
| 38 |  |  | 530 | 246 |  | 1669 | 331 |  | 5201 | 199 |
| 39 |  |  | 346 | 165 |  | 820 | 8042 |  | 12334 | 3273 |
| 40 |  |  | 2540 | 207 |  | 1290 | 339 |  | 188001 | 537 |
| 41 |  |  | 199 | 234 |  | 1169 | 1093 |  | 214790 | 1692 |
| 42 |  |  | 152 | 118 |  | 703 | 3593 |  | 1340 | 1333 |
| 43 |  |  | 206 | 122 |  | 4479 | 2950 |  | 3312 | 4452 |
| 44 |  |  | 37326 | 124 |  | 215 | 236 |  | 217 | 2548 |
| 45 |  |  | 2111 | 288 |  | 1714 | 3461 |  | 95640 | 8830 |
| 46 |  |  | 1585 | 164 |  | 549 | 3077 |  | 9632 | 7124 |
| 47 |  |  | 270594 | 155 |  | 203 | 239 |  | 250 | 1809 |
| 48 |  |  | 2878 | 318 |  | 1078 | 6615 |  | 277 | 3692 |
| 49 |  |  | 121 | 245 |  | 1708 | 2957 |  | 204624 |  |
| Avg. | 65188.1 | 3799.3 | 4183.3 |  |  | 2959.6 |  |  | 19095.2 | 3423.7 |
| # of Periods | 16 | 15 | 133 |  |  | 133 |  |  | 49 | 48 |

**Table SI 11.** The bound and free period lengths of the complexes in q4MD-CD. All lengths of periods are in 10 ps.

**Table SI 10.** The bound and free period lengths of the complexes in GAFF-CD. All lengths of periods are in 10 ps.
|  | 1-Propanol |  |  |  |  |  | 1-Butanol |  |  |  | Methyl Butyrate |  | t-Butanol |  |
| --- | --- | --- | --- | --- | --- | --- | --- | --- | --- | --- | --- | --- | --- | --- |
|  | Bound |  |  | Free |  |  | Bound |  | Free |  | Bound | Free | Bound | Free |
| 1 | 1066 | 101 | 1337 | 927 | 3992 | 3705 | 1669 | 311 | 5868 | 2015 | 4075 | 254 | 2079 | 4536 |
| 2 | 726 | 751 | 155 | 1994 | 3029 | 356 | 4436 | 1216 | 2529 | 1686 | 7411 | 560 | 200 | 2727 |
| 3 | 110 | 2499 | 247 | 3265 | 157 | 1527 | 3418 | 1606 | 691 | 1731 | 2199 | 1595 | 4888 | 2110 |
| 4 | 388 | 1001 | 815 | 9801 | 107 | 1917 | 675 | 4533 | 4797 | 415 | 4978 | 715 | 1518 | 581 |
| 5 | 909 | 463 | 4468 | 1944 | 1384 | 7095 | 839 | 1397 | 403 | 610 | 7610 | 10426 | 40660 | 3023 |
| 6 | 211 | 1323 | 581 | 5500 | 5008 | 3284 | 3525 | 219 | 4471 | 4280 | 7614 | 1374 | 17924 | 996 |
| 7 | 328 | 105 | 476 | 10021 | 292 | 2142 | 3339 | 721 | 1554 | 3039 | 2318 | 3102 | 19064 | 4732 |
| 8 | 642 | 1930 | 538 | 1772 | 4243 | 1017 | 1718 | 4044 | 1887 | 803 | 59426 | 1601 | 225 | 2162 |
| 9 | 3051 | 104 | 404 | 798 | 357 | 268 | 8429 | 2363 | 1076 | 1430 | 1513 | 2075 | 42030 | 5703 |
| 10 | 321 | 1181 | 455 | 4932 | 3457 | 4589 | 7477 | 2014 | 1874 | 2083 | 19404 | 1543 | 162 | 1199 |
| 11 | 823 | 1104 | 366 | 962 | 3717 | 4783 | 1597 | 1987 | 4394 | 898 | 21044 | 1334 | 109 | 1899 |
| 12 | 1468 | 613 | 2245 | 1553 | 2755 | 3235 | 123 | 12994 | 793 | 4889 | 22974 | 765 | 12323 | 1948 |
| 13 | 644 | 554 | 111 | 2321 | 10977 | 2270 | 130 | 3018 | 337 | 1162 | 6295 | 852 | 84946 | 8435 |
| 14 | 418 | 1459 | 295 | 986 | 3376 | 545 | 3550 | 2099 | 1987 | 101 | 5330 | 2041 | 11078 | 728 |
| 15 | 749 | 104 | 216 | 488 | 6438 | 2996 | 8262 | 4062 | 1588 | 428 | 9781 | 1596 | 9311 | 4121 |
| 16 | 1158 | 728 | 105 | 258 | 794 | 2695 | 12898 | 6707 | 220 | 2743 | 45096 | 3450 | 1798 | 5267 |
| 17 | 271 | 642 | 1766 | 1840 | 100 | 5162 | 5868 | 1271 | 711 | 171 | 16006 | 569 | 5173 | 332 |
| 18 | 411 | 156 | 2064 | 147 | 8925 | 5071 | 276 | 1564 | 3439 | 7825 | 1751 | 5507 | 7726 | 6462 |
| 19 | 2034 | 493 | 151 | 5741 | 1848 | 4353 | 375 | 4661 | 2854 | 759 | 15608 | 701 | 1875 | 3918 |
| 20 | 1274 | 1126 | 257 | 8362 | 2226 | 660 | 1082 | 553 | 2271 | 219 | 796 | 1086 | 2130 | 5128 |
| 21 | 227 | 1512 | 589 | 4757 | 2646 | 686 | 1014 | 679 | 810 | 3644 | 7949 | 1607 | 27334 | 1724 |
| 22 | 1112 | 1126 | 490 | 416 | 710 | 4381 | 211 | 7893 | 1368 | 321 | 22236 | 1400 | 16820 | 1317 |
| 23 | 1830 | 440 | 340 | 1205 | 1098 | 597 | 6426 | 911 | 1790 | 175 | 2675 | 1938 | 32273 | 2846 |
| 24 | 1121 | 571 | 633 | 826 | 2727 | 634 | 3235 | 2964 | 1530 | 1253 | 10181 | 487 | 22784 | 1384 |
| 25 | 921 | 1376 | 1172 | 4051 | 1468 | 1811 | 1562 | 371 | 4497 | 2381 | 2960 | 3447 | 141 | 458 |
| 26 | 2138 | 1502 | 154 | 4796 | 3993 | 1605 | 10609 | 156 | 2045 | 2131 | 5905 | 2755 | 558 | 839 |
| 27 | 634 | 413 | 323 | 1928 | 1560 | 768 | 670 | 175 | 990 | 4592 | 12412 | 1043 | 10296 |  |
| 28 | 118 | 127 | 1134 | 1078 | 6956 | 687 | 5644 | 480 | 2284 | 2993 | 1748 | 370 |  |  |
| 29 | 519 | 884 | 793 | 1290 | 1075 | 1204 | 4651 | 663 | 346 | 422 | 4834 | 4757 |  |  |
| 30 | 1167 | 194 | 995 | 566 | 1807 | 519 | 240 | 8417 | 1629 | 1100 | 5901 | 262 |  |  |
| 31 | 688 | 1792 |  | 575 | 3608 |  | 1358 | 6132 | 1072 | 1528 | 5626 | 466 |  |  |
| 32 | 460 | 1625 |  | 1589 | 453 |  | 2035 | 111 | 5150 | 4204 | 1066 | 1872 |  |  |
| 33 | 365 | 900 |  | 2779 | 6987 |  | 1284 | 416 | 7472 | 701 | 134 | 3888 |  |  |
| 34 | 666 | 257 |  | 3514 | 1546 |  | 9153 | 4337 | 479 | 2745 | 4463 | 3745 |  |  |
| 35 | 108 | 129 |  | 5846 | 7895 |  | 161 | 1744 | 317 | 1528 | 117 | 4174 |  |  |
| 36 | 2889 | 183 |  | 5320 | 845 |  | 4359 | 2431 | 844 | 6045 | 2955 | 1884 |  |  |
| 37 | 2551 | 379 |  | 196 | 5664 |  | 6786 | 440 | 2917 | 957 | 8285 | 1671 |  |  |
| 38 | 510 | 900 |  | 1777 | 4104 |  | 8748 | 3703 | 2521 | 595 | 196 | 3873 |  |  |
| 39 | 509 | 910 |  | 926 | 844 |  | 5470 | 4059 | 3535 | 6623 | 5821 | 1554 |  |  |
| 40 | 909 | 115 |  | 2147 | 3121 |  | 208 | 2320 | 861 | 4446 |  | 968 |  |  |
| 41 | 513 | 260 |  | 3317 | 6226 |  | 3263 |  | 485 |  |  |  |  |  |
| 42 | 2092 | 600 |  | 608 | 1280 |  | 1238 |  | 2300 |  |  |  |  |  |
| 43 | 251 | 715 |  | 6986 | 877 |  | 1517 |  | 4515 |  |  |  |  |  |
| 44 | 199 | 664 |  | 209 | 1552 |  | 629 |  | 1418 |  |  |  |  |  |
| 45 | 2411 | 604 |  | 955 | 3575 |  | 2902 |  | 720 |  |  |  |  |  |
| 46 | 813 | 238 |  | 261 | 140 |  | 2728 |  | 1749 |  |  |  |  |  |
| 47 | 777 | 1167 |  | 3399 | 5357 |  | 667 |  | 3268 |  |  |  |  |  |
| 48 | 494 | 3235 |  | 4399 | 1890 |  | 1279 |  | 198 |  |  |  |  |  |
| Avg. | 848.6 |  |  | 2722.8 |  |  | 2994.0 |  | 2119.6 |  | 9402.4 | 2082.7 | 13904.6 | 2868.3 |
| # of Periods | 128 |  |  | 128 |  |  | 89 |  | 89 |  | 39 | 40 | 27 | 26 |

|  | 1-Naphthyl Ethanol |  | Aspirin |  | 2-Naphthyl Ethanol |  |
| --- | --- | --- | --- | --- | --- | --- |
|  | Bound | Free | Bound | Free | Bound | Free |
| 1 | 19239 | 890 | 29988 | 737 | 388957 | 1465 |
| 2 | 138 | 2805 | 3198 | 593 | 103585 | 203 |
| 3 | 77015 | 5636 | 9970 | 2614 | 105790 |  |
| 4 | 167 | 1597 | 11443 | 400 |  |  |
| 5 | 280005 | 996 | 94979 | 1252 |  |  |
| 6 | 88794 | 4855 | 13411 | 436 |  |  |
| 7 | 12888 | 2118 | 44896 | 672 |  |  |
| 8 | 102857 |  | 46931 | 521 |  |  |
| 9 |  |  | 4055 | 320 |  |  |
| 10 |  |  | 112077 | 1570 |  |  |
| 11 |  |  | 626 | 661 |  |  |
| 12 |  |  | 31966 | 1974 |  |  |
| 13 |  |  | 40488 | 1767 |  |  |
| 14 |  |  | 64727 | 535 |  |  |
| 15 |  |  | 14034 | 1247 |  |  |
| 16 |  |  | 1007 | 1287 |  |  |
| 17 |  |  | 426 | 717 |  |  |
| 18 |  |  | 58475 |  |  |  |
| Avg. | 17360.1 | 1184.2 | 22624.7 | 1831.1 | 23275.2 | 949.3 |
| # of Periods | 8 | 7 | 18 | 17 | 3 | 2 |

### 16. Comparison of ΔG_Comp1_ and ΔG_Comp2_

**Figure SI 12.**
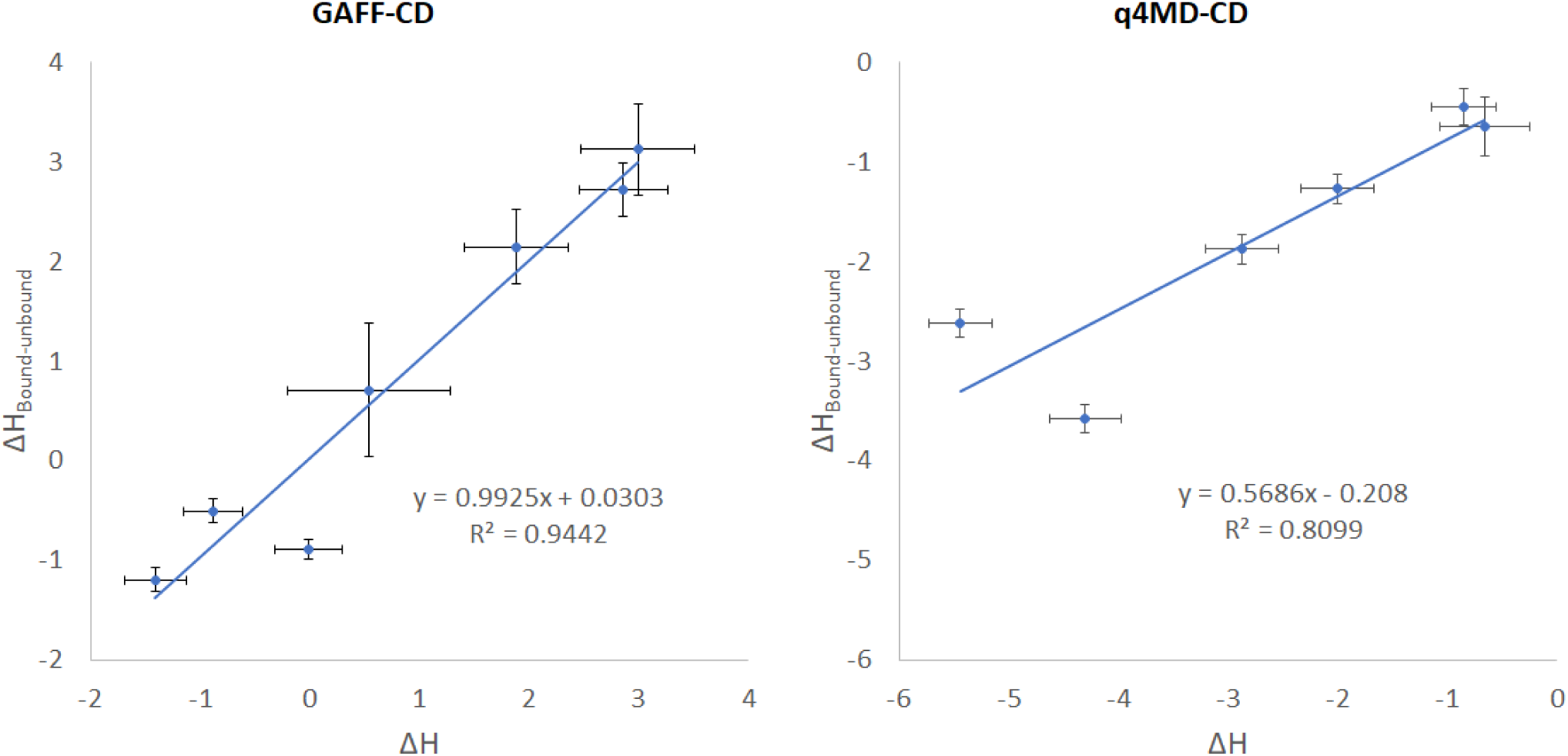
The correlations between ΔH, ΔH_Bound-unbound_ and experimental values. ΔH_Bound-unbound_ was computed by using average potential energy of bound and unbound conformations in the trajectories of the complexes. The correlations are label correspondingly. Error bars indicate the uncertainties.

